# Mechanistic origin of cell-size control and homeostasis in bacteria

**DOI:** 10.1101/478818

**Authors:** Fangwei Si, Guillaume Le Treut, John T. Sauls, Stephen Vadia, Petra Anne Levin, Suckjoon Jun

## Abstract

Evolutionarily divergent bacteria share a common phenomenological strategy for cell-size homeostasis under steady-state conditions. In the presence of inherent physiological stochasticity, cells following this “adder” principle gradually return to their steady-state size by adding a constant volume between birth and division regardless of their size at birth. However, the mechanism of the adder has been unknown despite intense efforts. In this work, we show that the adder is a direct consequence of two general processes in biology: (1) threshold -- accumulation of initiators and precursors required for cell division to a respective fixed number, and (2) balanced biosynthesis -- maintenance of their production proportional to volume growth. This mechanism is naturally robust to static growth inhibition, but also allows us to “reprogram” cell-size homeostasis in a quantitatively predictive manner in both Gram-negative *Escherichia coli* and Gram-positive *Bacillus subtilis*. By generating dynamic oscillations in the concentration of the division protein FtsZ, we were able to oscillate cell size at division and systematically break the adder. In contrast, periodic induction of replication initiator protein DnaA caused oscillations in cell size at initiation, but did not alter division size or the adder. Finally, we were able to restore the adder phenotype in slow-growing *E. coli*, the only known steady-state growth condition wherein *E. coli* significantly deviates from the adder, by repressing active degradation of division proteins. Together these results show that cell division and replication initiation are independently controlled at the gene-expression level, and that division processes exclusively drive cell-size homeostasis in bacteria.

**HIGHLIGHTS:**

- The adder requires accumulation of division proteins to a threshold for division.
- The adder requires constant production of division proteins during cell elongation.
- In *E. coli* and *B. subtilis*, initiation and division are independently controlled.
- In *E. coli* and *B. subtilis*, cell division exclusively drives size homeostasis.

**GRAPHICAL ABSTRACT:** 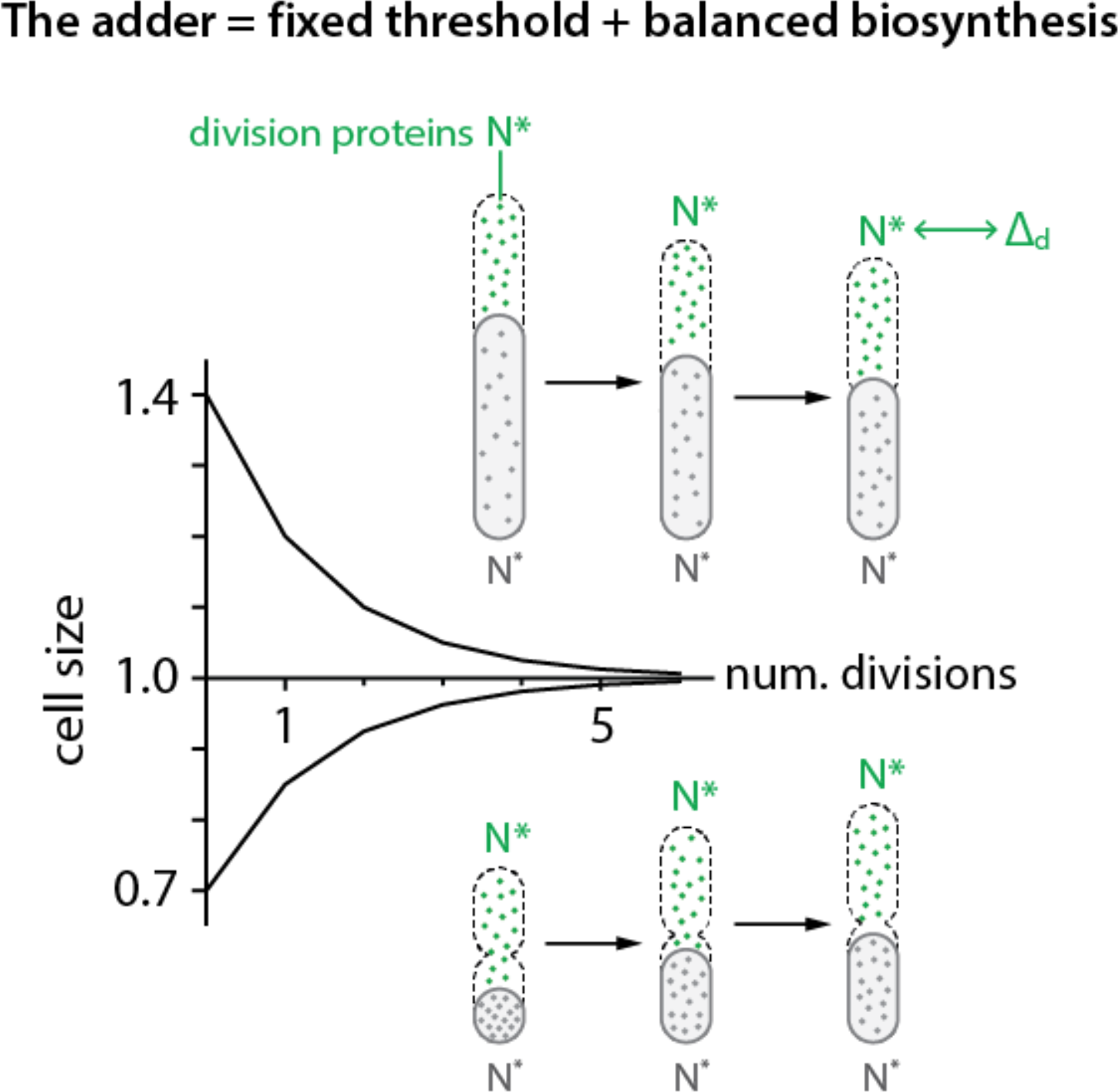

**eTOC Blurb:** Si and Le Treut *et al*. show that cell-size homeostasis in bacteria is exclusively driven by accumulation of division proteins to a threshold and their balanced biosynthesis during cell elongation. This mechanistic insight allowed them to reprogram cell-size homeostasis in both *E. coli* and *B. subtilis*. Evolutionary implications are discussed.

## INTRODUCTION

Cellular physiology is composed of inherently stochastic processes [1]. Cell size at birth can fluctuate due to asymmetric division events or alterations in the timing or speed of constriction. Without homeostatic control, cell size in a continuous lineage would diverge with each division cycle. Evolutionarily divergent organisms ensure size homeostasis at the single-cell level by following a phenomenological principle known as the “adder” [2–14]. A central property of the adder is that newborn cells deviating from the average size at birth add a nearly fixed volume between birth and division, allowing them to exponentially converge to the population average in each division cycle (Fig. 1). The adder sharply contrasts with a “sizer,” in which cells divide when they reach a fixed size. The adder principle has been extended to eukaryotes from yeast [9, 10, 13] to mammalian cells [11, 12] that have long been considered as sizers employing cell-cycle checkpoints.

**Figure 1:**
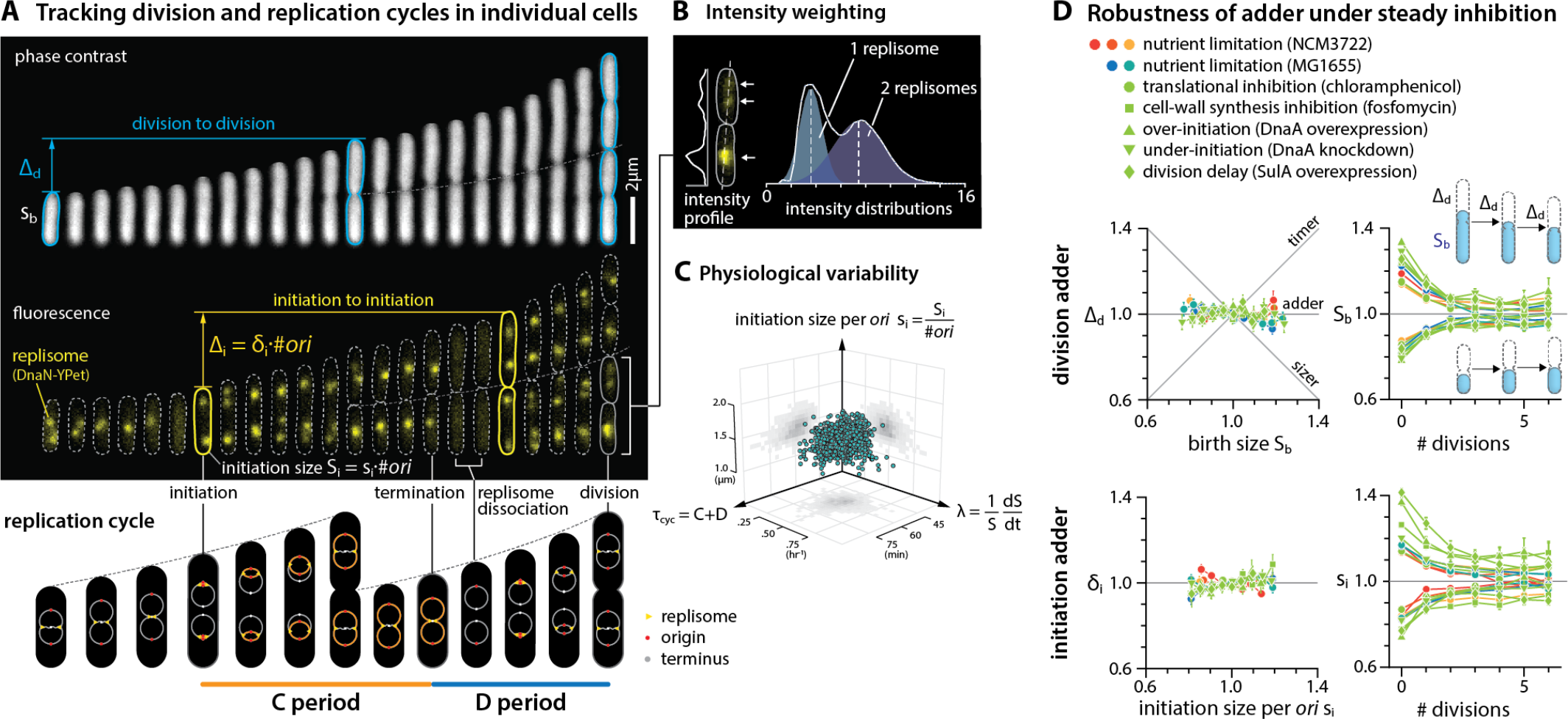
*E. coli* follows both the initiation adder and division adder, and both are robust to static inhibition of growth and biosynthesis. **(A)** Division adder vs. initiation adder. Upper: ∆_d_ is the added size between birth size S_b_ and division size S_d_, and ∆_i_ is the added size between total cell size at two consecutive initiations. Initiation size per *ori* s_i_ is defined as the initiation size S_i_ per number of replication origins, and δ_i_ represents ∆_i_ per origin. Cell length is used as cell size because cell width is mostly constant [3] (Figs. S3 and S4). Lower: Illustration of the replication cycle with two overlapping cell cycles. The growth condition is MOPS minimal medium with 0.2 % glucose. **(B)** Resolving overlapping foci using intensity weighting. The two peaks of intensity distribution of fluorescence foci indicate 1 and 2 replisomes, respectively (STAR Methods). **(C)** Three major measured physiological parameters, growth rate λ, τ_cyc_, and s_i_ all show variability and cross-correlation between 8%-20%. Each dot represents the data measured from one division cycle of a single cell. **(D)** Under static biosynthetic inhibition, *E. coli* robustly corrects deviations in initiation size per *ori* and division size following initiation adder and division adder, respectively. Each dot is the binned data and error bar indicates standard error of mean, same for other figures. In the correlation plots, the variables are normalized by their means. 6 μM of chloramphenicol and 0.05 μg/ml of fosfomycin were used. See sample size in Table S5.

The identification of the “adder” represented a major shift in our understanding of cell-size homeostasis [15, 16]. Naturally, many models have been proposed to explain the mechanistic origin of the adder phenotype. Most of these models can be classified into different groups by each model’s proposed implementation point of size control on the cell cycle. For example, recent works have suggested that the adder is governed by a replication-initiation-centric mechanism and division timing is determined by initiation in individual cells [17]. These models are based on the observation that cell size at initiation of DNA replication is invariant [18] at both single-cell [19] and population level [20]. These models are in contrast to a division-centric view of size homeostasis proposed earlier based on computer simulations [2] or biological constraints imposed on cellular resource allocation to division proteins [3, 21, 22]. Theoretical combination of replication and division controls has also been suggested at the phenomenological level [23, 24]. Alternatively, cell shape, or more specifically the surface-to-volume ratio of the cell, has also been suggested as the determining factor for size control [25].

In this work, we explain the mechanistic origin of cell-size homeostasis common to *E. coli* and *B. subtilis*, bacteria that diverged over a billion years ago. Specifically, we show that the adder phenotype is a direct consequence of two general processes in biology: (i) [threshold] accumulation of division initiators and precursors to a fixed threshold number per cell; and (ii) [balanced biosynthesis] their production is proportional to the growth of cell volume under steady-state condition. This mechanism allows us not only to “break” but also to “restore” the adder phenotype in a predictive manner under all major growth conditions.

Before proceeding to our results, we want to clarify the terminology. We use the term “cell-size control” for how cells determine their absolute size, and “cell-size homeostasis” for how cells correct deviations in size under steady-state growth. The two concepts are therefore closely related, yet differ with regard to whether emphasis is given to the requirement for threshold (for size control) or for balanced biosynthesis (for size homeostasis).

## RESULTS AND DISCUSSION

### Tracking replication and division cycles at the single-cell level

To illuminate the mechanisms underlying the adder principle, we performed a series of single-cell growth and cell-cycle tracking experiments under various growth conditions. We used a functional fluorescently-labeled replisome protein (DnaN-YPet) to image replication cycles, and a microfluidic mother machine to follow continuous lineages during steady-state growth [3, 26] (Fig. 1A, Fig. S1; STAR Methods).

A major technical challenge arises in studying replication dynamics when two replisome foci spatially overlap, which makes it difficult to analyze overlapping replication cycles. To resolve this issue, we tracked multiple replication forks from initiation to termination by extending previous imaging methods [8, 19, 27, 28] using the “intensity weighting” techniques [29, 30] developed for super-resolution microscopy. This method allowed us to resolve overlapping replisome foci based on the number of peaks measured in the intensity distribution (Fig. 1B; see STAR Methods). These measurements showed 8%-20% of coefficients of variation (CV) for physiological parameters consistent with previous measurements, with the CV of cell size at initiation exhibiting one of the narrowest distributions (CV=8%) (Figs. 1C and 2A; see also Supplemental Theory I).

**Figure 2:**
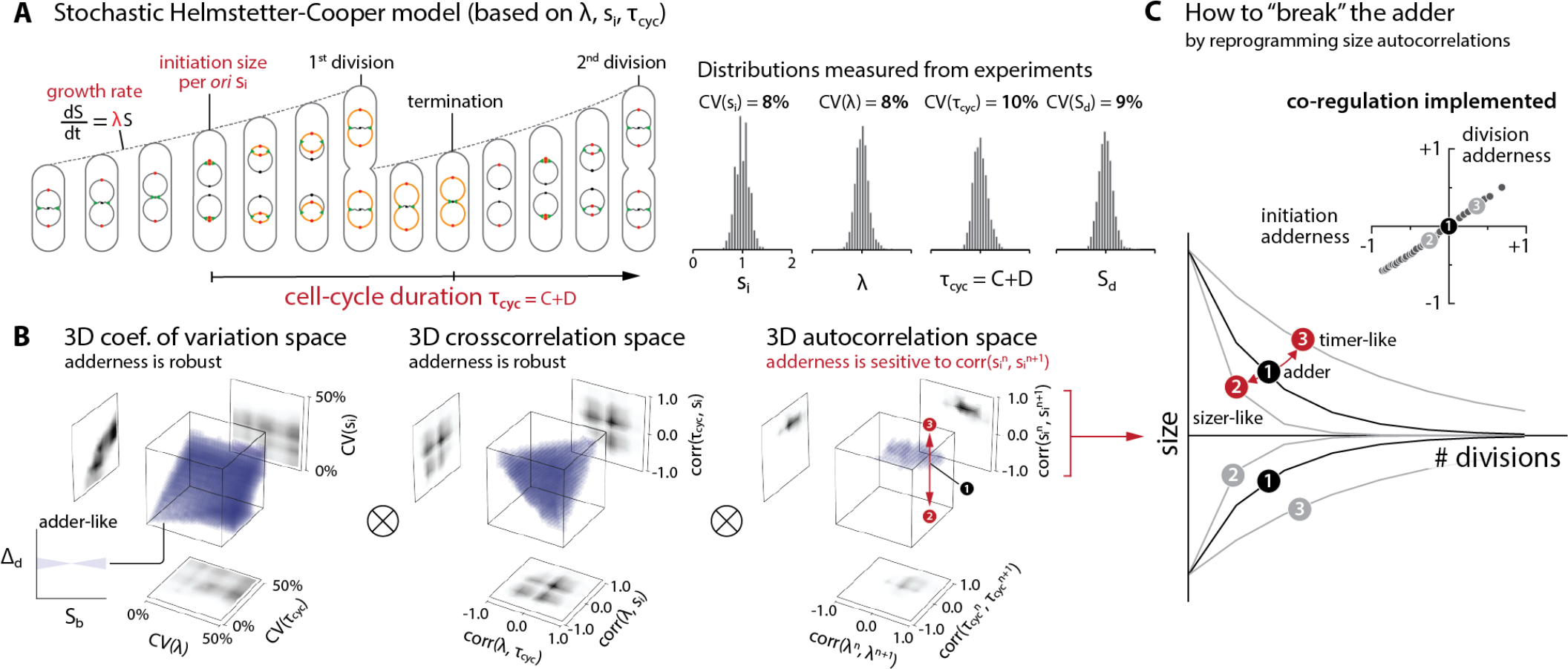
Survey of the 9-dimensional cell-size homeostasis space via stochastic Helmstetter-Cooper model assuming a co-regulation hypothesis between replication initiation and cell division. **(A)** The schematics of single-cell simulation. The simulation of cell growth and cell cycle progression took into account the stochasticity of three major physiological variables λ, τ_cyc_, and s_i_ measured from experiments. (STAR Methods; Supplemental Theory I). We did not consider variability in the septum position because it is the least variable physiological parameter in *E. coli* [3]. **(B)** The variability of division adderness in the 9-dimensional space surveyed by simulations. Pearson coefficient was used to quantify both cross-correlations (e.g. corr(λ, τ_cyc_)) and mother-daughter autocorrelations (e.g. corr(λ_n_, λ_n+1_)). Each 3-D plot is based on 1,000 simulations, and each simulation computed 10,000 division cycles (Supplemental Theory I). Purple color indicates an adder-like behavior defined as −0.1 < corr(∆_d_, S_b_) < 0.1 (see inset on bottom left). ⊗ means the actual simulation took the convolution of all nine dimensions. **(C)** Simulations revealed the unique role of autocorrelation of initiation size per *ori* in determining division adderness. The division adderness corr(∆_d_, S_b_) is monotonically dependent on the initiation adderness corr(δ_i_, s_i_) when varying the autocorrelation of initiation size per *ori* corr(s_n_, s_n+1_).

### *E. coli* follows an ‘initiation adder’ and a ‘division adder’, both robust to static inhibition of biosynthesis

Observation of wild type cells growing at steady state indicated the presence of two types of adder in *E. coli*: one functioning at division (hereafter a “division adder”) and the other at replication initiation (an “initiation adder”) (Fig. 1). Parallel to the “division adder” [2, 3], the “initiation adder” is characterized by the addition of a nearly constant size per origin between consecutive replication cycles. This ensures that deviations in cell size (per *ori*) at initiation exponentially converge to the population average in each replication cycle [8, 31] (Fig. 1D).

We next wanted to clarify the contribution of initiation and division to their respective adders. We utilized either tunable CRISPR interference [20, 32] to inhibit expression of *dnaA* that encodes the major bacterial DNA replication initiation protein, or an inducible-repressive promoter to modulate expression of a division inhibitor protein SulA. As expected, delays in replication and division both increased the average cell size (Fig. S2B). However, neither perturbation had a detectable effect on the initiation adder or division adder (Fig. 1D).

We also tested whether perturbations to global biosynthesis affect cell-size homeostasis, as they cause *E. coli* to deviate from the “growth law” of cell size, namely the well-established exponential relationship between the average cell size and the nutrient-imposed growth rate [20, 33]. In addition, previous work proposed accumulation of a fixed amount of cell-wall precursors as the mechanism of division adder [25]. We thus used either chloramphenicol or fosfomycin to target ribosomes or synthesis of cell-wall precursors, respectively, with the expectation that cells treated with these antibiotics would no longer exhibit the adder phenotype. In both cases, however, we found that defects in these major biosynthetic pathways did not affect either type of adder (Fig. 1D).

Together these data show that *E. coli* possess the capacity to buffer steady inhibitions of cell cycle progression or general biosynthesis to maintain robust size homeostasis.

### Using stochastic simulations to identify experimental conditions whereby *E. coli* should deviate from the adder

The robustness of adder posed unforeseen challenges for our attempts to identify the biological processes underlying the adder phenotype. Although we considered other types of perturbations or genetic screens, we realized that the physiological space was unrealistically large for brute-force search via single-cell time-lapse experiments. To circumvent the experimental challenges, we resorted to single-cell stochastic simulations and surveyed the entire physiological landscape (Fig. 2A).

A subtle but important problem in our initial stochastic simulations was how to decide the timing of cell division. This issue is related to an outstanding question in bacterial physiology: whether replication and division are independently controlled or co-regulated [34]. We implemented the Helmstetter-Cooper model [35, 36] that is often interpreted to mean that initiation triggers division after a fixed elapsed time *τ*_cyc_ = *C* + *D* (Figs. 1A and 2A) [17, 37]. To take into account biological stochasticity (Figs. 1C and 2A, Fig. S2A), we allowed for fluctuations in the three physiological variables, three cross-correlations, and three mother/daughter correlations (see STAR Methods and Supplemental Theory I). When we incorporated this implicit co-regulation hypothesis and stochasticity [17] in our simulations, we observed that the initiation adder leads to the division adder (Figs. 2B and 2C).

A brute-force numerical investigation of the entire parameter space suggested conditions under which size homeostasis should deviate from the adder and, importantly, an experimental means to “break” the adder. Specifically, when we varied the mother-daughter autocorrelation of the initiation size per *ori s*_i_ away from 0.5, cell-size homeostasis significantly deviated from the division adder. Otherwise, most other perturbations to physiological parameters did not severely affect the adder at division, reinforcing the general robustness of adder observed in our inhibition experiments (Fig. 1D). In fact, in the stochastic Helmstetter-Cooper model, the mother-daughter autocorrelation of *s*_i_ alone completely determines the initiation adderness (Fig. 2C; Supplemental Theory I). Since autocorrelation 0.5 is equivalent to exponential convergence of size deviations, we realized that the adder would break if we can experimentally modulate the speed of convergence [38].

### Dynamic perturbation of replication initiator synthesis to the synthesis of replication initiators breaks the initiation adder

To experimentally test our predictions from simulations, we sought to alter the autocorrelation of cell size at initiation to modify cell size homeostasis. We found the properties of the DNA replication protein DnaA made it ideal for our test. DnaA is a widely conserved essential protein required for initiation of DNA replication in bacteria. In bacteria in which it has been examined, replication initiation depends in part on accumulation of a sufficient number of DnaA molecules at the origin of replication [39–41]. Previous studies and our data showed that an underexpression of *dnaA* causes an initiation delay, whereas an overexpression of *dnaA* causes premature initiation (Fig. S3A) [20, 42].

The relationship between *dnaA* expression level and initiation size led us to a relatively simple strategy to break the adder. If we periodically induce *dnaA*, the initiation size would oscillate at the same frequency as the induction. This should introduce negative autocorrelations to both initiation size and division size as illustrated in Fig. 3A. The negative autocorrelations would be maximal when the period of oscillation *T* is two times the doubling time *τ*, because small-born cells add a larger size until division, whereas large-born cells add a smaller size until division, at every other division cycle (Fig. 3A). Consequently, cell-size homeostasis during oscillations is sizer-like.

**Figure 3:**
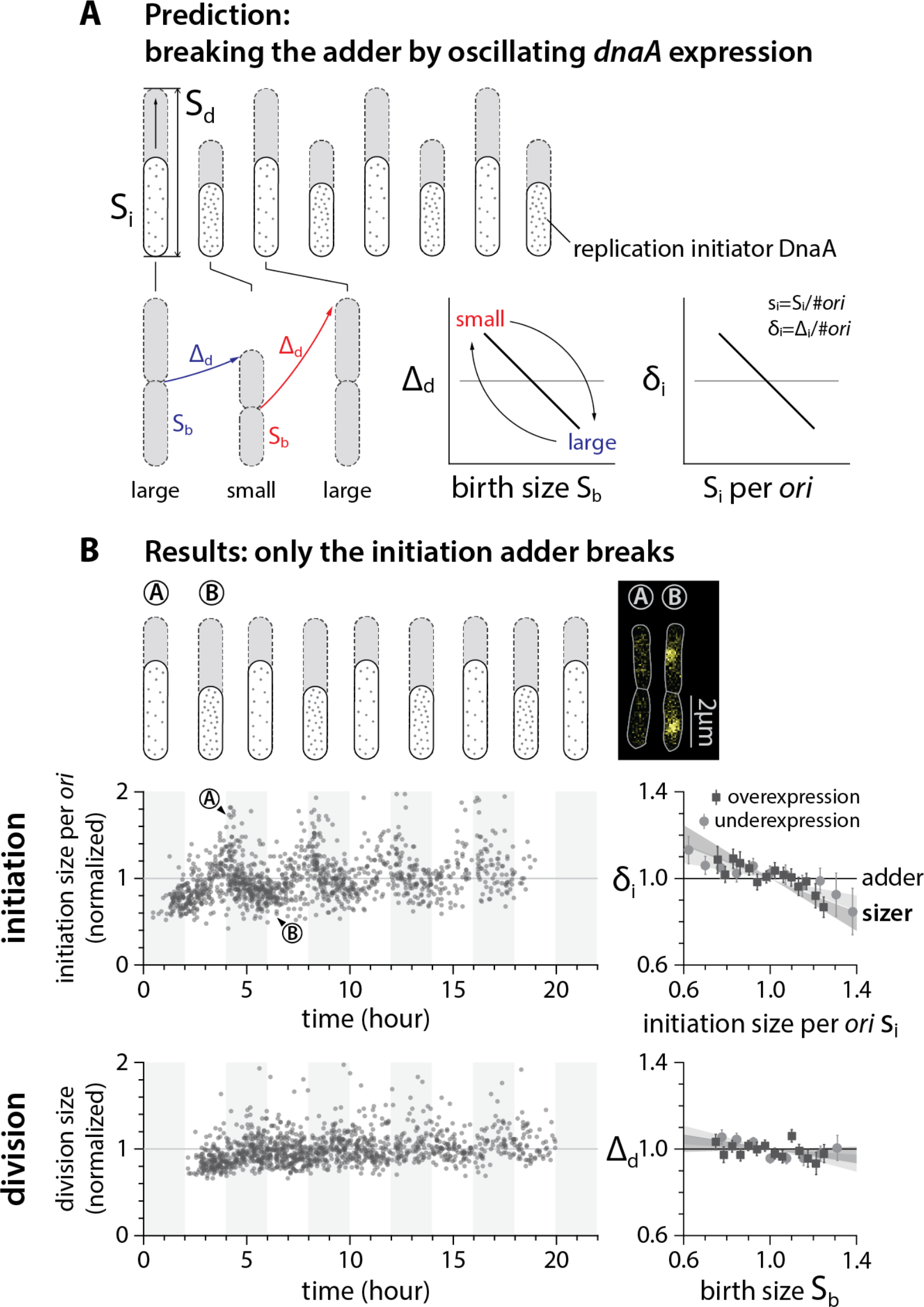
Dynamic perturbation of DnaA production breaks the initiation adder but not the division adder. **(A)** The array of cell cartoons represent the initiation and division events in consecutive generations. The initiation adder and the division adder can be perturbed via a periodic modulation of the initiation process. Ideally, a period T=2τ yields the most negative mother-daughter autocorrelation. **(B)** Periodic underexpression or overexpression of DnaA dynamically modulated initiation size per *ori*, but not division size. This caused deviations in the initiation adder but not in the division adder. The period of the oscillation (T) of the inducer IPTG is about 4τ instead of 2τ considering the actual response time of initiation to DnaA induction (Supplemental Theory III.E). The cell images overlay phase contrast with fluorescence of replisome markers. The plots on the left show the data of periodic underexpression of *dnaA*. The grey and white stripes represent the induction-to-underexpression cycles between 200 μM and 0 μM IPTG (see Fig. S3 and experiment details in Table S5). Each dot corresponds to one division cycle of a single cell. The grey solid lines represent the time averages. In the correlation plots, the red shaded area represents the 95% confidence interval of linear fit to the respective raw scatter plot. The results in (**B**) refute the co-regulation hypothesis that division timing is determined by initiation timing via a fixed τ_cyc_. The independence of initiation and division controls warrants a major revision in interpreting the Helmstetter-Cooper model at the single-cell level (Supplemental Theory II). See detailed experimental information in Fig. S3, and Table S3 and S5 [43].

In our actual experiments, we had to use *T ≈* 4*τ* because of a significant induction and dilution time of *dnaA* (Fig. S4A and Supplemental Theory III.E) [43]. Nevertheless, these experiments showed clear oscillations in the initiation size without noticeable changes in the growth rate (Fig. 3B and Fig. S3). The measured autocorrelations of initiation size decreased accordingly, decisively breaking the initiation adder as predicted by our simulations (Fig. 3B).

### The division adder is independent of initiation control, refuting the co-regulation hypothesis

To our surprise, and counter to the co-regulation hypothesis [17], the division adder remained intact even when the initiation adder no longer held by periodic induction of DnaA expression (Fig. 3B). When initiation is delayed, the division size remains mostly constant as long as replication termination timing does not exceed the division timing. Thereafter, initiation delay causes an increase in division size (Fig. S3D) [20]. Decoupling between the initiation adder and the division adder suggested that the timing of cell division, in fact, has its own independent control at the level of gene expression [2, 24, 44, 45]. We further reasoned that division timing is regulated by the dynamics of proteins and precursors required *for division*, rather than that of DnaA and other proteins required for replication initiation. We thus set out to break the division adder without breaking the initiation adder.

### Dynamic perturbation to the synthesis of division initiators breaks the division adder but not the initiation adder

Cell division requires assembly of more than a dozen types of proteins and biomolecules at the future septum, including the enzymes required for synthesis of the septal cell wall. We elected to use the tubulin-like GTPase, FtsZ, because (like DnaA) it is highly conserved and assembles in an expression-level dependent manner. FtsZ-ring formation is required for assembly of all other components of the cell division machinery [46, 47], and the timing of division has been shown to be systematically delayed when FtsZ is underexpressed [48, 49]. FtsZ also has practical advantages since its genetic and cytological properties have been extensively characterized [50].

To determine if oscillations in FtsZ production break the division adder in the same manner that oscillations in DnaA break the initiation adder, we adopted a strain in which the wild-type *ftsZ* was expressed under the control of an inducible promoter (Fig. 4A) [48, 51]. We also tracked replication dynamics using the fluorescent replisome marker. When we periodically underexpressed *ftsZ* with *T ≈* 4*τ*, cell size at division oscillated with the same period, exclusively breaking the division adder without affecting initiation size (Fig. 4B, Figs. S4A and S4D). In addition, we obtained the same results by periodically producing the division inhibitor protein SulA (Fig. 4C and Fig. S4B).

**Figure 4:**
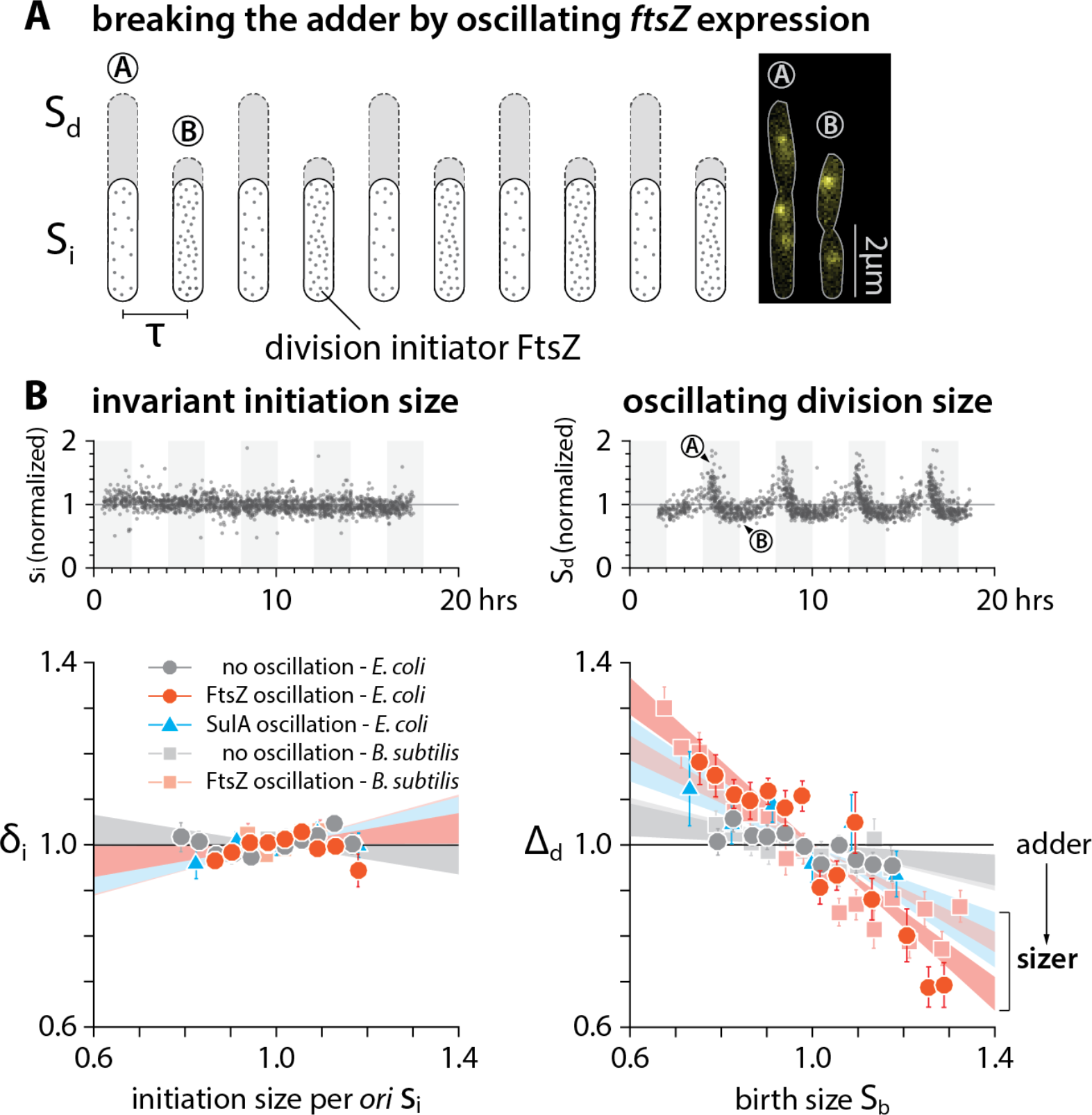
Dynamic perturbation to FtsZ production breaks the division adder but not the initiation adder. **(A)** The array of cell cartoons represent the initiation and division events in consecutive generations, similar to Fig. 3A. Dynamic modulation of division protein FtsZ oscillates the division size but not the initiation size. In *E. coli*, the native promoter of *ftsZ* was replaced with a *P*_tac_ promoter. In *B. subtilis*, the endogenous *ftsZ* was deleted while an alternative copy of *ftsZ* under *P*_xyl_ was inserted at a different loci of the chromosome (see Figs. S3 and S4; STAR Methods). Both strains produce fluorescent fusion proteins of DnaN for replication tracking. The cell images overlay phase contrast with fluorescence of replisome markers in *E. coli*. **(B)** Dynamic modulation of FtsZ was conducted at the oscillation period 4τ, with IPTG concentration oscillating between 0 µM and 10 µM for *E. coli*, and xylose concentration between 0.1% w/v and 1% w/v for *B. subtilis*. The period of dynamic modulation of SulA for *E. coli* was also 4τ, with IPTG concentration oscillating between 0 µM and 40 µM (see Fig. S4). Under FtsZ-level oscillations, both size homeostasis of *E. coli* and *B. subtilis* become sizer-like, yet their initiation adderness remains intact. The upper plots present the *E. coli* data. Each dot corresponds to one division cycle of a single cell. The grey and white stripes represent the induction-to-underexpression cycles between 10 μM and 0 μM IPTG, and the grey solid lines represent the time averages (also see Fig. S4). In the correlation plots, the variables are normalized by their means and each shaded area represents the 95% confidence interval of the linear fit to the respective raw scatter plot.

We repeated our experiments under different induction levels of *ftsZ* keeping the induction frequency same as before. The degree of deviations from the division adder systematically increased, yet the initiation adder remained intact, underscoring the independence between the initiation adder and the division adder in cell-size homeostasis (Fig. 4B, Figs. S4A and S4D; Supplemental Theory III).These results also show that cell division processes exclusively drive cell-size homeostasis in *E. coli*.

### *E. coli* and *B. subtilis* likely share the same mechanistic origin of cell-size homeostasis

Next, we asked whether the exclusive role of cell division on size homeostasis, and its independence of initiation control, is a general feature of bacteria. To explore this idea, we repeated the FtsZ oscillation experiments in a model Gram-positive bacterium *B. subtilis*. *B. subtilis* is particularly interesting because, while DnaA and FtsZ are conserved in both bacteria, the mechanisms governing both replication initiation and division in *B. subtilis* differ in fundamental ways from those in *E. coli* [47, 52]. We constructed a strain that encodes *ftsZ* under an inducible *P*_xyl_ promoter as the sole source of FtsZ, in addition to the functional DnaN-GFP fusion protein (Fig. 4A) [53]. Together, these constructs permit periodic modulation of FtsZ levels and simultaneous tracking of replication dynamics.

Similar to *E. coli*, *B. subtilis* exhibited systematic deviations from the division adder when *ftsZ* expression was varied periodically. Furthermore, we found *B. subtilis* to be an initiation adder regardless of the oscillations (Fig. 4B, Figs. S4C and S4E). These results strongly suggested that *E. coli* and *B. subtilis* share the same mechanistic origin of cell-size homeostasis.

### Mechanistic origin of cell-size homeostasis in bacteria: threshold and balanced biosynthesis

Our data so far indicated it is possible to break the adder phenotype using periodic oscillations in the production rate of cell-cycle proteins to perturb initiation size or division size (DnaA for the initiation adder and FtsZ for the division adder). This finding suggests that balanced biosynthesis of cell-cycle proteins is likely an important requirement for the adder phenotype.

In balanced biosynthesis, the protein production rate is proportional to the rate cells increase their volume, irrespective of the protein concentration at birth. Cells therefore on average add a fixed number of proteins per unit volume during growth, and the total number of newly synthesized proteins is directly proportional to the total cell volume added since birth. Assuming balanced biosynthesis, a cell would be a division adder if division is triggered after accumulating a fixed number of division proteins, namely a fixed volume (Fig. 5A; Supplemental Theory II). In other words, two experimentally testable assumptions are sufficient to explain the adder phenotype: (1) Threshold -- accumulation of initiators and precursors required for cell division to a fixed number, and (2) Balanced biosynthesis -- maintenance of their production proportional to volume growth.

**Figure 5:**
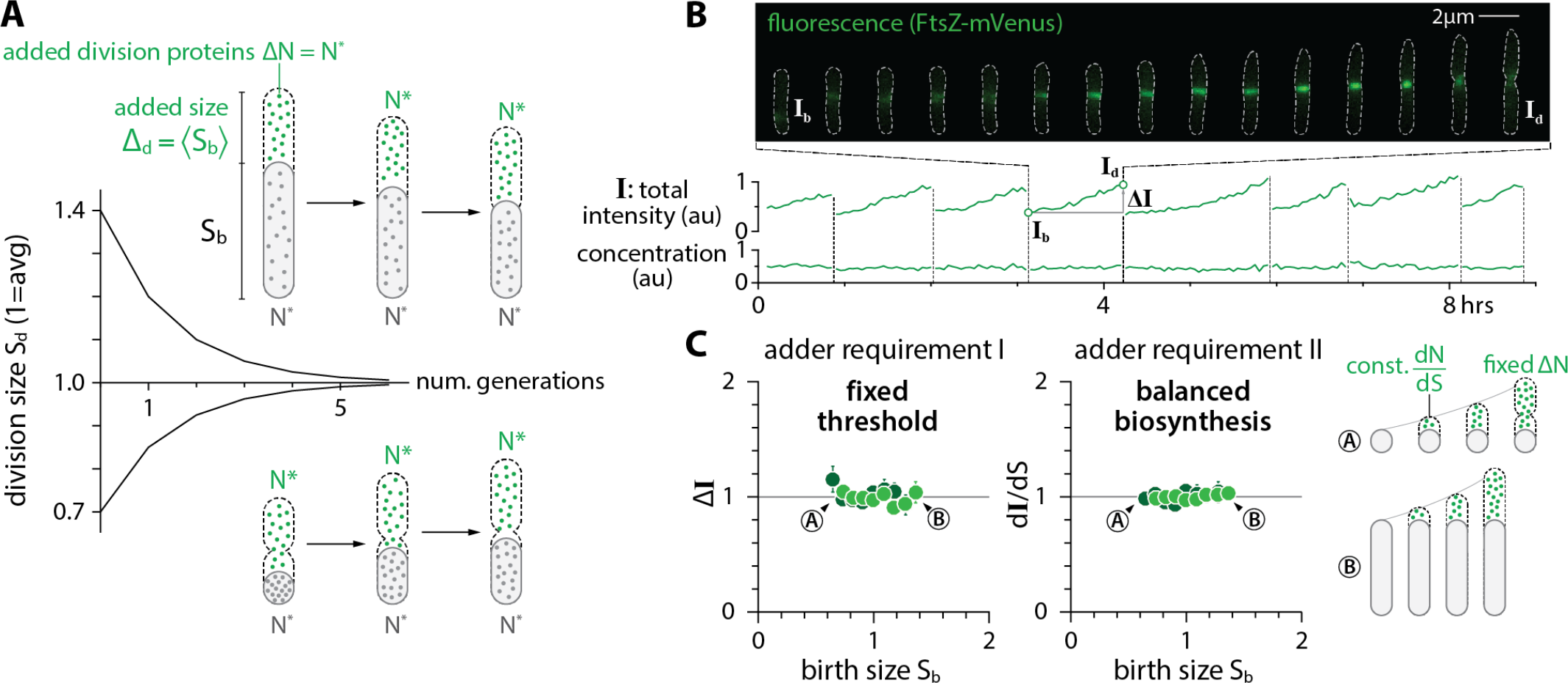
The mechanistic origin of the adder. **(A)** The adder phenotype requires accumulation of division proteins to a fixed amount 2N* to trigger division, and their balanced biosynthesis during growth. Under these conditions, newborn cells are born either larger or smaller than the population average, but they on average contain N* division proteins. The two adder requirements ensure that both small-born and large-born cells add a constant size (or N* division proteins) in each generation. **(B)** A typical timelapse sequence with FtsZ-mVenus. The total intensity was obtained by integrating the FtsZ-mVenus fluorescence intensity over the entire cell, which increases steadily from birth to division, tracking the total size of the cell. As a result, the FtsZ-mVenus concentration stays nearly constant within fluctuations. **(C)** The synthesis and accumulation of FtsZ in *E. coli* cells fulfills both requirements for adder. The total added number ΔN (estimated by the added fluorescence ΔI) and the synthesis per unit volume dN/dS were constant and independent of cell size at birth (see Fig. S5). Symbol colors indicate repeats of experiments.

To test this idea, we measured the production rate and the accumulation of FtsZ in single cells. We adopted an *E. coli* strain expressing a nearly functional fusion *ftsZ*-mVenus as the sole endogenous copy of *ftsZ* [54]. We used the total fluorescence per cell, **I**, to estimate the total copy number of FtsZ per cell (Fig. 5B; STAR Methods), and indeed found that **I** increased proportionally to the increase in cell volume in individual cells (Fig. S5B). The production of FtsZ-mVenus per unit volume, d**I**/dt, during growth was independent of the cell size or FtsZ concentration at birth, consistent with the balanced biosynthesis hypothesis (Fig. 5C; Fig. S5C) [55, 56]. Furthermore, the total accumulation of FtsZ-mVenus between birth and division, Δ**I** = **I**_d_ - **I**_b_, was also constant and independent of cell size or FtsZ concentration at birth **I**_b_, supporting the threshold hypothesis (Fig. 5C; Fig. S5C) [21, 25, 57–59].

These results extend the previous observations that the Z-ring appears at mid-cell shortly after birth and FtsZ accumulates at the Z ring steadily over the course of the division cycle [60–63] (Fig. 5B, Fig. S5A). We also found that the onset of constriction coincides with when the total Z-ring intensity reaches its max value. The maximal Z-ring intensity was independent of the cell size or FtsZ concentration at birth (Figs. S5A and S5C), reinforcing the molecular basis for the threshold model. As explained below, we further verified these hypotheses in our oscillation experiments.

### Testing the mechanism of the adder in the FtsZ oscillation experiments

As the steady-state growth experiments supported the threshold and balanced biosynthesis hypotheses, we further tested them in new oscillation experiments. We combined the *ftsZ-mVenus* strain with the inducible system used in the oscillation experiments (Fig. S6A; STAR Methods). As expected, FtsZ-mVenus concentration oscillated in response to the periodic induction, while the division size exhibited clear out-of-phase oscillations (Fig. 6A; Supplemental Theory III). Despite the oscillations in FtsZ-mVenus concentration, the maximal Z-ring intensity at mid-cell and the total added fluorescence remained remarkably constant throughout the experiments regardless of the FtsZ concentration or cell size at birth (Fig. 6A; Fig. S6C).

**Figure 6:**
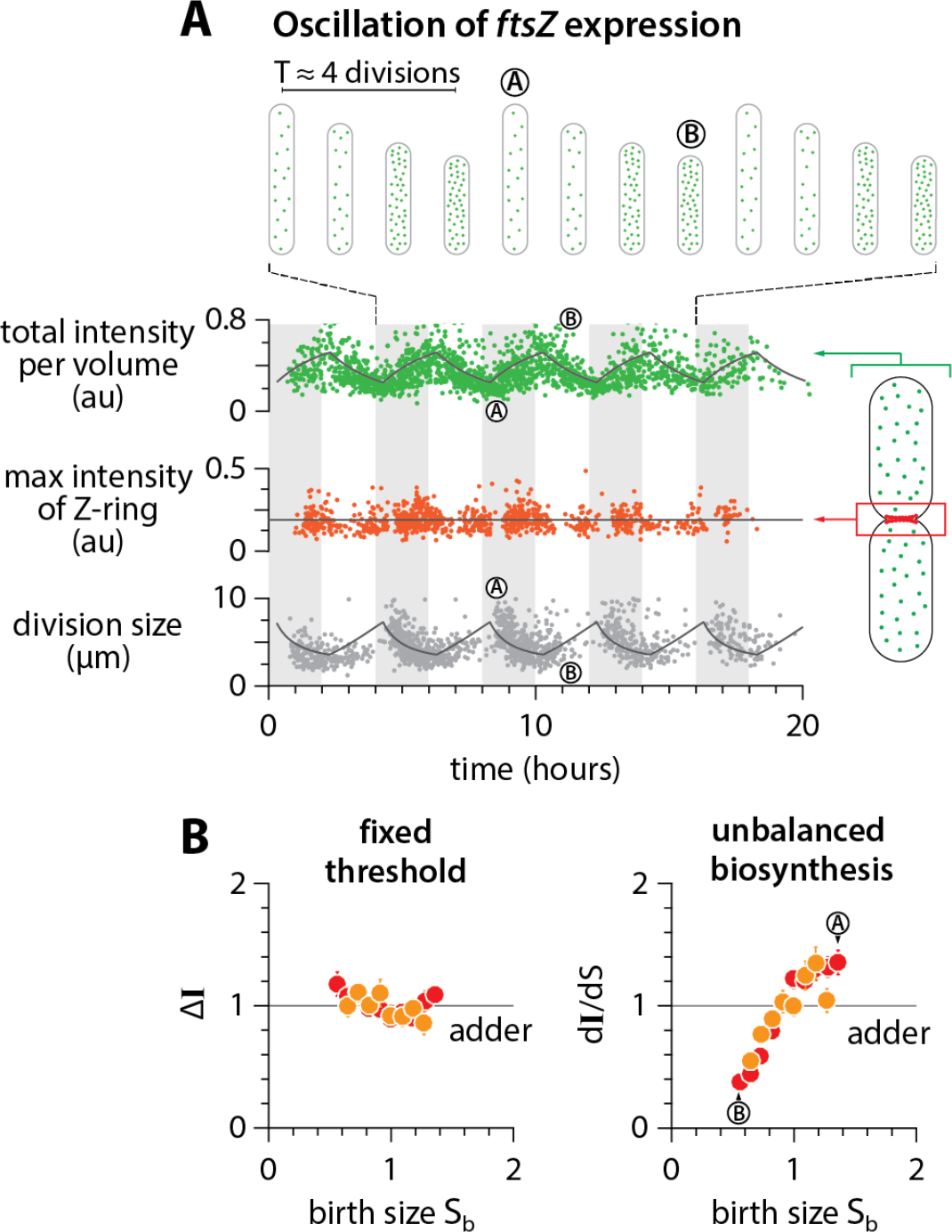
Testing the mechanism of adder in the FtsZ oscillation experiment. (A) Total FtsZ-mVenus concentration oscillates in response to the periodic induction, but the threshold amount at the septum is invariant. The amount of FtsZ accumulated in the septum ring was estimated by integrating the fluorescence intensity within a fixed area enclosing the mid-cell region (STAR Methods; Fig. S5). The solid lines represent the prediction based on balanced biosynthesis and threshold model (see Supplemental Theory III). (B) Despite the oscillations in the total FtsZ concentration, the total added fluorescence ΔI and the max Z-ring intensity remain invariant with respect to birth size (see also Fig. S6C). In contrast, the production rate of FtsZ was variable due to oscillations (see Supplemental Theory III for the prediction of d**I**/dS vs. S_b_). Symbol colors indicate repeats of experiments.

The sizer-like behavior can also be explained by the out-of-phase oscillations in the FtsZ concentration and division size. During periodic induction of *ftsZ*, small-born daughter cells contain higher concentrations of FtsZ, because their mother cells accumulated FtsZ at a faster rate during high-level induction and therefore divided early. These new small-born daughter cells in turn experience low-level induction (Fig. 6B, bottom), thus accumulate FtsZ to the fixed threshold number at a slower-rate and elongate longer to reach division (Fig. 6B, top). Indeed, the added size Δ_d_ vs. newborn size s_b_ shows a characteristic sizer-like negative slope (Fig. S6C; Fig. 4).

### How to restore the adder phenotype in slow-growing *E. coli*

While a wide range of evolutionarily divergent organisms are adders [15], a major exception has also been reported for *E. coli*. Specifically, Wallden *et al*. reported that size homeostasis, during slow growth in nutrient limitation, deviates from the adder [19]. We re-analyzed the published data in Wallden *et al*. and also performed our own experiments in the same growth condition. In contrast to Wallden *et al*., we found that the slope −0.31 in the Δ_d_ vs. newborn size S_b_ is in fact much closer to the adder (slope = 0) than the sizer (slope = −1) in both experiments. At the same time, the deviations from the adder is statistically significant (p-value = 1.4⨉10^−6^) as pointed out by Wallden *et al*.

Wallden *et al*. provided a possible explanation for their observation, based on the Helmstetter-Cooper model at the single-cell level. That is, the cell size at division can be written as S_d_ = s_i·_exp(λ*τ*_cyc_) for non-overlapping cell cycles when cells grow in nutrient poor media. If s_i_ is invariant while *τ*_cyc_ is inversely proportional to λ at the single cell level because all biosynthesis equally limiting in slow growth conditions [19], λ_·_*τ*_cyc_ = constant and therefore the invariant s_i_ implies S_d_ is fixed regardless of the birth size, thus the sizer. While elegant, this explanation is based on the co-regulation hypothesis and predicts both slope in Δ_d_ vs. s_b_ and Δ_i_ vs. s_i_ to be −1. For this prediction to be valid, both λ_·_*τ*_cyc_ and s_i_ should be uncorrelated with birth size, which is in conflict with our data (Fig. S7A). Indeed, our data obtained from the same growth condition as Wallden *et al*. instead shows that slow-growing *E. coli* is an initiation adder, and mildly deviates from the division adder (Fig. 7; Figs. S7B and S7C).

**Figure 7:**
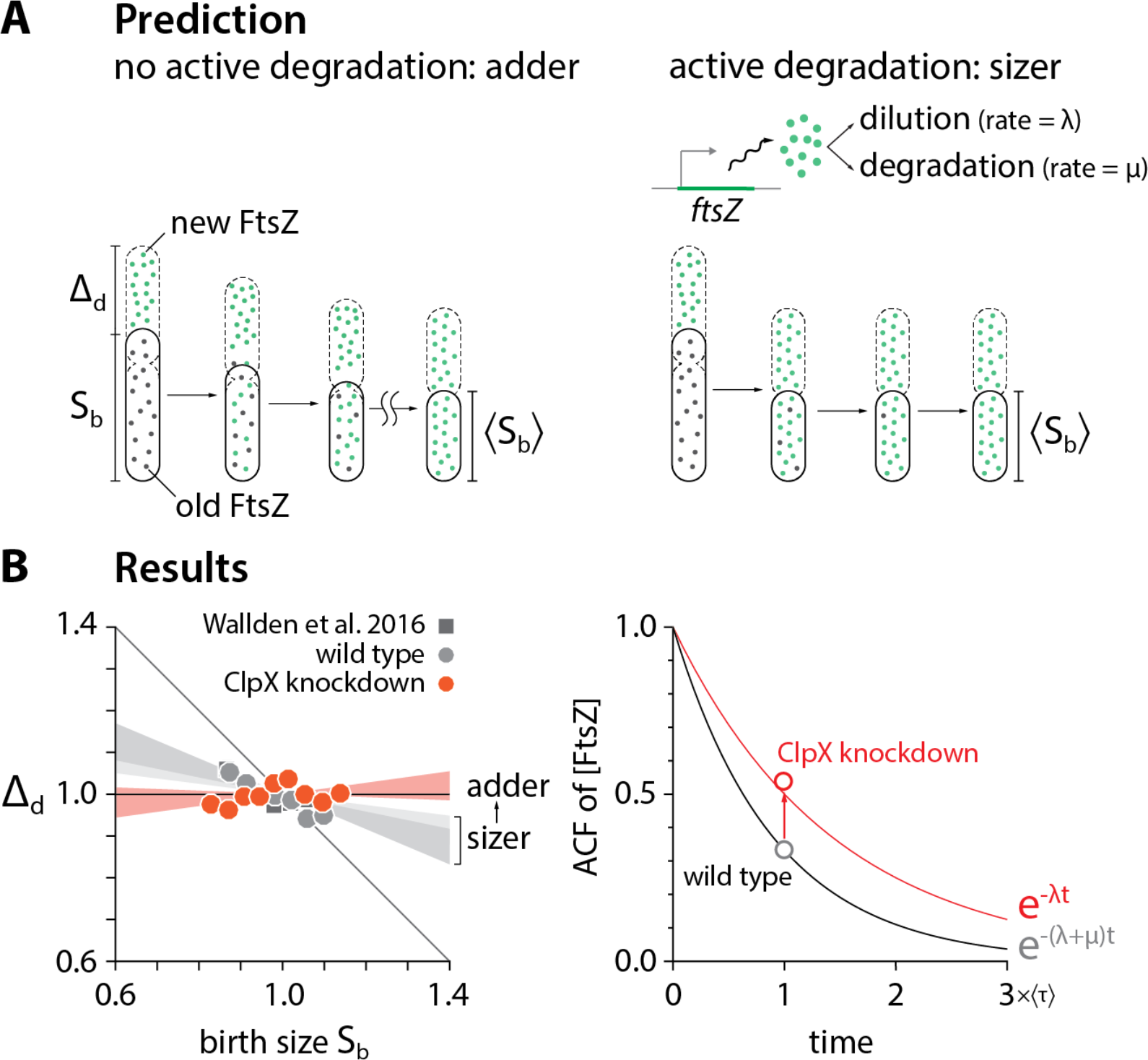
Restoring the division adder. **(A)** In fast-growing cells, the constant production and dilution of division proteins explains adder. Differently, in slow-growing cells, a significant active degradation of FtsZ is sufficient to deviate the homeostasis of both protein concentration and cell size. **(B)** Slow-growing *E. coli* cells are initiation adders (this work; Fig. S7C) but not division adders (this work and Wallden *et al*. 2016; the doubling time was about 4 hours). Repressing *clpX* expression via tCRISPRi restored the division adder. Each shaded area represents the 95% confidence interval of linear fit to the respective raw scatter plot. The mother-daughter autocorrelation of division size was less than 0.5 in slow-growing cells, indicating a more sizer-like behavior. The autocorrelation function of FtsZ concentration was also less than 0.5. The repression of *clpX* expression alleviated both deviations (see STAR Methods).

Our intuition for the discrepancy was that the slow-growing *E. coli* violates one or both requirements for the division adder (Fig. 5). Specifically, we asked if FtsZ is actively degraded in slow growth conditions (see, also, [64, 65]), resulting in a higher turnover rate. Active degradation of FtsZ should decrease both autocorrelation of FtsZ concentration and division size (Figs. 7A and 7B; STAR Methods). We further predicted that suppression of the activity of FtsZ would restore the division adder.

We tested our prediction by repressing *clpX* expression using our tCRISPRi system [32]. We found that *clpX* repression was indeed sufficient to fully restore the division adder (Fig. 7B). The initiation adder was intact with or without the *clpX* repression (Fig. S7C). These results provide strong experimental evidence for balanced biosynthesis and threshold as the requirements for cell-size homeostasis to be an adder.

### Relationship with previous works

Balanced biosynthesis and threshold are general concepts in biology and have been implied in a number of papers since the 1970s from replication initiation [31] or cell division [3, 21, 59] in bacteria to mitotic control in eukaryotes [66]. The threshold model has also been explicitly put forward as the trigger of cell division as starved *E. coli* cells resume growth [64]. A recent work addressed whether cell shape contributes to size control [25], but we recognize its core implicit assumptions are balanced biosynthesis of cell-wall precursors and their accumulation to a threshold to build the septum.

Previous work independently showed that *E. coli* is a division adder but also questioned whether size control is implemented at initiation or division [2]. We have shown that division drives size homeostasis in *E. coli* and *B. subtilis*, but they are both initiation and division adders in steady state (Fig. 1D). The independence between the two types of adders can only be revealed in non-steady-state growth (Figs. 3 and 4). Subsequent analysis [67] has shown that the experimental evidence in Campos *et al*. may in fact agree with the initiation adder. As we show in Supplemental Theory IV, the initiation control model in [2] can result in unstable cell size regulation, but can be corrected when growth by a constant size *per origin* is implemented at initiation in steady-state growth (see Supplemental Theory IV).

Another notable proposal for cell-size control in *E. coli* is a negative feedback imposed on cell size [68]. The hypothetical feedback exclusively relied on transient “oscillations” observed in the autocorrelation function (ACF) of cell size in experimental data and simulation data of an autoregressive model. However, it is well known that the ACF of the autoregressive model they used is an exponential function, in contradiction with the claimed oscillations. In other words, it is likely that the “oscillations” observed in both experimental and simulation ACFs are fortuitous, and caused by an insufficient sampling (approximately N=70 generations in each lineage) that fails to produce statistically meaningful autocorrelation coefficients.

## CONCLUSION AND OUTLOOK

Altogether, we have shown that it is cell division [2, 3, 21, 25], not replication initiation [17, 37, 67], that drives cell-size homeostasis in bacteria. Initiation control is important in cell-size control, rather than cell-size homeostasis, because initiation defines unit cellular volume (or “unit cell”) so that the average cell size in any steady-state population is given by the sum of all unit cells [20]. From the cell-cycle control point of view, we showed that initiation and division are independently controlled in both *E. coli* and *B. subtilis*, thereby providing a conclusive answer to the long-standing question whether replication initiation regulates cell division in bacteria [34].

The mechanism underlying the adder phenotype for size homeostasis reduces to two biological hypotheses: (1) balanced biosynthesis of division proteins and precursors and (2) their accumulation to a threshold number in individual cells. In this work, we provided direct experimental evidence that support these two hypotheses for cell-size control and homeostasis. In our view, a next major question for the future is how a threshold model is implemented at the molecular level in division control and cell cycle control in general, while continuing a constructive dialog between quantitative phenomenological principles and mechanistic investigation.

The mechanism of adder has obvious implications for its applicability to other biological problems such as homeostasis of organelle content [56]. From an evolutionary point of view, cell-cycle dependent degradation of cyclins may explain why some eukaryotes show clear departure from adder by actively modulating physiological memory. But perhaps a more curious case is the mechanism of size homeostasis of the First Cells or synthetic cells, for which the simplicity of balanced growth makes adder an intriguing possibility.

## ACKNOWLEDGEMENTS

We thank Dongyang Li, Rodrigo Reyes-Lamothe, Tsutomu Katayama, Anders Løbner-Olesen, Harold Erickson, William Margolin, and Paul Wiggins for providing the strains. This work was supported by the Paul G. Allen Family Foundation, Pew Charitable Trust, NSF CAREER grant MCB-1253843, and NIH grant R01 GM118565-01 (to S.J.), and R35-400 GM127331 (to P.A.L).

## AUTHOR CONTRIBUTIONS

F.S., G.L.T., and S.J. conceived the study. All authors conducted research and wrote the paper.

## STAR METHODS

### CONTACT FOR REAGENT AND RESOURCE SHARING

Further information and requests for resources and reagents should be directed to and will be fulfilled by the Lead Contact, Suckjoon Jun.

### EXPERIMENTAL MODEL AND SUBJECT DETAILS

#### Strains

##### A. E. coli

###### Strain background

The *E. coli* strains used in all mother machine experiments are with either K-12 NCM3722 or K-12 MG1655 background. Both strains were sequenced and extensively tested in previous studies [20, 69, 70]. Strains with NCM3722 background were only used in steady-state growth, and strains with MG1655 background were used in both steady-state and oscillation experiments. Detailed information of strain genotypes are included in Suppl. Tables 1 and 2 below.

###### Parent strains

Some of the strains used in this study were constructed based on existing parent strains which have been tested and published. The original strain with the DnaN-YPet replisome marker was a kind gift of Rodrigo Reyes-Lamothe [71]. The strain for DnaA knockdown is based on the tunable CRISPR interference system developed in the Jun lab [32]. The DnaA overexpression strain used in the steady-state inhibition experiment is based on a strain with a plasmid carrying an extra copy of *dnaA* under P_*lac*_ promoter which was a kind gift of Tsutomu Katayama [72]. The DnaA overexpression strain used in the oscillation experiment is based on a similar strain with a plasmid carrying an extra copy of dnaA under P_*lac*_ promoter which was a kind gift of Anders Løbner-Olesen [43]. The construct of FtsZ-mVenus was a kind gift of Harold Erickson. In this construct, mVenus is inserted into the linker between domains of FtsZ, which has minimal effect on the function of FtsZ [54]. The system of P_*tac*_::FtsZ was developed in Miguel Vicente’s lab [48, 51], and the strain VIP205 containing this system was a kind gift of William Margolin. We are grateful to the researchers mentioned above for these gifts of strains.

##### B. B. subtilis

###### Strain background

The *B. subtilis* strains used in all mother machine experiments had JH642 background which is autoxophoic and requires supplementation of tryptophan, phenylalanine and threonine.

###### Parent strains

The original strain with DnaN-GFP replisome marker was developed in Alan Grossman’s lab [53] and was a kind gift of Paul Wiggins whose lab has conducted the cell cycle measurement using this strain [28].

See detailed strain information in Suppl. Tables 1 and 2.

#### Growth media

For *E. coli*, we used MOPS or M9 minimal media supplied with different carbon sources and amino acids. For *B. subtilis*, S750 minimal media with different carbon sources and other supplements were used. The details of the media used are listed in Suppl. Tables 3 and 4.

#### Experimental conditions and sample size

The detailed growth conditions, experimental parameters and samples size of each experiment included in this study are listed in Suppl. Table 5.

## METHODS DETAILS

### Microfluidics

Mother machine microfluidic devices were used in this study to monitor single cell growth for 10-50 generations in both steady state and oscillation experiments. Syringe pumps (PHD Ultra, Harvard Apparatus, MA) were programmed to infuse fresh growth media into microfluidic device at either a constant rate or in an oscillatory manner.

### Cell preparation

Before every time-lapse imaging, cells were picked from a single colony on an agar plate which was streaked no more than 7 days before use. The cells were inoculated into 1 mL lysogeny broth (LB) with proper selection antibiotics. After shaking for 12-18 hours at 30°C or 37°C in a water bath shaker, cells were diluted 1,000-fold into 2 mL of defined medium same as that used in the microfluidic experiments. After shaking at 37°C in the water bath till OD_600_ = 0.1-0.4, cells were diluted again 100-to 1,000-fold into the same medium and shaken at 37°C in water bath till OD600 = 0.1-0.4. The cell culture was then concentrated 10-to 100-fold and injected into a microfluidic mother machine device via a 1 mL syringe. 0.5 mg mL^−1^ BSA (Bovine serum albumin, Gemini Bio Products, CA) was added to the fresh growth media to reduce the adhesion of cells to the surface of microfluidic channels. The media were then added to 10 mL, 20 mL or 60 mL plastic syringes (BD) with 0.22 µm filters (Genesee Scientific, CA) for the time-lapse imaging. All imaging experiments were conducted at 37°C in an environmental chamber [3].

### Microscopy and image acquisition

We performed simultaneous phase-contrast and epifluorescence imaging on an inverted microscope (Nikon Ti-E) with Perfect Focus 3 (PFS3), 100× oil immersion objective (PH3, numerical aperture = 1.45), Obis lasers 488LX or 561LS (Coherent Inc., CA) as fluorescence light source, and Andor NEO sCMOS (Andor Technology) or Prime 95B sCMOS camera (Photometrics). The laser power was 18 mW for 488 nm excitation and 17 mW for 561 nm, respectively. Exposure time was set between 50-200 ms. Imaging frequencies were calibrated at about 20 frames per doubling time such that no physiological effects on the cells were discernible.

### Image processing

#### A. Cell segmentation, lineage reconstruction and cell dimension measurement

We developed custom imaging processing software using Python 2.7. The work flow is as follows. First, phase contrast images of each field-of-view (FOV) were sliced into small images each containing one growth channel of the mother machine device. Second, to enhance the contrast, the empty channels were subtracted from those containing cells in the same FOV. Third, subtracted images were thresholded using Otsu’s method to create a binary mask and then applied with morphological operations and a distance filter to create labeled markers. Markers were used to seed a watershedding algorithm on the subtracted images to create the segmented image. Lastly, lineages were constructed using a decision tree which tracked the time-evolution of the cell segments. The cell dimension was measured based on Feret diameter method: the cell length was calculated as the intercept of the cell’s long axis through the cell center and the outline of the segmented cell, and the cell width was calculated as the the mean of intercepts of the cell’s short axis through the cell quarter positions and the outline of the segmented cell.

#### B. Replisome foci analysis

The images of replisome markers were processed using the segmentation and lineage information from the phase contrast images of the same cells. Background subtraction was done by subtracting the mean value of multiple empty channels from those containing cells in the same FOV. Unfiltered fluorescence foci were identified as local maxima using a Laplacian of Gaussian method (blob_log function in Skimage v0.11.3). The localization for each identified focus was obtained using 2-D elliptic Gaussian fitting, and all foci were filtered again according to their peak-to-background value. The total fluorescence of each replisome focus was estimated as the total intensity of each blob. The distribution of fluorescence intensity of all foci was plotted and fitted with a double Gaussian distribution. The position of the second peak of the fitted Gaussian was typically two times that of the first peak, suggesting that two fluorescence foci were often spatially overlapping and undistinguishable due to the diffraction limit (Fig. 1B). Therefore, a focus with higher probability of falling into the second peak region (integral of the intensity distribution between that foci intensity and the second peak > that of the first peak) was counted as two focus. The single-cell cell cycle analysis was carried out using a custom Matlab software. Intracellular positions and intensities of all foci in the same cells were plotted against time for the whole cell lineage. The start and end points of each foci trace were determined as the replication initiation and termination with respect to division cycles (Fig. 1A, Fig. S1).

#### C. FtsZ-mVenus concentration analysis

Fluorescence images of FtsZ-mVenus were used to estimate the total amount FtsZ per cell, the total concentration of FtsZ, the total fluorescence of the Z-ring and the cytoplasmic concentration of FtsZ. Compared to the replisome marker images, extra calibration of systematic errors was done as follows. (1) We corrected the photobleaching effect by truncating the time points when the average fluorescence of cells have not reached steady state. (2) The illumination of the laser was often non-uniform across the FOV. The profile of illumination was obtained from the average intensity of all cells in the same FOV. The fluorescence intensity of each cell was thus calibrated according to the profile and their position in the FOV. (3) The FOV-to-FOV variations were typically less than 5%, so no calibration was applied. The first 5-10 generations of the time-lapse images were discarded to ensure that photobleaching reaches stationarity of the timelapse data. The total fluorescence of FtsZ per cell was used to estimate the total amount FtsZ per cell. The total fluorescence normalized by cell volume was used to estimate the total concentration of FtsZ. The amount of FtsZ in the mid-cell area was quantified by integrating the fluorescence intensity within a fixed box with dimensions of 1 µm along cell long-axis and 1.5X cell width along short axis. This area is centered at max intensity position of the line profile along cell long-axis. This quantity was used as an approximation of the total fluorescence in the Z-ring. The cytoplasmic concentration of FtsZ was estimated as the total fluorescence within an area of the same size centered at a cell-quarter position along the cell long-axis. The cytoplasmic concentration of FtsZ was shown to be much higher than the cellular autofluorescence. We showed this by co-growing and imaging the FtsZ-mVenus strain and wildtype parental strain in the same mother machine device. The autofluorescence level of the wild type strain is less than 10% of the cytoplasmic FtsZ-mVenus fluorescence (Fig. S5D).

### Stochastic simulations of the Helmstetter-Cooper model

To investigate what determines cell-size homeostasis we developed stochastic simulations of the Helmstetter-Cooper cell cycle model (Cooper and Helmstetter 1968; Donachie 1968). In this model, three coarse-grained physiological parameters describe the progression of the cell cycle and cell size: the growth rate *λ*, the cell size per origin at replication initiation *s*_i_, and the length of cell cycle *τ*_cyc_ = *C+D*, namely the duration that spans one complete round of replication (C period) and division that corresponds to replication termination (D period). We introduced stochasticity to these parameters (*λ*, *s*_i_, *τ*_cyc_) and numerically probed the resulting behavior of cell-size homeostasis (Fig. 1C). See more details in Supplemental Theory I. The stochastic fluctuations constituted a 9-dimensional physiological space consisting of and three coefficient-of-variations (CVs), three cross-correlations and three autocorrelations (Fig. 2A), with each physiological dimension representing specific biological constraints. For instance, positive autocorrelations in the growth rate *λ* mean that on average fast-growing mother cells produce fast-growing daughter cells. When these refinements were added, our stochastic simulations self-consistently reproduced the experimentally observed adder behavior for all tested growth conditions without any adjustable parameters (Supplemental Theory I). In Fig. 2A, we set out to systematically vary physiological parameters along all nine dimensions to probe the adder behavior. Each simulation generated a lineage of 10,000 cells. The adder correlation ρ(*∆*_d_, *S*_b_) was defined as the Pearson correlation between the variables *∆*_d_ and *S*_b_ in the simulated lineage. We adopted the same definition for the initiation adder correlation ρ(*δ*_i_, *s*_i_). Eventually, we found that deviations in the autocorrelation of initiation size per *ori s*_i_ from 0.5 significantly affected the division adderness. In contrast, deviations from adder resulting from other perturbations were weaker or less systematic, reinforcing the general robustness of adder observed in our inhibition experiments (Fig. 1D). This sensitivity of adder to *s*_i_ autocorrelation is clearly seen in the fraction of physiological space represented by adder (Fig. 2A). It is also intuitive since in the Helmstetter-Cooper model, division timing is regulated by chromosome replication initiation. As reference physiological values, we used experimental measurements obtained for strain NCM3722 in slow growth condition (MOPS minus NH_4_Cl, 0.4 % glucose, 5 mM arginine). Namely, where appropriate we parametrized the joint probability distribution using the mean and coefficient-of-variations:

| Variable | $\lambda$ | $\tau_{cyc}$ | $s_i$ |
| --- | --- | --- | --- |
| Mean | 0.693 | 0.7 | 1 |
| CV | 15% | 15% | 10% |

and the Pearson cross-correlations and autocorrelations:

| Pearson correlation | Value |
| --- | --- |
| $\rho(\lambda, \tau_{\text{cyc}})$ | -0.5 |
| $\rho(\lambda, s_i)$ | -0.2 |
| $\rho(\tau_{\text{cyc}}, s_i)$ | -0.3 |
| $\rho(\lambda^{(n+1)}, \lambda^{(n)})$ | 0.5 |
| $\rho(\tau_{\text{cyc}}^{(n+1)}, \tau_{\text{cyc}}^{(n)})$ | 0.3 |
| $\rho(s_i^{(n+1)}, s_i^{(n)})$ | 0.5 |

Note that we chose the generation time as unit of time and the cell size per origin at initiation as unit of volume. For this particular condition, the generation time was $\ln(2)/\langle\lambda\rangle = 112$ minutes and the cell size per origin at replication initiation was $s_i = 0.30 \mu\text{m}^3$ .

Note that we chose the generation time as unit of time and the cell size per origin at initiation as unit of volume. For this particular condition, the generation time was ln(2)/ 〈λ〉 = 112 minutes and the cell size per origin at replication initiation was *s*_i_ = 0.30 µm^3^.

### Analysis of FtsZ oscillation experiment results

Let us consider a single cell, experiencing a switch in induction, corresponding to a change of steady-state concentration from *c*\* to *c*\*\*. Denoting *S*_ind_ the cell size reached when the switch in induction occurs, by applying Eq. 22 and Eq. 23 from the Supplemental Theory II, we obtain:

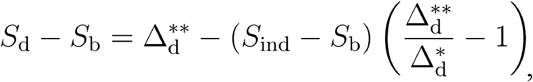

where Δ_d_* = *N*_0_/(2*c*\*) and Δ_d_** = *N*_0_/(2*c*\*\*) are the added size in each induction phase. Assuming exponential elongation of cell size at the rate λ, we may express:

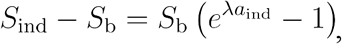

where *a*_ind_ is the age of the cell when the switch in induction occurs. We therefore obtain for the conditional average:

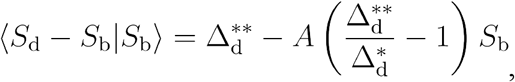

where *A* = 〈e^λ*a*^ - 1〉, and therefore *A* > 0.

### Effect of ClpX on cell size homeostasis

In the presence of ClpX, we consider that division proteins are actively degraded at a rate µ. Denoting *N* the copy number of division proteins, the balanced biosynthesis of division proteins is modified to:

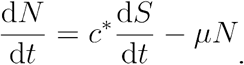

Assuming that the cell volume grows exponentially at the rate λ, the previous ODE can be solved, and one obtains the following relation between copy number and cell volume:

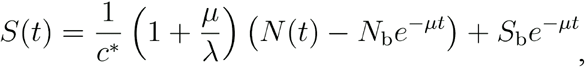

where *N*_b_ = *N*(*t*=0) is the copy number at cell birth and *S*_b_ = *S*(*t*=0) is the cell volume at cell birth. We assume even partitioning of division proteins at division, so that their number at birth is half the threshold: *N*_b_ = *N*_0_ / 2. We can now get some insight on cell size homeostasis by considering the two limiting cases (1) µ ≪λ and (2) µ ≫λ. In case (1), we obtain to order zero in µ/λ that *S*_d_ - *S*_b_ = *N*_0_/(2c*), which is the adder model. On the contrary in case (2), we obtain asymptotically: *S*_d_ = (µ/λ)·*N*_0_/c*, which is the sizer model. In summary, the cell size behavior transitions from the adder model to the sizer model when active degradation of division proteins is introduced.

## QUANTIFICATION AND STATISTICAL ANALYSIS

The error bars in all main figures and supplemental figures represent standard error mean of binned data. In the correlation plots in Figs. 3, 4 and 7, the boundary of shaded area indicate 95% confidence interval of linear fit coefficients assuming the measurement errors are normally distributed and centered at zero. All the fittings were performed in Igor Pro 6 (Wavemetrics, Inc.). The typical sample size of each experiment is about 10^3^ cell. detailed sample size of each experiment is listed in Table S5. In Fig. 7B, the significance of linear correlation (p-value < 0.001) was estimated using Student’s t test in Matlab. In the simulations (see Fig. 2 and Supplemental Theory I), Pearson coefficient was used to quantify both cross-correlations and mother-daughter autocorrelations.

## DATA AND SOFTWARE AVAILABILITY

We share a dataset as the downloadable supplemental data. This dataset includes single-cell physiological data in different steady-state growth conditions. We also would like to share all other data upon request.

## KEY RESOURCES TABLE

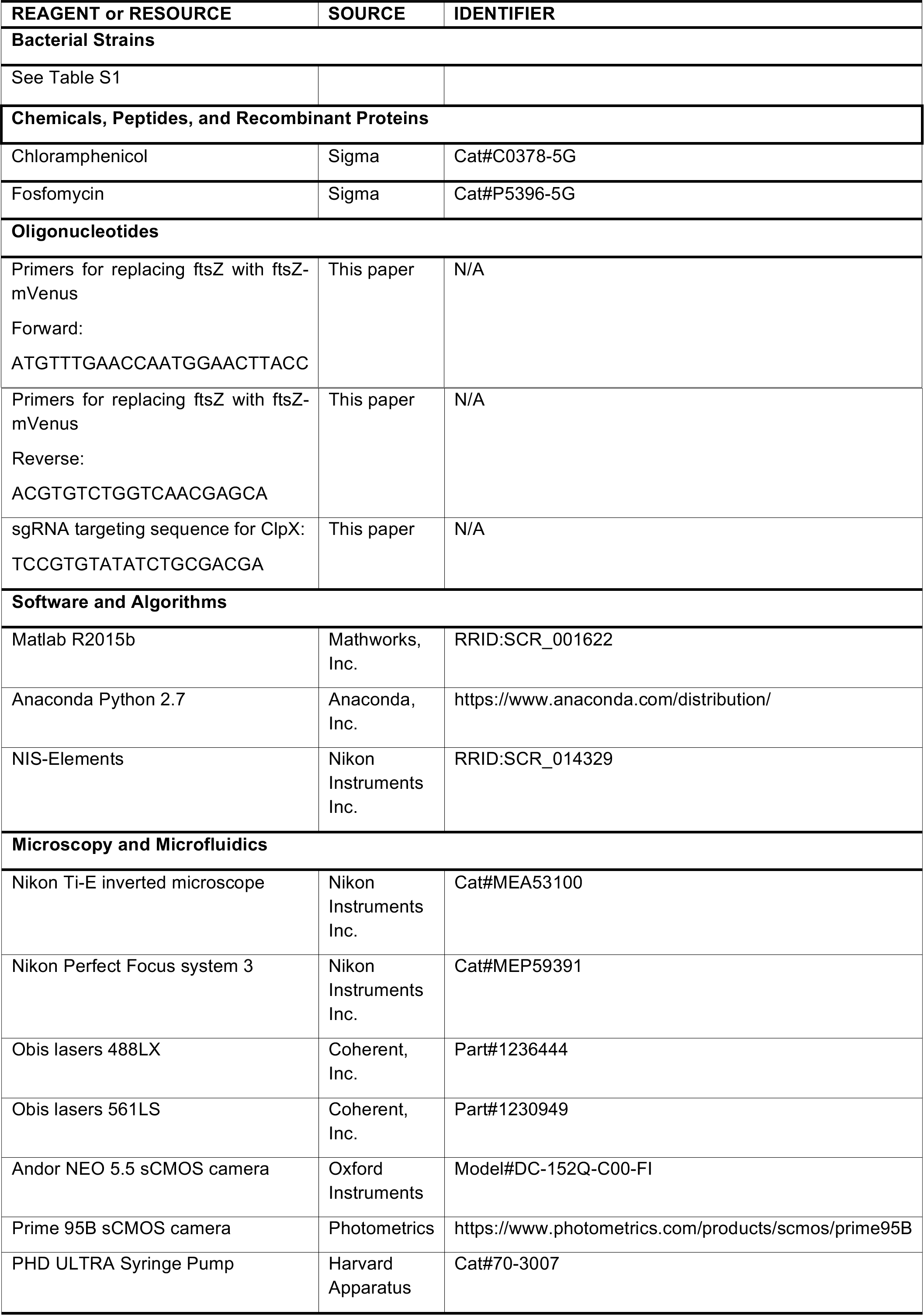

---

**Supplemental Figure S1:**
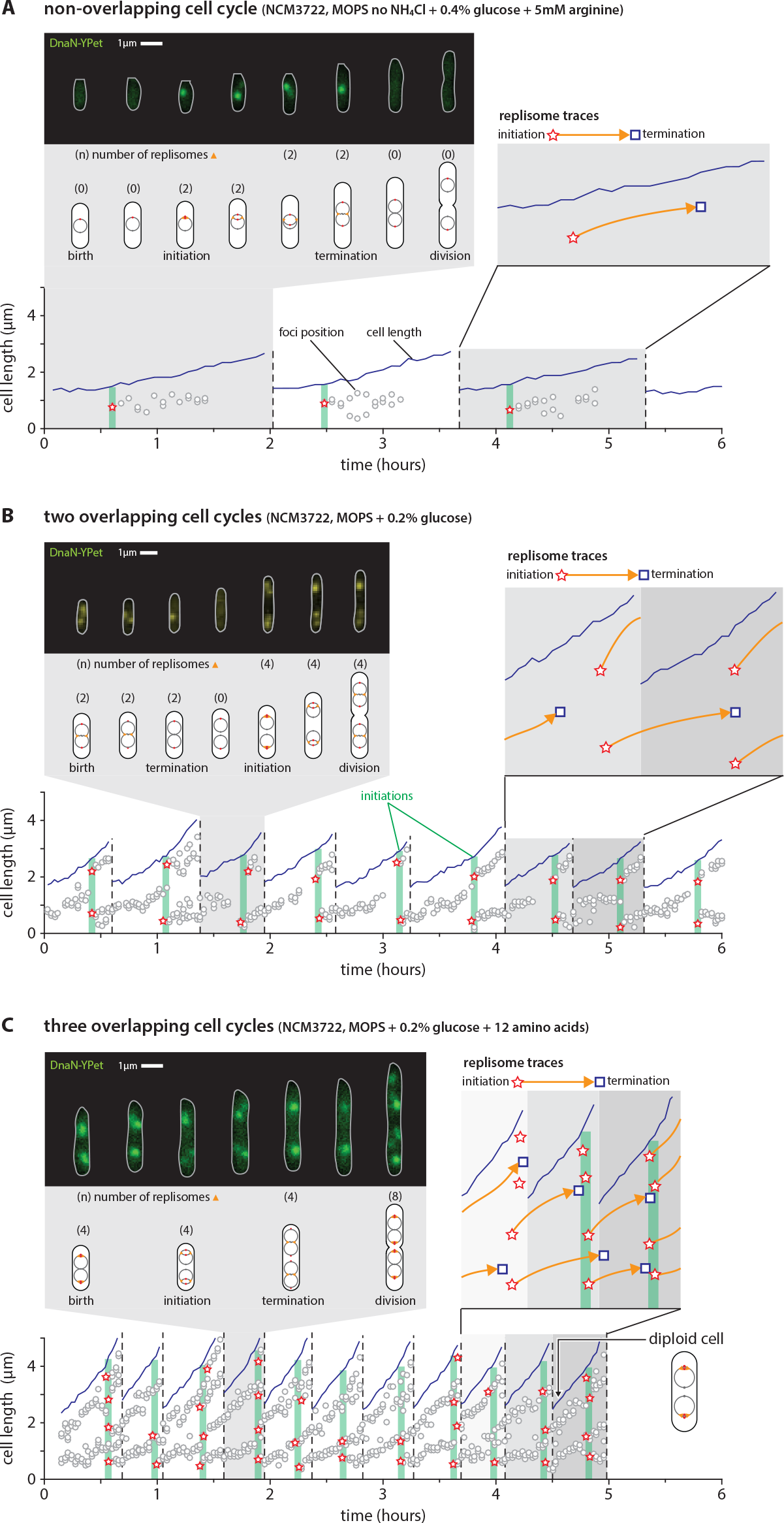
Visualizing and quantifying cell cycle in single bacterial cells in different growth conditions. Related to Figure 1. We measured cell cycle and cell size simultaneously for many consecutive cell division cycles using microfluidic mother machine in different growth conditions. Here **(A)**-**(C)** show the results of three nutrient conditions where the cells are growing with non-, two and three overlapping cell cycles, respectively. The cartoons show the configuration of chromosome replication in one division cycle, similar to that in Fig. 1. The foci positions along the long axis of the cell clearly display the trace of replisomes, making it possible for high-throughput analysis using custom software (see details of image analysis in STAR Methods). Note that, in fast growth conditions such as **(C)**, the termination of replication often finishes before the cell birth, and the daughter cells are born as diploid.

**Supplemental Figure S2:**
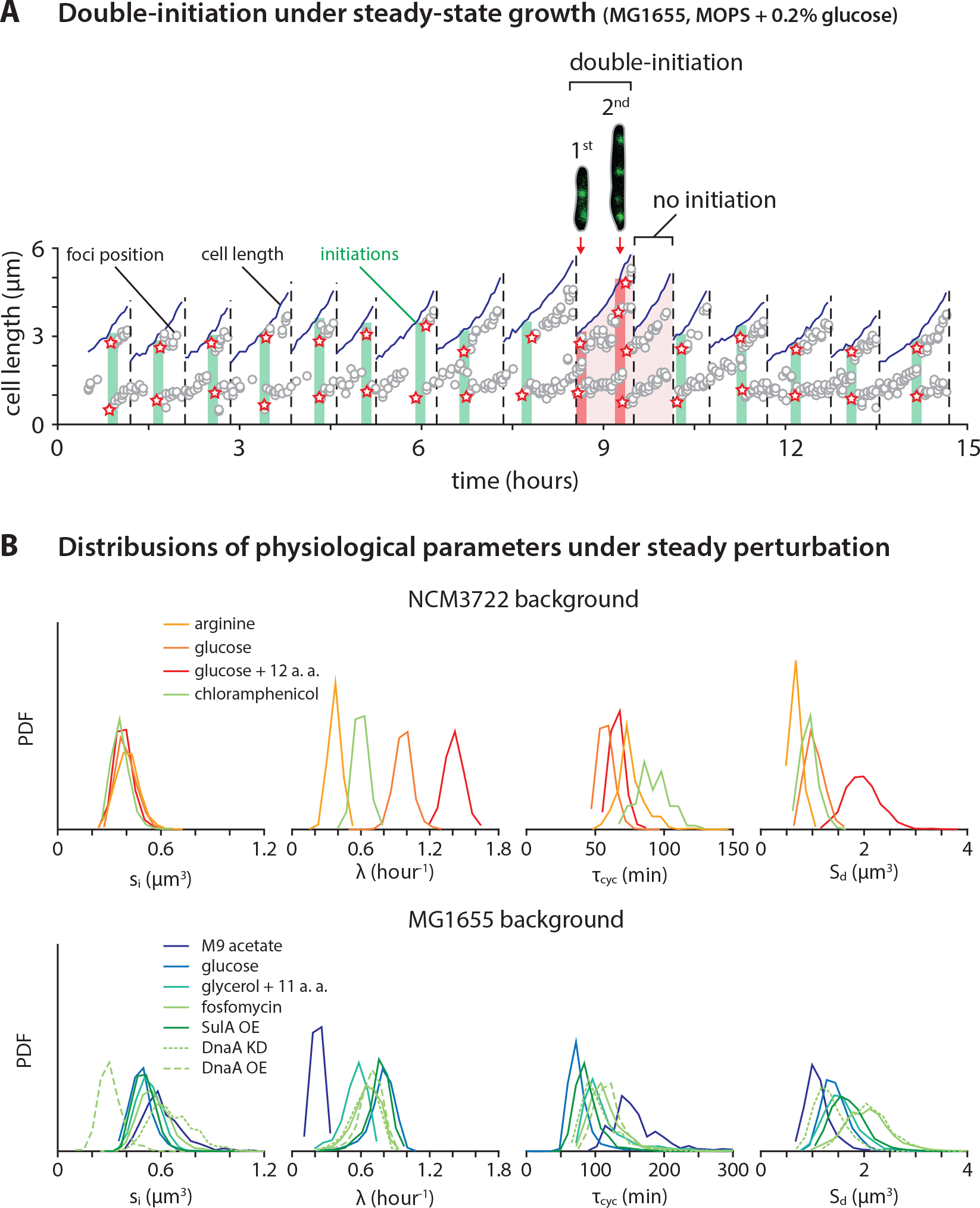
Double-initiation events during steady-state growth due to stochastic cell physiology, and change in major physiological parameters during steady inhibition experiments. Related to Figure 1. **(A)** The decoupling between replication initiation and cell division is evident even from our results of steady-state growth. In the growth condition shown in the figure, cells mostly are with two-overlapping cell cycle, namely, origins duplicate from 2 to 4 at initiation. However, during some division cell cycles, cells fire two rounds of initiations before division, resulting in cells in the next division cycle undergo no initiation. This result disputes the hypothesis that a cell always ensures one-to-one initiation-division correspondence for every division cycle. **(B)** The distributions of major physiological parameters measured from all steady-state experiments as shown in Fig. 1D. For a particular strain background, the initiation size per *ori* largely remains the same, despite the changes in growth rate, cell cycle duration (*τ*_cyc_) or division size. These single-cell results confirm the invariance of initiation mass observed in previous population-level study [1].

**Supplemental Figure S3:**
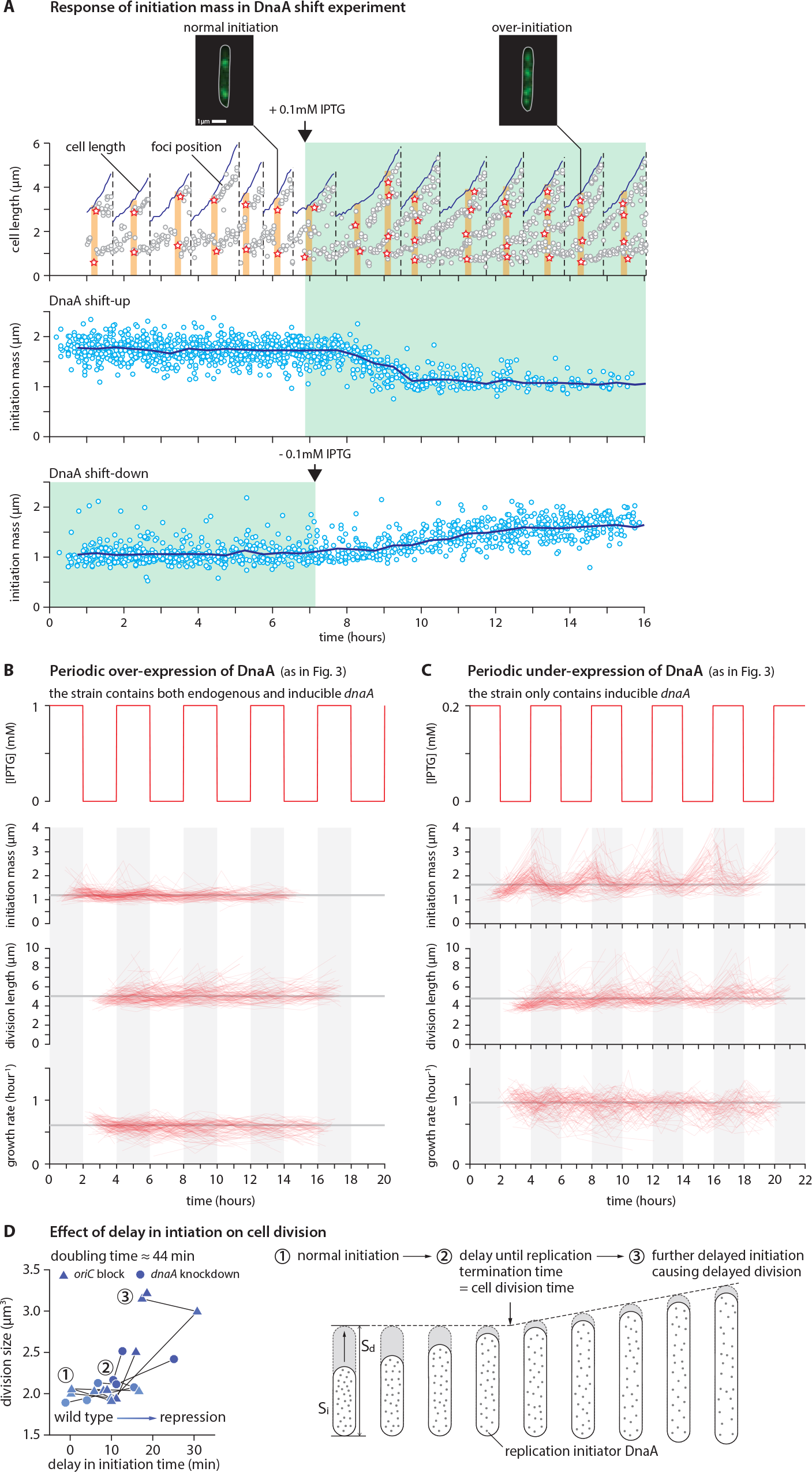
Dependency of initiation mass on DnaA level in the DnaA overexpression experiment, and the full data of dynamic perturbation to DnaA expression. Related to Figure 3. **(A)** Using a strain carrying extra *dnaA* under *P*_lac_ promoter on plasmids (see strain information in STAR Methods and Table S1), we induced the overexpression of DnaA by using 0.1mM IPTG and measured the single-cell cell cycle. In this growth condition (MOPS + 0.4% glycerol + 11 aminos acids), cells over-initiated after induction (DnaA shift-up); the origins duplicated from 2 to 4 before induction and 4 to 8 afterwards. Thus our results show that the initiation mass is dependent on DnaA induction level. The reverse process was observed when 0.1mM IPTG was removed (DnaA shift-down). The response time of both shift-up and shift-down took more than one doubling times (average doubling time ≈ 61min). **(B)** and **(C)** Grey lines indicate the time averages. Each thin trace connects the values of each generation from a single lineage, same for Fig. S4. During the oscillation of DnaA induction, initiation mass changed periodically while growth rate and division size were mostly constant. **(D)** The independent control of initiation and division is seen from our previous population-level data [1]. Using the tunable CRISPR interference system, we delayed the initiation time in a gradual manner. The initiation was delayed by either repressing *dnaA* or blocking *oriC*. When initiation is delayed, the division size remains mostly constant as long as replication termination timing does not exceed the division timing. Thereafter, initiation delay causes an increase in division size.

**Supplemental Figure S4:**
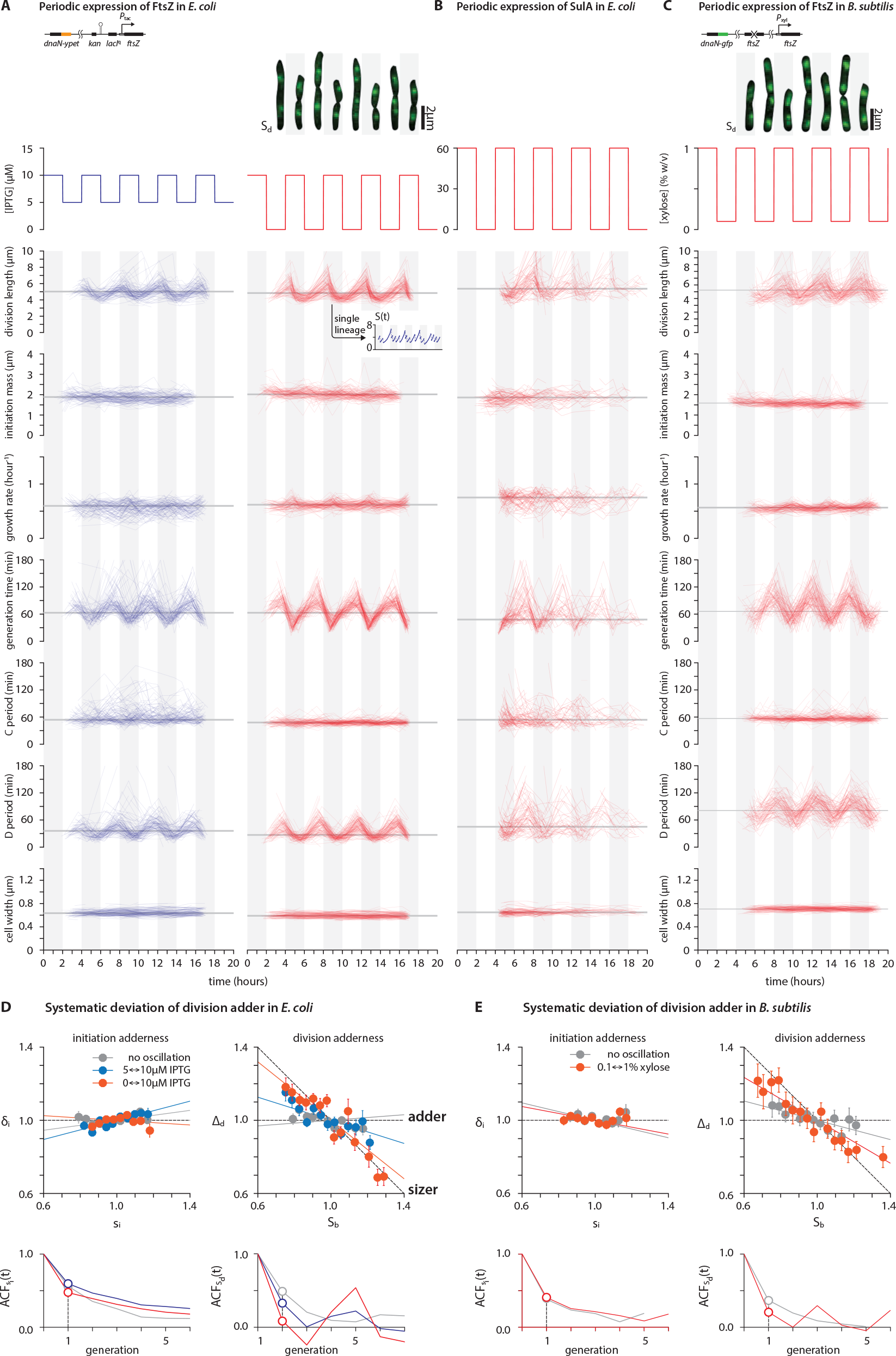
Full data of dynamic perturbation to FtsZ expression in *E. coli* and *B. subtilis*. Related to Figure 4. **(A)** Top left: The schematics of genetic modifications for the inducible system of division protein and fluorescence replisome markers. In *E. coli*, the native promoter of ftsZ was replaced with a *P*_tac_ promoter. Top right: The cell images show oscillations of division size in *E. coli* from a single cell lineage (replisome markers overlaid). Bottom: When FtsZ level was oscillated, division size, generation time and D period were oscillating accordingly. In contrast, growth rate and initiation mass were mostly unchanged. The inset in the right column shows how the division size oscillated seen from a single lineage. **(B)** The periodic expression of SulA in *E. coli* has similar effect on the physiological parameters to that of FtsZ. **(C)** Top: In *B. subtilis*, the endogenous ftsZ was deleted while an alternative copy ftsZ under *P*_xyl_ was inserted at a different loci of the chromosome. The cell images show oscillations of division size in *B. subtilis* from a single cell lineage (replisome markers overlaid). Bottom: When FtsZ level was oscillated, division size, generation time and D period were oscillating accordingly. In contrast, growth rate and initiation mass were mostly unchanged. **(D)** and **(E)** Reprogramming size homeostasis by dynamic modulation of FtsZ in *E. coli* and *B. subtilis* at the oscillation period 4*τ*. IPTG concentrations: blue = 5 *µ*M −10 *µ*M, red = 0 *µ*M −10 *µ*M. Xylose concentration: blue = 0.1% w/v −1 % w/v. In the correlation plots, the variables are normalized by their means and solid lines are model predictions from Supplemental Theory I.E.

**Supplemental Figure S5:**
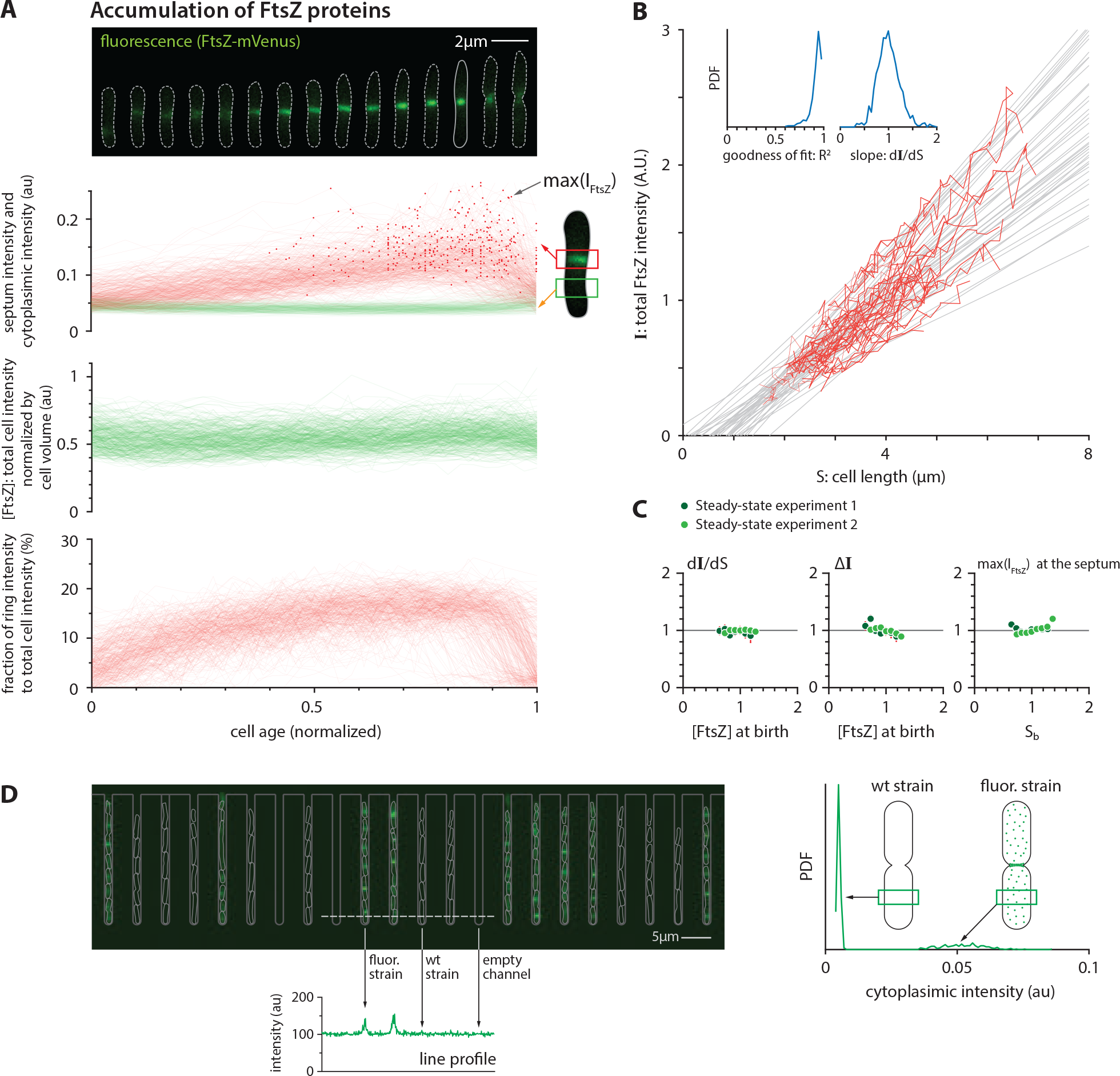
Quantification of FtsZ-mVenus in *E. coli* under steady-state growth. Related to Figure 5. **(A)** Steady accumulation of FtsZ at the Z ring “scaffold” The amount of FtsZ accumulated in the ring was estimated by integrating the total fluorescence intensity within a fixed area enclosing the mid-cell region (septum intensity). The cytoplasmic intensity was measured similarly at the first and third quarter positions. The dark red dot on each trace indicates the max total fluorescence in the Z ring, namely, the peak value of the accumulation trace. The ratio of ring intensity to total cell intensity was calculated as the septum intensity subtracting the cytoplasmic intensity divided by the total fluorescence per cell. This ratio approached a nearly constant value in the first half of the division cycle. During this period, the amount of FtsZ at septum well mirrors the total amount in the cell. **(B)** Single-cell growth traces showing that the total FtsZ intensity per cell is linearly proportional to cell volume. The slope of the linear fit was used to estimate d*I*/d*S* for each single cell. **(C)** Both the production per volume growth estimated by d*I*/d*S* and the threshold estimated by ∆*I* are largely independent of FtsZ concentration at birth. The max total fluorescence in the Z ring is independent of birth size. In the correlation plots, the variables are normalized by their means. **(D)** The autofluorescence of cytoplasm is negligible compared to the fluorescence of cytoplasmic FtsZ-mVenus. Left: To show this, the strain with FtsZ mVenus (SJ1725) and the parental wild type MG1655 strain (SJ81) were co-grew in mother machine. The cytoplasmic autofluorescence of wild type cells is almost same as that from the empty channels. Right: The mean value of cytoplasmic intensity of wild type cell (n=468) is about 9.5% of that of FtsZ-mVenus cells.

**Supplemental Figure S6:**
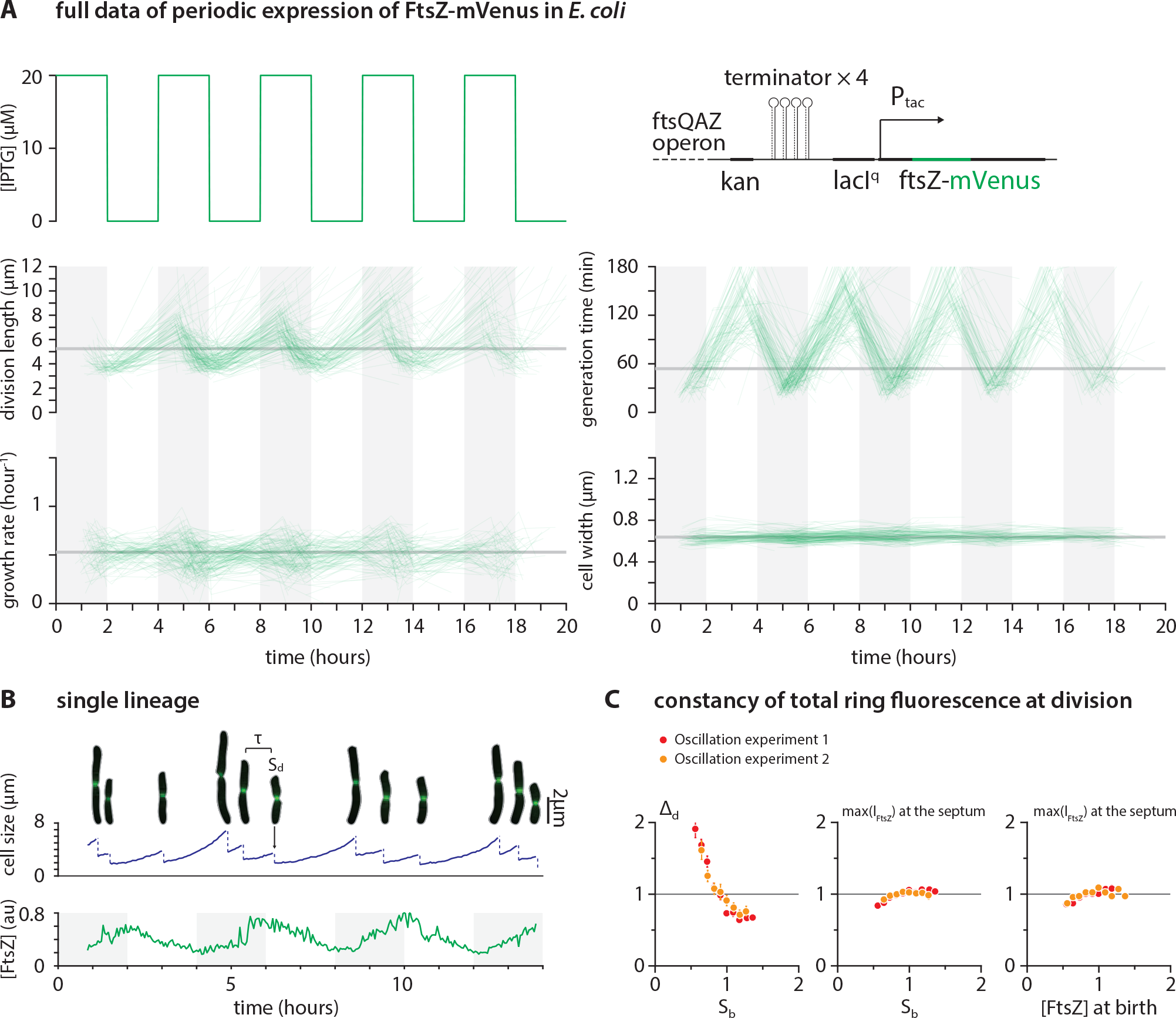
Full data of periodic expression of FtsZ-mVenus in *E. coli*. Related to Figure 6. **(A)** When FtsZ level was oscillated, division size and generation time were oscillating accordingly. In contrast, the growth rate showed very mild change. The illustration at top right shows the design of inducible system for *ftsZ* (see strain information in STAR Methods and table S1). **(B)** The oscillation of division size and FtsZ concentration can be seen from a continuous single lineage. **(C)** The maximal total fluorescence of Z-ring is largely independent of FtsZ concentration and cell size at birth. In the correlation plots, the variables are normalized by their means.

**Supplemental Figure S7:**
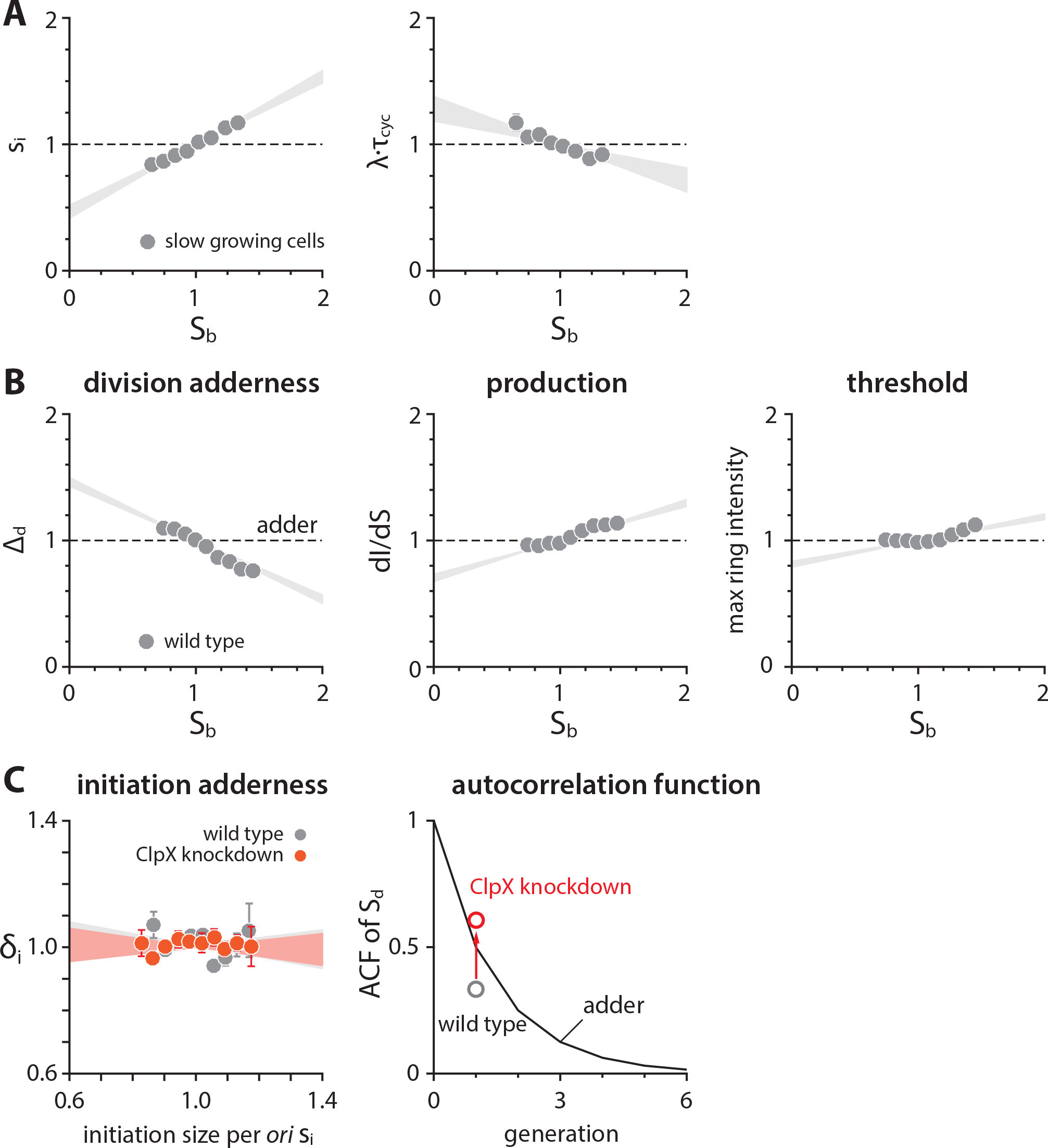
Cell-size homeostasis and production of FtsZ-mVenus in slow growth conditions. Related to Figure 7. **(A)** Both *s*_*i*_ and *λ ⋅ τ*_cyc_ show correlation with birth size. **(B)** In slow growth condition, the cell-size homeostasis deviates from adder and the production per volume growth d*I*/d*S* is also no longer independent of birth size. In the correlation plots, the variables are normalized by their means. **(C)** In slow-growing *E. coli* cells, repressing *clpX* expression via tCRISPRi restored the division adder, while initiation adder was invariant. Each shaded area represents the 95% confidence interval of linear fit to the respective raw scatter plot.

**Supplemental Table S1:**
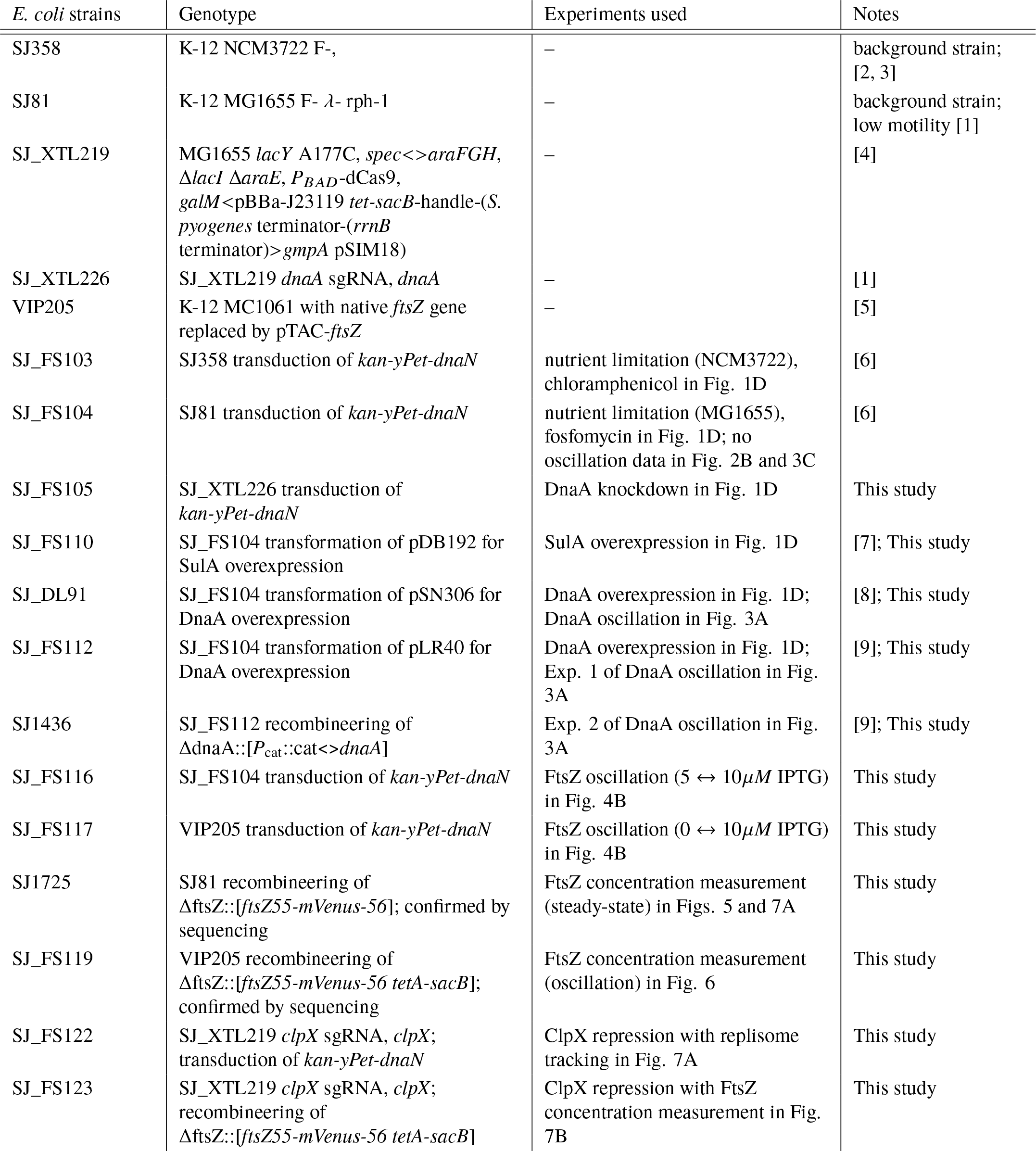
Strain Information of *E. coli*.

| *E. coli* strains | Genotype | Experiments used | Notes |
| --- | --- | --- | --- |
| SJ358 | K-12 NCM3722 F-, | – | background strain; [2, 3] |
| SJ81 | K-12 MG1655 F- λ-rph-1 | – | background strain; low motility [1] |
| SJ_XTL219 | MG1655 *lacY* A177C, *spec*<>*araFGH*, Δ*lacI* Δ*araE*, *P*<sub>BAD</sub>-dCas9, *galM*<pBBa-J23119 *tet-sacB*-handle-(*S. pyogenes* terminator-(*rrnB* terminator)>*gmpA* pSIM18) | – | [4] |
| SJ_XTL226 | SJ_XTL219 *dnaA* sgRNA, *dnaA* | – | [1] |
| VIP205 | K-12 MC1061 with native *ftsZ* gene replaced by pTAC-*ftsZ* | – | [5] |
| SJ_FS103 | SJ358 transduction of *kan-yPet-dnaN* | nutrient limitation (NCM3722), chloramphenicol in Fig. 1D | [6] |
| SJ_FS104 | SJ81 transduction of *kan-yPet-dnaN* | nutrient limitation (MG1655), fosfomycin in Fig. 1D; no oscillation data in Fig. 2B and 3C | [6] |
| SJ_FS105 | SJ_XTL226 transduction of *kan-yPet-dnaN* | DnaA knockdown in Fig. 1D | This study |
| SJ_FS110 | SJ_FS104 transformation of pDB192 for SulA overexpression | SulA overexpression in Fig. 1D | [7]; This study |
| SJ_DL91 | SJ_FS104 transformation of pSN306 for DnaA overexpression | DnaA overexpression in Fig. 1D; DnaA oscillation in Fig. 3A | [8]; This study |
| SJ_FS112 | SJ_FS104 transformation of pLR40 for DnaA overexpression | DnaA overexpression in Fig. 1D; Exp. 1 of DnaA oscillation in Fig. 3A | [9]; This study |
| SJ1436 | SJ_FS112 recombineering of Δ*dnaA*::[*P*<sub>cat</sub>::cat<>*dnaA*] | Exp. 2 of DnaA oscillation in Fig. 3A | [9]; This study |
| SJ_FS116 | SJ_FS104 transduction of *kan-yPet-dnaN* | FtsZ oscillation (5 ↔ 10 μM IPTG) in Fig. 4B | This study |
| SJ_FS117 | VIP205 transduction of *kan-yPet-dnaN* | FtsZ oscillation (0 ↔ 10 μM IPTG) in Fig. 4B | This study |
| SJ1725 | SJ81 recombineering of Δ*ftsZ*::[*ftsZ55-mVenus-56*]; confirmed by sequencing | FtsZ concentration measurement (steady-state) in Figs. 5 and 7A | This study |
| SJ_FS119 | VIP205 recombineering of Δ*ftsZ*::[*ftsZ55-mVenus-56 tetA-sacB*]; confirmed by sequencing | FtsZ concentration measurement (oscillation) in Fig. 6 | This study |
| SJ_FS122 | SJ_XTL219 *clpX* sgRNA, *clpX*; transduction of *kan-yPet-dnaN* | ClpX repression with replisome tracking in Fig. 7A | This study |
| SJ_FS123 | SJ_XTL219 *clpX* sgRNA, *clpX*; recombineering of Δ*ftsZ*::[*ftsZ55-mVenus-56 tetA-sacB*] | ClpX repression with FtsZ concentration measurement in Fig. 7B | This study |

**Supplemental Table S2:** Strain Information of *B. subtilis*.

| *B. subtilis* strains | Genotype | Experiments used | Notes |
| --- | --- | --- | --- |
| PAW885 | JH642, *dnaN-gfp-spec* | – | [10, 11] |
| SJ_BS29 | PAW885 transformation of *motAB::Tn917* | – | This study |
| SEV645 | SJ_BS29 transformation of *ftsZ::cm* and *thrC::Pxyl-ftsZ* | steady state and FtsZ oscillation in Fig. 4B | This study |

**Supplemental Table S3:**
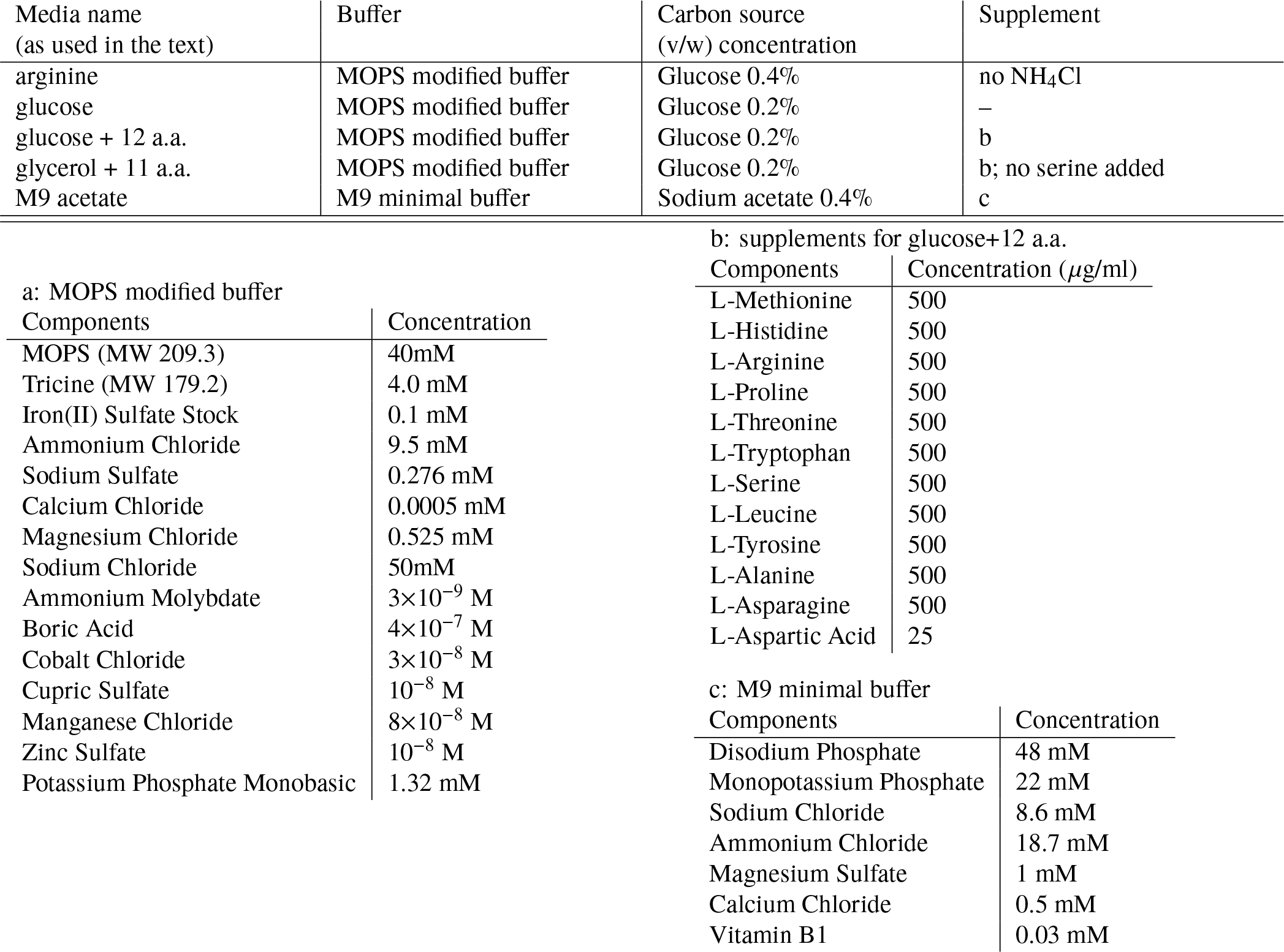
List of growth media, carbon sources and supplements for *E. coli*.

**Supplemental Table S4:**
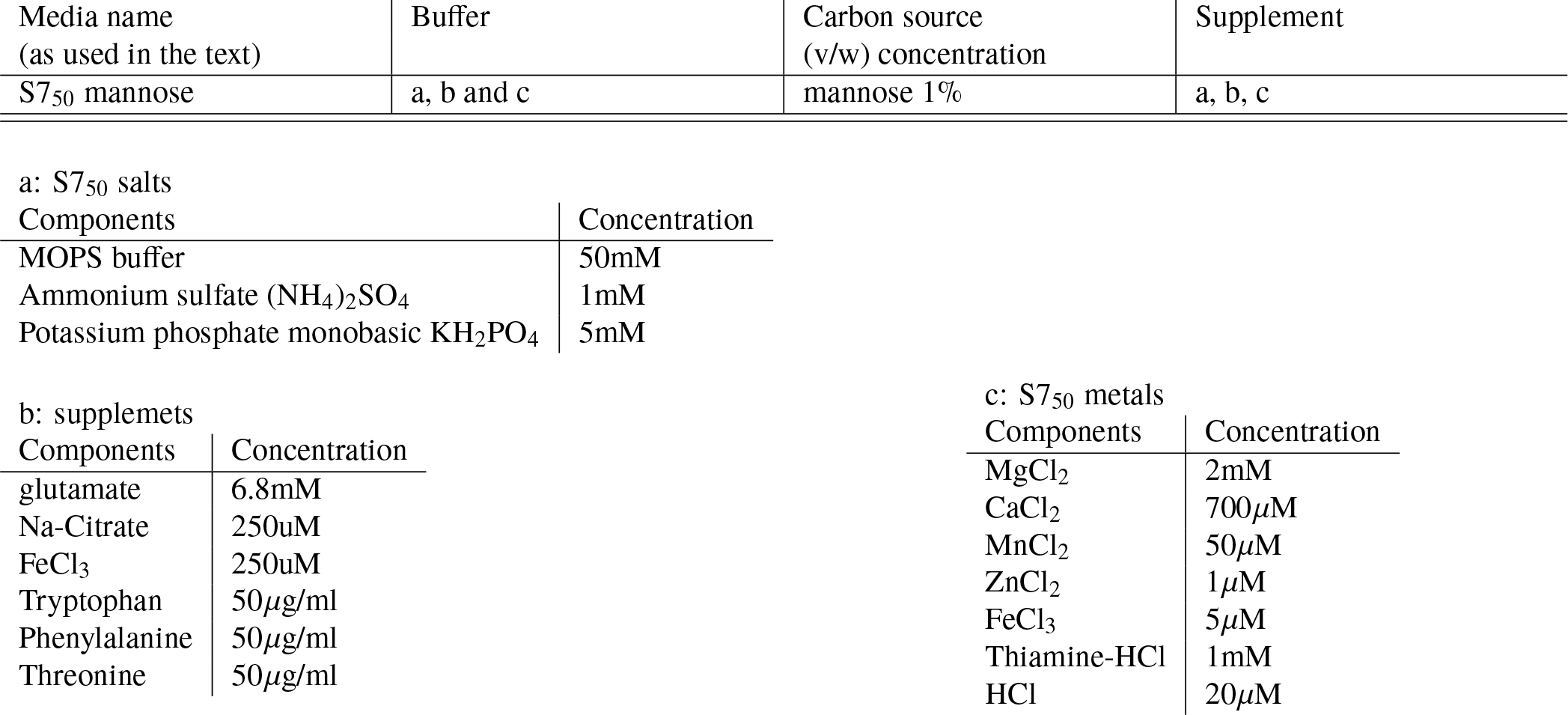
List of growth media, carbon sources and supplements for *B. subtilis*.

| Media name<br>(as used in the text) | Buffer | Carbon source<br>(v/w) concentration | Supplement |
| --- | --- | --- | --- |
| S7<sub>50</sub> mannose | a, b and c | mannose 1% | a, b, c |

a: S7<sub>50</sub> salts
| Components | Concentration |
| --- | --- |
| MOPS buffer | 50mM |
| Ammonium sulfate (NH<sub>4</sub>)<sub>2</sub>SO<sub>4</sub> | 1mM |
| Potassium phosphate monobasic KH<sub>2</sub>PO<sub>4</sub> | 5mM |

b: supplemets
| Components | Concentration |
| --- | --- |
| glutamate | 6.8mM |
| Na-Citrate | 250uM |
| FeCl<sub>3</sub> | 250uM |
| Tryptophan | 50μg/ml |
| Phenylalanine | 50μg/ml |
| Threonine | 50μg/ml |

| Components | Concentration |
| --- | --- |
| MgCl<sub>2</sub> | 2mM |
| CaCl<sub>2</sub> | 700μM |
| MnCl<sub>2</sub> | 50μM |
| ZnCl<sub>2</sub> | 1μM |
| FeCl<sub>3</sub> | 5μM |
| Thiamine-HCl | 1mM |
| HCl | 20μM |

**Supplemental Table S5:**
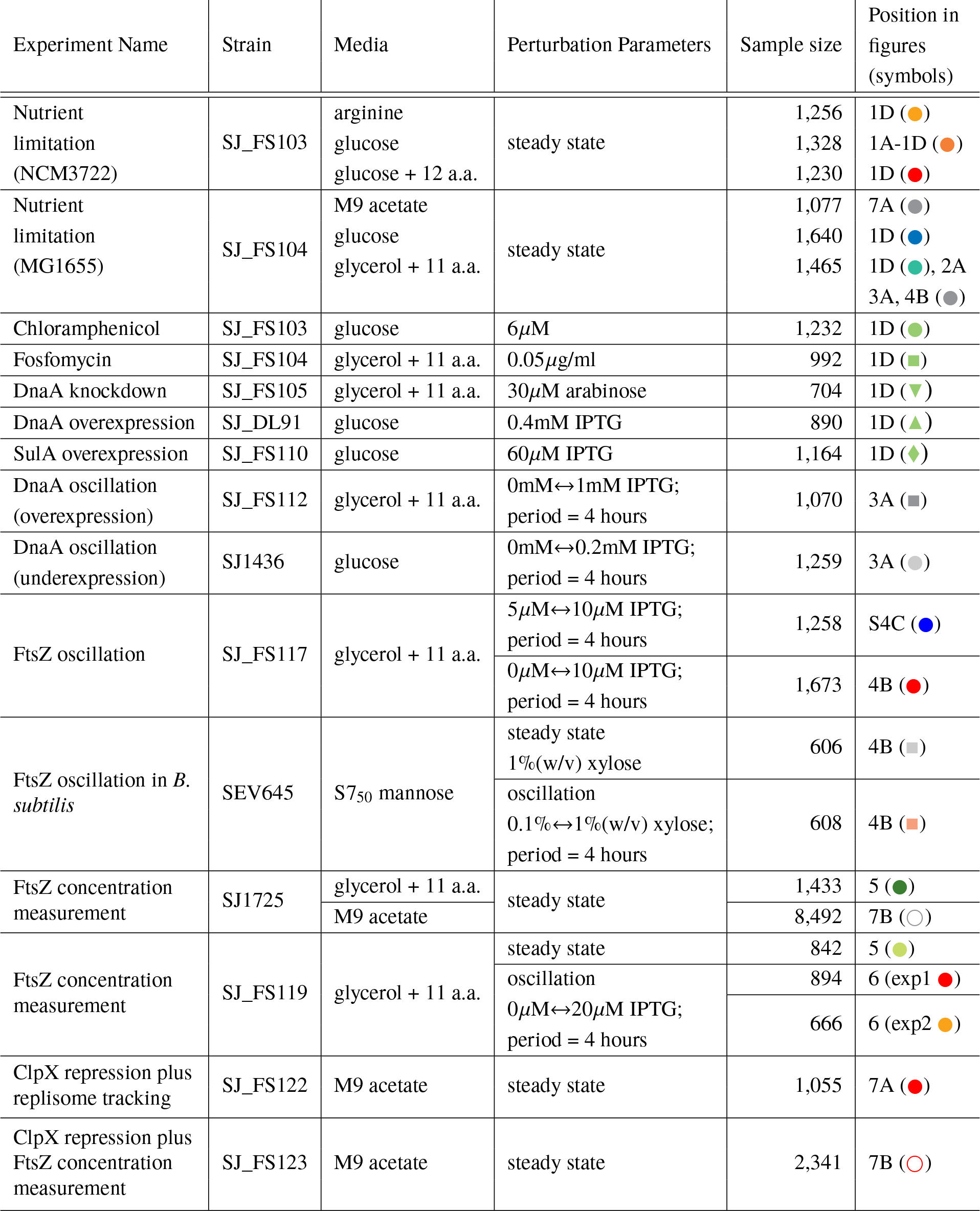
Experimental conditions and sample size. The sample size represents the number of single cell division cycles measured from each mother machine experiment. The symbols in the rightmost columns are the same as those in the corresponding main figures. ‘↔’ indicates that the oscillation experiment was conducted between the two concentrations of inducer on both sides.

| Experiment Name | Strain | Media | Perturbation Parameters | Sample size | Position in figures (symbols) |
| --- | --- | --- | --- | --- | --- |
| Nutrient limitation (NCM3722) | SJ_FS103 | arginine<br>glucose<br>glucose + 12 a.a. | steady state | 1,256<br>1,328<br>1,230 | 1D (●)<br>1A-1D (●)<br>1D (●) |
| Nutrient limitation (MG1655) | SJ_FS104 | M9 acetate<br>glucose<br>glycerol + 11 a.a. | steady state | 1,077<br>1,640<br>1,465 | 7A (●)<br>1D (●)<br>1D (●), 2A<br>3A, 4B (●) |
| Chloramphenicol | SJ_FS103 | glucose | 6μM | 1,232 | 1D (●) |
| Fosfomycin | SJ_FS104 | glycerol + 11 a.a. | 0.05μg/ml | 992 | 1D (■) |
| DnaA knockdown | SJ_FS105 | glycerol + 11 a.a. | 30μM arabinose | 704 | 1D (▼) |
| DnaA overexpression | SJ_DL91 | glucose | 0.4mM IPTG | 890 | 1D (▲) |
| SulA overexpression | SJ_FS110 | glucose | 60μM IPTG | 1,164 | 1D (◆) |
| DnaA oscillation (overexpression) | SJ_FS112 | glycerol + 11 a.a. | 0mM↔1mM IPTG;<br>period = 4 hours | 1,070 | 3A (■) |
| DnaA oscillation (underexpression) | SJ1436 | glucose | 0mM↔0.2mM IPTG;<br>period = 4 hours | 1,259 | 3A (●) |
| FtsZ oscillation | SJ_FS117 | glycerol + 11 a.a. | 5μM↔10μM IPTG;<br>period = 4 hours | 1,258 | S4C (●) |
|  |  |  | 0μM↔10μM IPTG;<br>period = 4 hours | 1,673 | 4B (●) |
| FtsZ oscillation in *B. subtilis* | SEV645 | S7<sub>50</sub> mannose | steady state<br>1%(w/v) xylose | 606 | 4B (■) |
|  |  |  | oscillation<br>0.1%↔1%(w/v) xylose;<br>period = 4 hours | 608 | 4B (■) |
| FtsZ concentration measurement | SJ1725 | glycerol + 11 a.a. | steady state | 1,433 | 5 (●) |
|  |  | M9 acetate |  | 8,492 | 7B (○) |
| FtsZ concentration measurement | SJ_FS119 | glycerol + 11 a.a. | steady state | 842 | 5 (●) |
|  |  |  | oscillation<br>0μM↔20μM IPTG;<br>period = 4 hours | 894 | 6 (exp1 ●) |
|  |  |  |  | 666 | 6 (exp2 ●) |
| ClpX repression plus replisome tracking | SJ_FS122 | M9 acetate | steady state | 1,055 | 7A (●) |
| ClpX repression plus FtsZ concentration measurement | SJ_FS123 | M9 acetate | steady state | 2,341 | 7B (○) |

## Supplemental theory for “The mechanistic origin of cell-size homeostasis in bacteria”

### I. SINGLE-CELL STOCHASTIC HELMSTETTER-COOPER MODEL

#### A. Deterministic Helmstetter-Cooper model

Here we model the growth of individual *Escherichia coli* cells. Based on experimental measurements, we posit that a single-cell of size *S* elongates exponentially [12, 13, 14]:

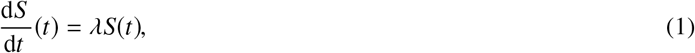

where *λ* is the growth rate. For rod-shaped bacteria such as *E. coli* the width is nearly constant, so the size means either the length or volume of an individual cell (to a proportionality constant). In order to determine the division timing, we adopt the Helmstetter-Cooper model, which couples the replication of the chromosome to the cell division (see Suppl. Fig. T1). This model proposes that cell division occurs after a prescribed time has elapsed since replication initiation. This time is the duration of the cell cycle, denoted *τ*_cyc_ [15, 14]. It comprises the time required to fully replicate the chromosome, known as the C-period, and the time following replication termination until division, known as the D-period. Hence *τ*_cyc_ = *C* + *D*. In other words, an initiation event commits the cell to division after the duration of one cell cycle. In bacteria, multiple cell cycles can overlap. This occurs when the cell cycle duration is larger than the generation time: *τ*_cyc_ > *τ*. In order to maintain its DNA content, a cell still needs to initiate chromosome replication once per generation (see Suppl. Fig. T1). At this stage, the problem of division timing is thus transfered to the problem of initiation timing. This is solved by considering Donachie’s conjecture [16], which is that a new round of replication initiates at a fixed size per number of origins of replication (*oriC*), denoted *s*_i_.

In summary, a model for the growth of a single cell is completely parametrized by the following “physiological variables”:

- the growth rate *λ*;
- the cell cycle duration *τ*_cyc_;
- the initiation size per *oriC s*_i_.

This model predicts that the cell at division is given by:

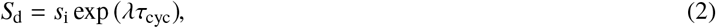

which was indeed verified experimentally at the population level [1]. Single-cell measurement of the initiation timing revealed that the initiation size is indeed tightly regulated. In fact, *s*_i_ is the physiological variable with the narrowest distribution (Suppl. Figs. T3 to T6).

**Supplemental Figure T1:**
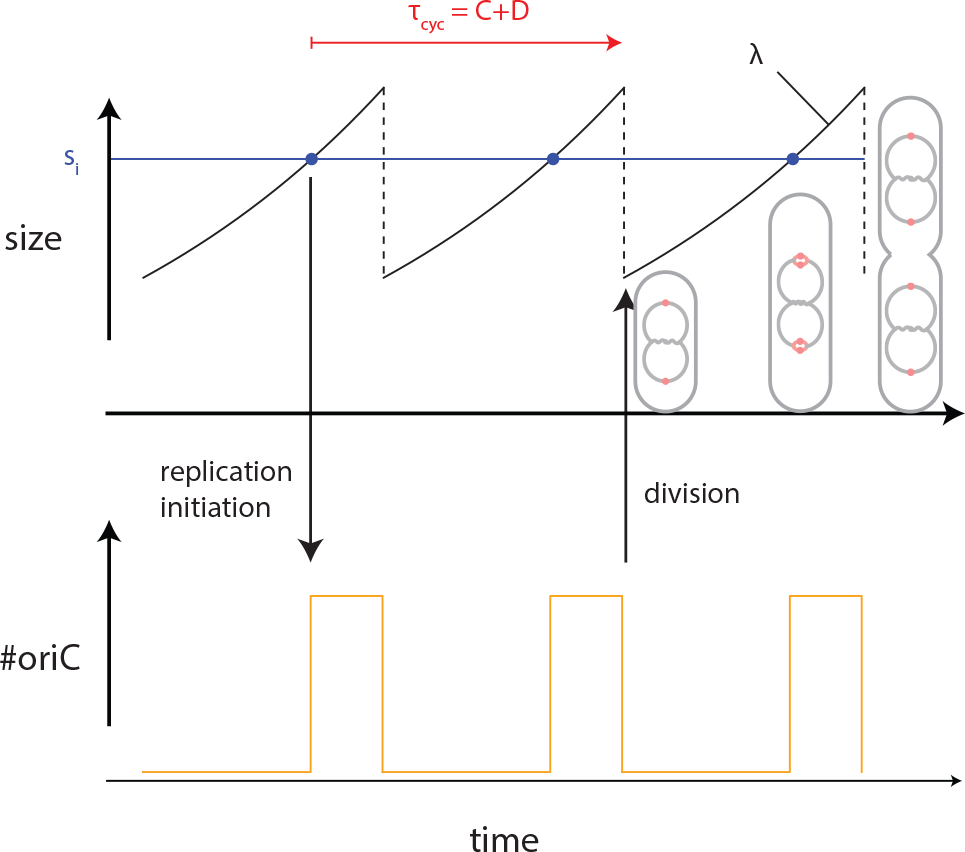
Helmstetter-Cooper model. In bacteria, several rounds of chromosome replication can overlap when the duration of the cell cycle τ_cyc_, including both C- and D-periods, is larger than the generation time.

#### B. Stochastic Helmstetter-Cooper model

In order to account for the experimental fluctuations in the physiological variables between individual cells, we treat them as stochastic variables. That is to say, at cell birth, the growth rate, the cell cycle duration and the unit cell are drawn from independent Gaussian distributions (Suppl. Fig. T2). Once the physiological variables are determined, a single-cell follows the deterministic growth as described in the previous section.

**Supplemental Figure T2:**
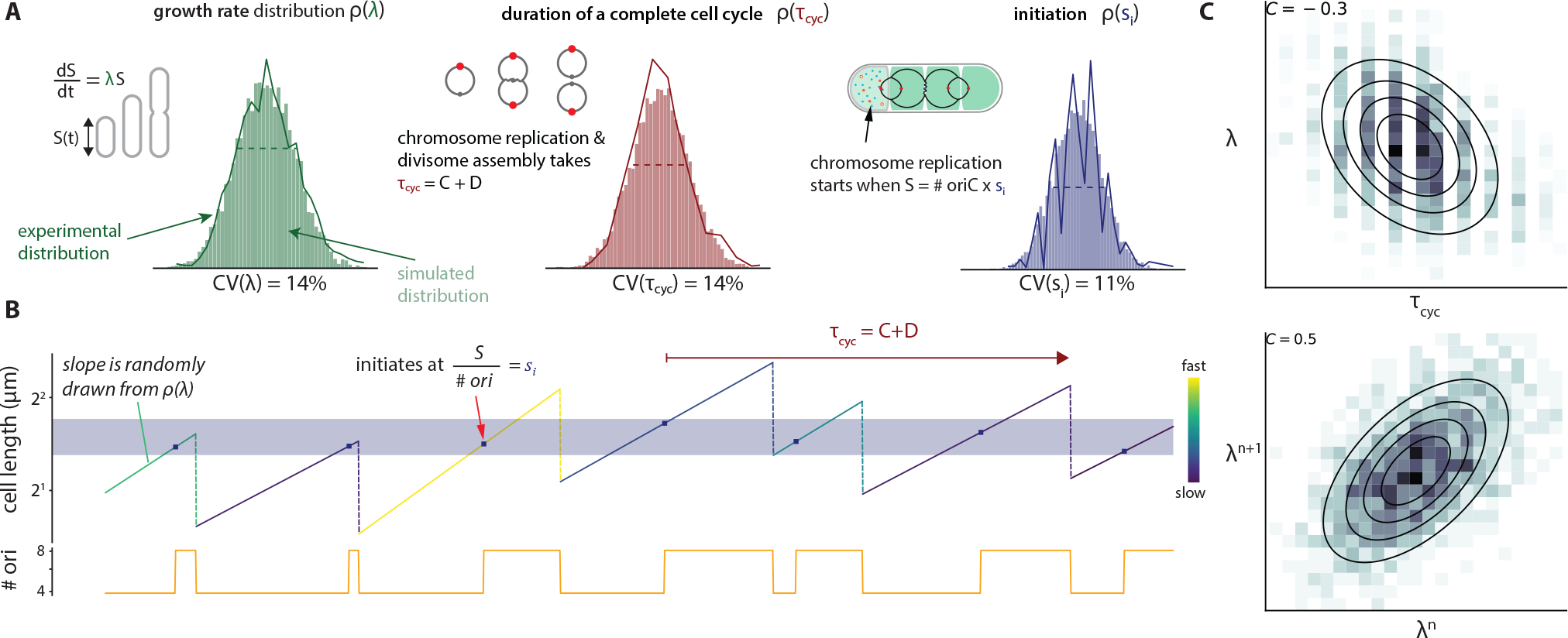
Stochastic Helmstetter-Cooper model. **(A)** At each new generation, the physiological variables are drawn from a multivariate Gaussian distribution with means and variances matching the experimental values. **(B)** In this example, cell division is coupled to an initiation event happening in the grandmother cell: there is three overlapping cell cycles. **(C)** Examples of negative cross-correlation between λ and τ_cyc_ and of positive mother/daughter correlation for λ.

In reality, we should expect that values taken by the physiological variables are not independent from each other. For example, the growth rate and the cell cycle duration are anti-correlated [14]. Another example comes from the observation that cells growing faster than the average growth rate also tend to produce fast growing daughter cells. We call the former type of correlation “cross-correlation” and the latter ones “autocorrelation”. In order to take into account those correlations, we modify the way physiological variables are determined at cell birth. Let us denote the physiological state of a single cell by the three-dimensional vector 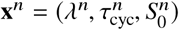. At cell birth, the physiological state of cell *n* + 1 is determined according to the rule:

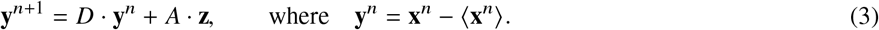

Here, *D* = Diag(α_1_, α_2_, α_3_) is a diagonal matrix enforcing mother/daughter correlations, *A* is a real symmetric matrix enforcing the cross-correlations (to be determined below) and **z** is a vector of three independent unit Gaussian variables.

Eq. (3) defines a discrete stochastic process. Being a sum of Gaussian random vectors, **y**^*n*^ has a (multivariate) Gaussian distribution. Furthermore, it can be shown that it converges toward the Gaussian distribution:

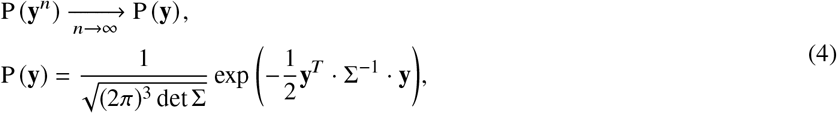

whose covariance matrix Σ = [σ_*ij*_]_*i,j*_ has the elements:

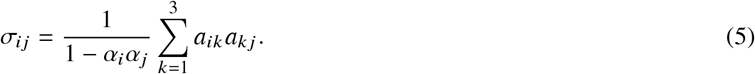

The previous relation can be inverted to express the matrix *A* as a function of the covariance matrix of the limiting distribution. We obtain:

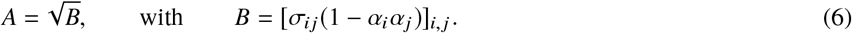

In short, with an appropriate choice of the matrix *A*, the stochastic process in Eq. (3) will sample random physiological vectors distributed according to a Gaussian distribution with the desired covariance matrix Σ, in agreement with experimental measurements. For the stochastic process in Eq. (3), the variances of the physiological variables are given by:

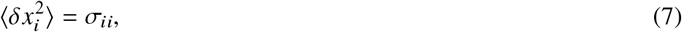

and the cross-correlations between the physiological variables are expressed as:

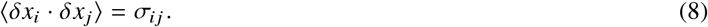

Note that the Pearson correlation coefficients between the physiological variables are expressed as:

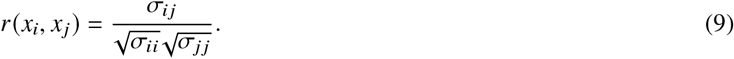

Concerning mother/daughter correlations, it can be shown that:

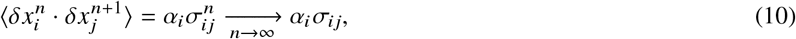

with the corresponding Pearson correlations:

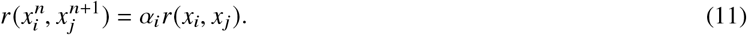

In particular, the autocorrelation Pearson coefficients are:

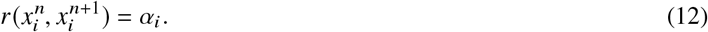

In summary, we draw the physiological variables at each new generation according to Eq. (3). This stochastic process is parametrized by the experimental values measured for the means, variances, cross-correlations and autocorrelations of the three physiological variables.

#### C. Implementation

The implementation used to generate a lineage of cells according to the stochastic Helmstetter-Cooper model is described in Algorithms 1 to 3. The simulations rely on the following input: (i) the means 〈λ〉, 〈τ_cyc_〉 and 〈*s*_i_〉; (ii) the variances 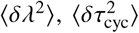 and 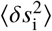; (iii) the Pearson cross-correlation coefficients *r*_*ij*_: = *r*(*x*_*i*_, *x*_*j*_) defined above; and (iv) the Pearson autocorrelation coefficients α_*i*_ defined above. These inputs can be measured experimentally and were used to set the joint-probability distribution of the physiological variables. Unless specified otherwise, we generated a single lineage of 10 000 cells in one simulation.

#### D. Comparison with experiments

We performed simulations according to the stochastic Helmstetter-Cooper for several experimental conditions (Suppl. Figs. T3 to T6). Overall, the agreement between experiments and simulations was good. For non-overlapping cell cycles conditions, most of the distributions for the cell size at birth *S*_b_, the cell size at division *S*_d_, the generation time *τ* and the added size between divisions ∆_d_ = *S*_d_ − *S*_b_ were well reproduced. This observation was less true for overlapping cell cycles. For example in the latter case, the distribution of division size predicted by stochastic Helmstetter-Cooper model was systematically broader than the experimental one. In general, we found that the predicted correlations between variables were in good agreement with the experimental measurements.

The simulations of the stochastic Helmstetter-Cooper model reproduced well the experimental behavior for cell size homeostasis, namely the adder behavior. In Suppl. Fig. T8, we illustrate how the adder behavior converges toward the experimental value when cross-correlations and autocorrelations are added to the model. Clearly, in the absence of cross-correlations and/or autocorrelations, the behavior deviates from the experimental measurements. This suggests that such correlations are essential for the Helmstetter-Cooper model to reproduce the experimental cell size homeostasis behavior.

#### E. Co-regulation hypothesis of chromosome replication and division

The Helmstetter-Cooper is often interpreted as to impose a fixed cell cycle duration, τ_cyc_. If in addition the growth rate is fixed, Eq. (2) implies that the cell size at division *S*_d_ is proportional to the cell size per origin of replication *s*_i_ at the initiation event that led to the division. The response of division sizes is then linked to the response of initiation size to perturbations. In particular, their Pearson autocorrelation coefficients are equal:

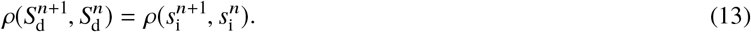

**Supplemental Figure T3:**
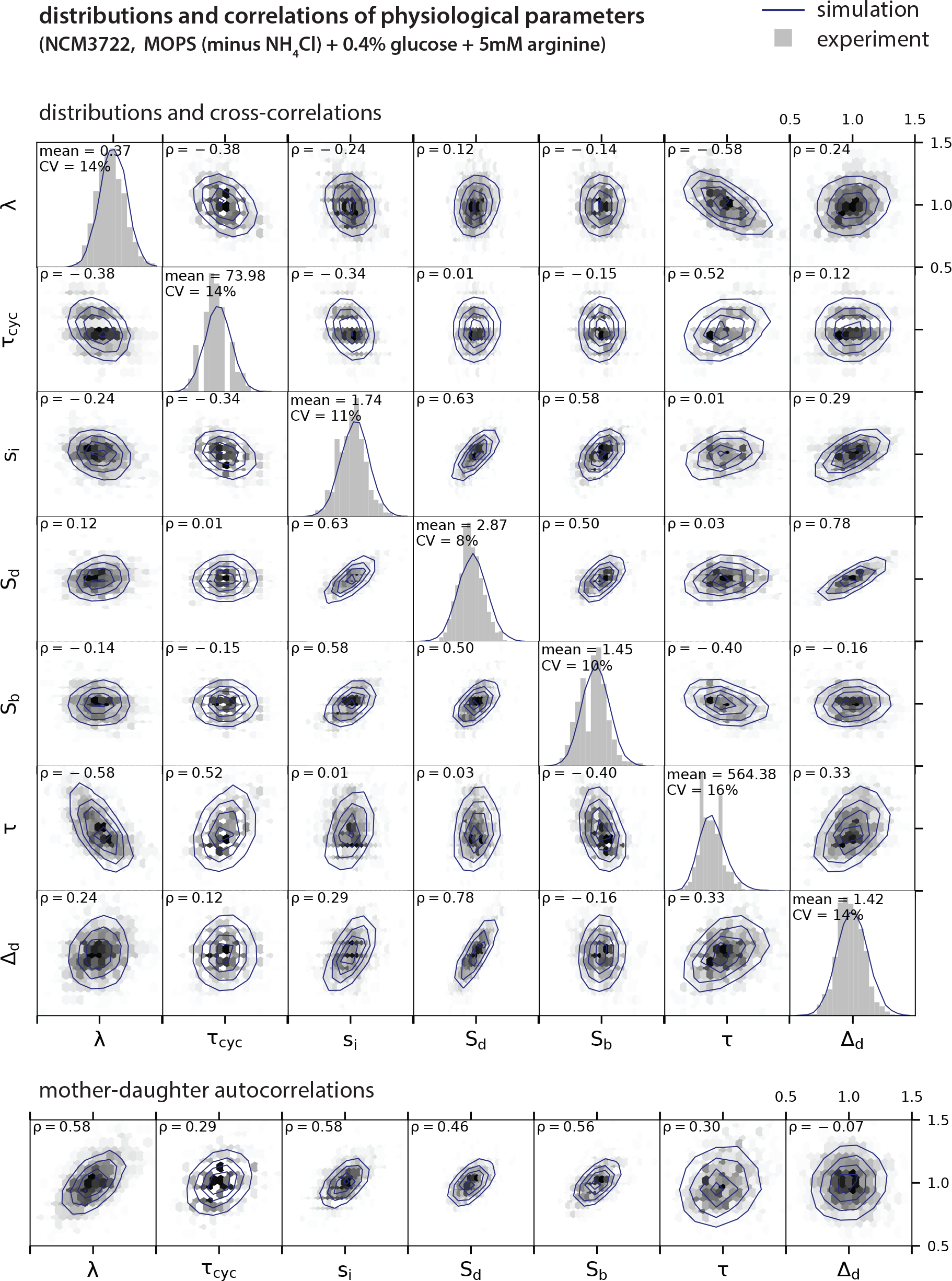
Comparison of experimental measurements with simulations of the Helmstetter-Cooper model. NCM3722 strain with no overlapping cell cycles as in Fig. 1D.

**Supplemental Figure T4:**
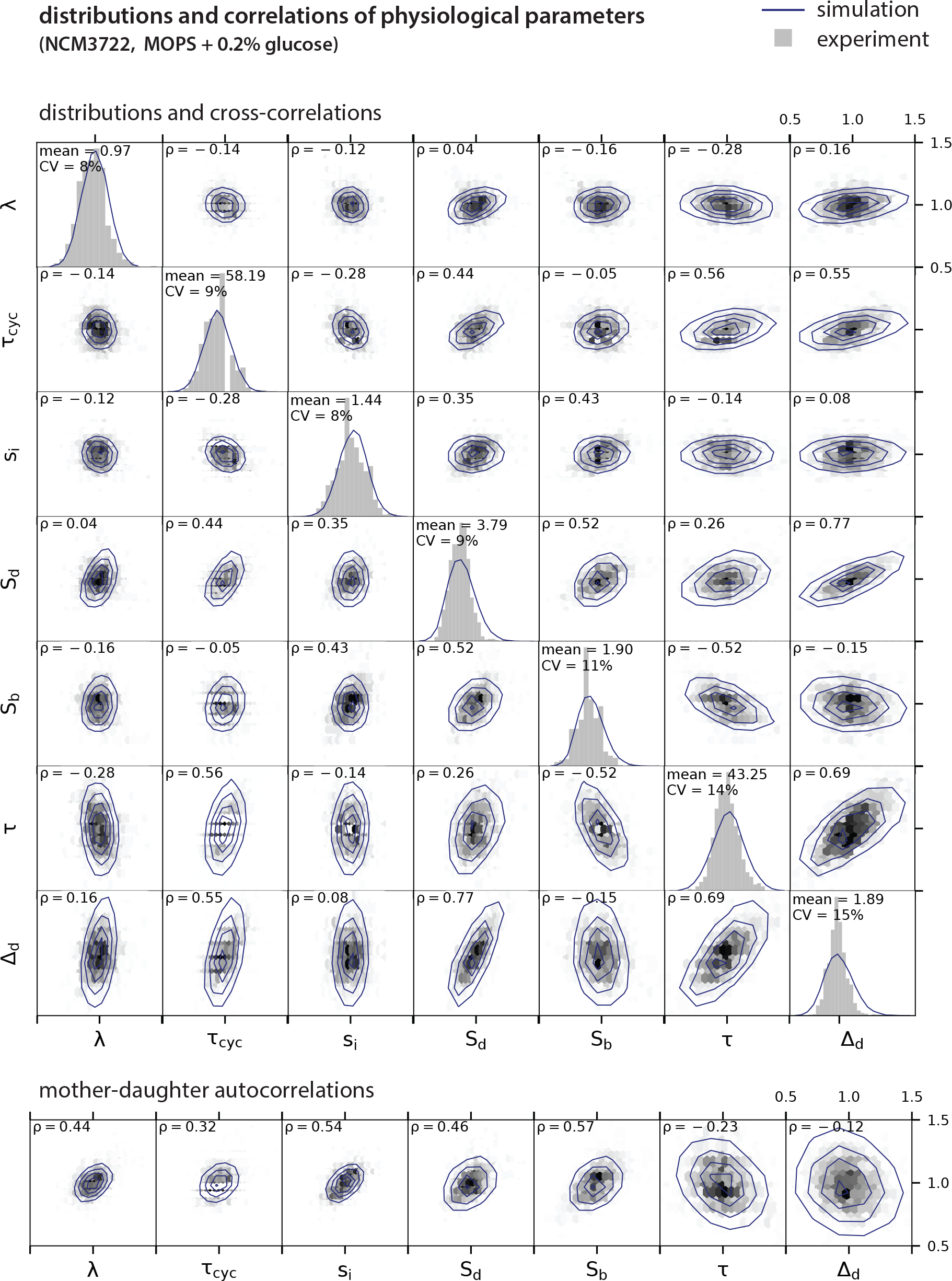
Comparison of experimental measurements with simulations of the Helmstetter-Cooper model. NCM3722 strain with two overlapping cell cycles as in Fig. 1D.

**Supplemental Figure T5:**
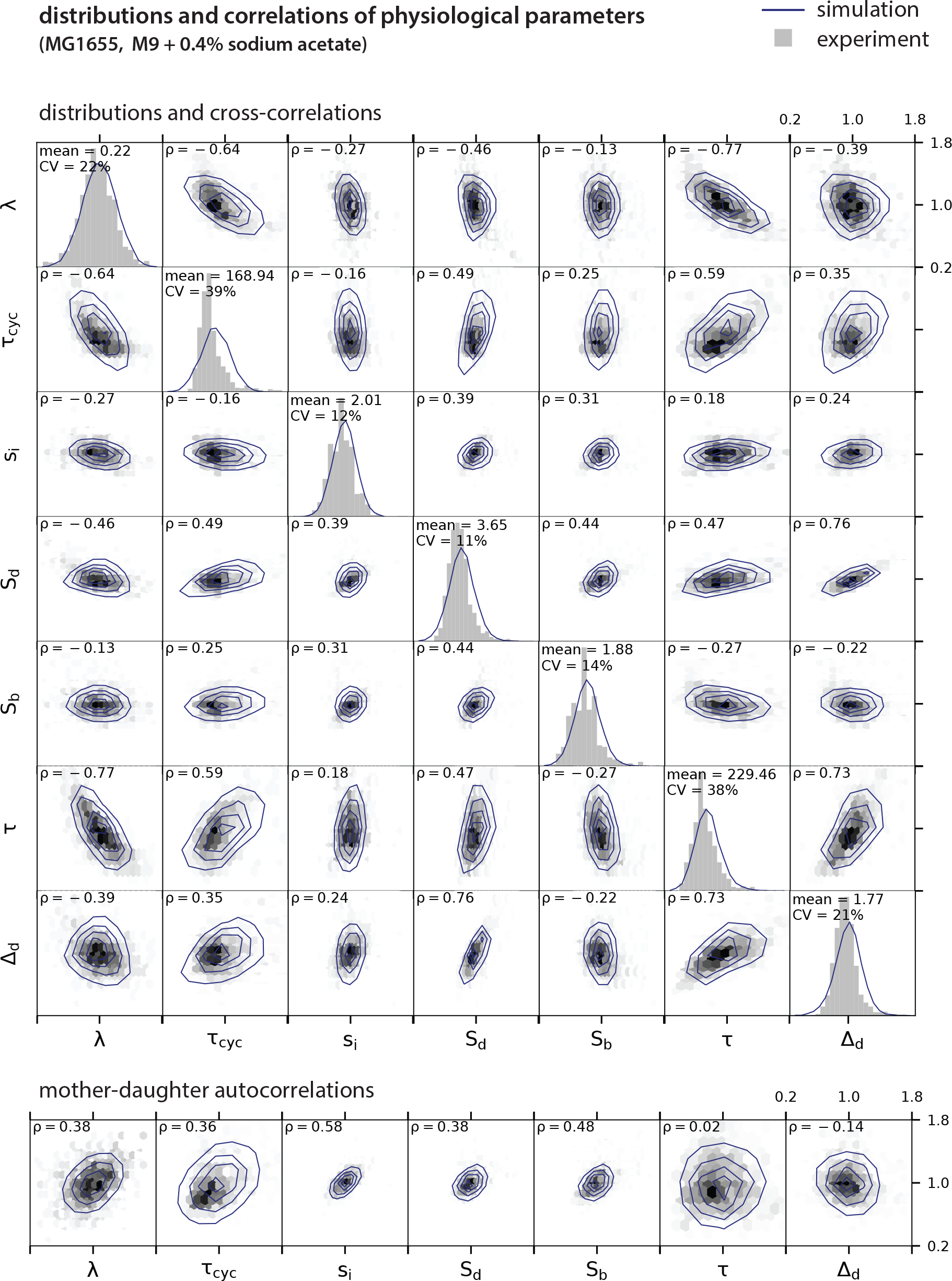
Comparison of experimental measurements with simulations of the Helmstetter-Cooper model. MG1655 strain with no overlapping cell cycles as in Fig. 1D.

**Supplemental Figure T6:**
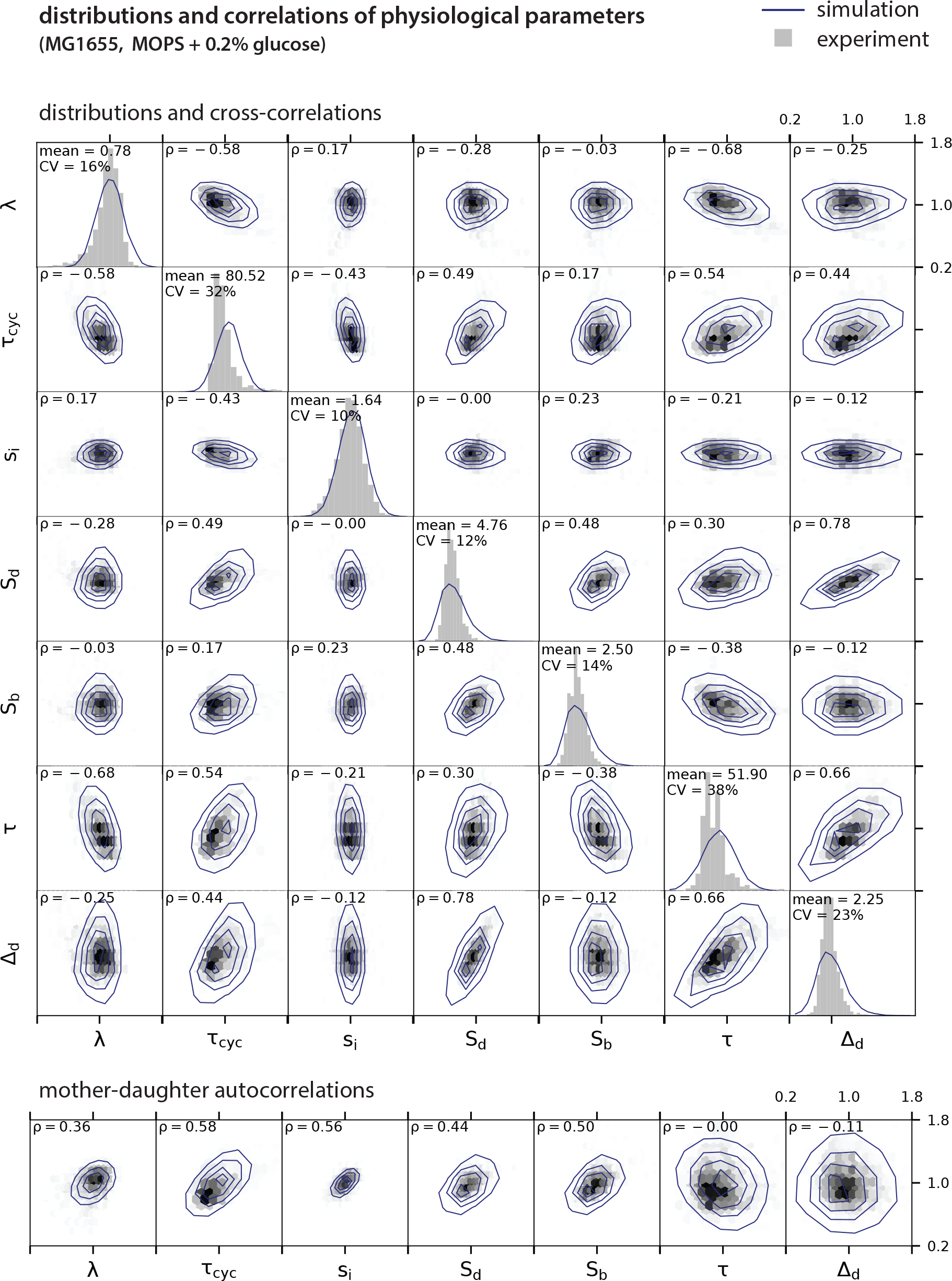
Comparison of experimental measurements with simulations of the Helmstetter-Cooper model. MG1655 strain with two overlapping cell cycles as in Fig. 1D.

**Supplemental Figure T7:**
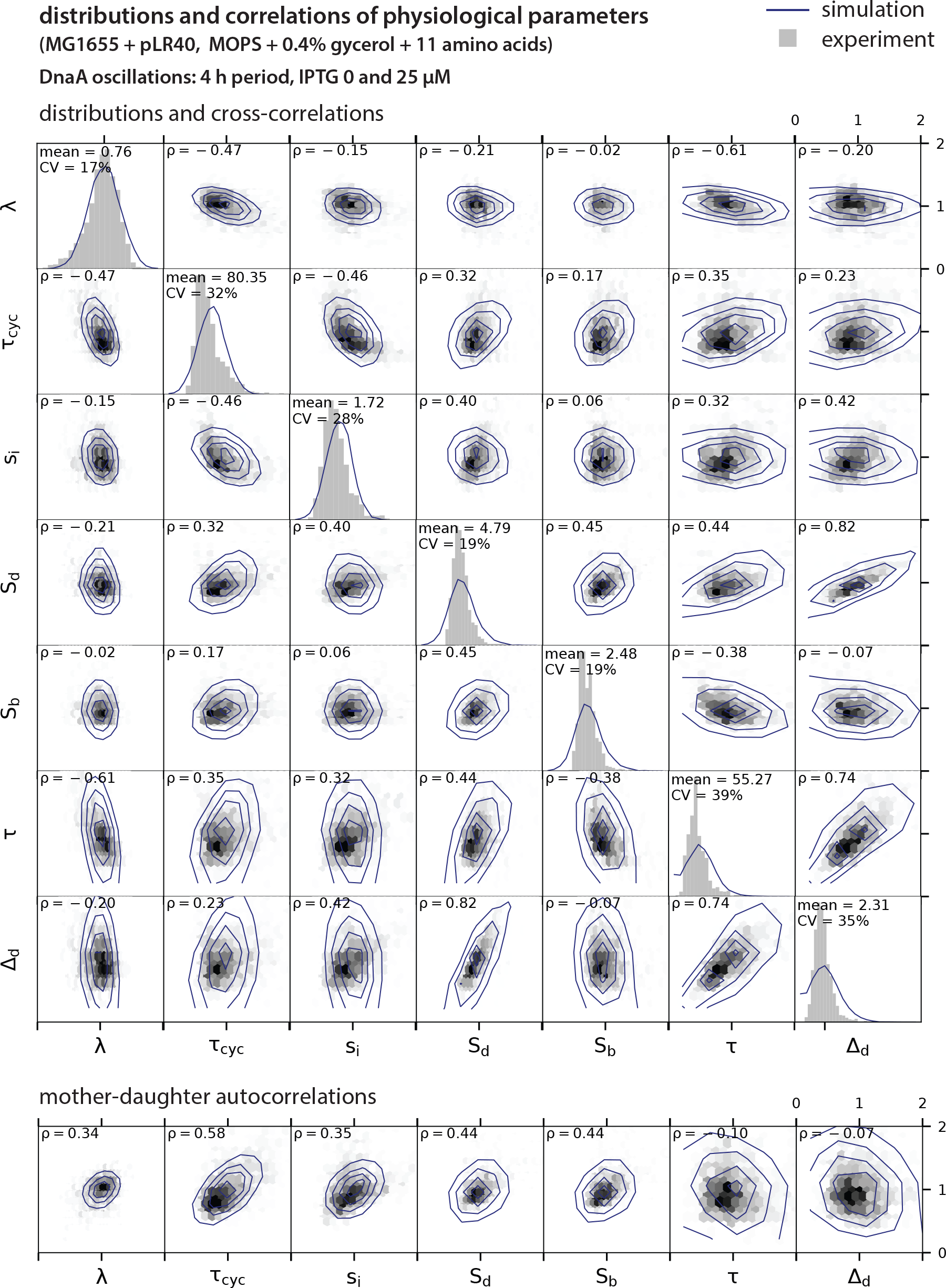
Comparison of experimental measurements with simulations of the Helmstetter-Cooper model. MG1655 strain transformed with pLR40 plasmid for DnaA oscillatory induction as in Fig. 3A.

##### Algorithm 1

Stochastic Helmstetter-Cooper simulation.

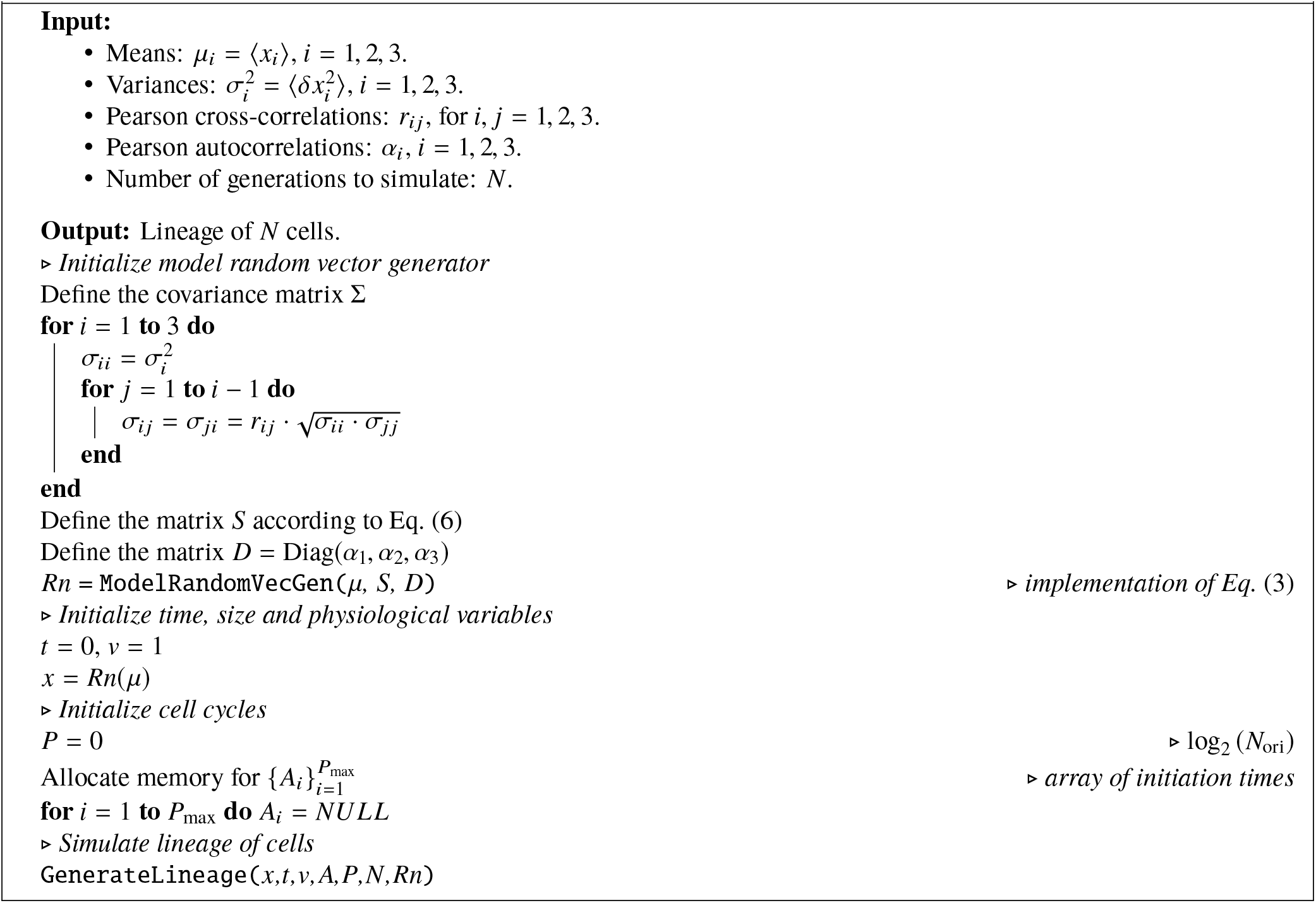

In fact, the Pearson correlation coefficient for the adder at division (resp. at initiation) is uniquely related to the Pearson autocorrelation coefficient of the cell size at division (resp. at initiation). Therefore, the previous equality implies:

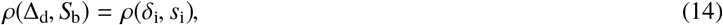

where 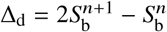 and 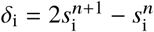.

When relaxing the condition that τ_cyc_ is fixed, Eq. (14) no longer holds. In Suppl. Fig. T9, we investigated whether the two types of adder are still equivalent when τ_cyc_ is allowed to fluctuate. We started from reference values for the parameters and then varied each of the coefficient-of-variations (CVs), cross-correlation and autocorrelation away from their reference value. More accurately, the CVs were varied between 1%-30% and the Pearson correlations between −0.9 and +0.9. However, we did not perturb *ρ*(*λ*, *s*_i_) and *ρ*(*τ*_cyc_, *s*_i_) because experimental data indicate that *s*_i_ should remain very robust to perturbations [1].

We chose 〈λ〉 = ln 2, making the generation time 〈τ〉 = ln 2/〈λ〉 the unit of time. We chose 〈*s*_i_〉 = 1 as unit of cell size. We investigated values of 〈τ_cyc_〉 = 0.5, 1.5, 2.5, corresponding to non-overlapping, 2 overlapping and 3 overlapping cell cycles, respectively. The reference values for the CVs were:

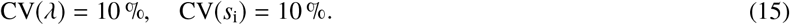

Note that for *τ*_cyc_, we took different CVs for each scenario so as to keep the same amplitude of fluctuations in the cell cycle duration. Specifically, we chose a standard deviation *σ*_*τ*_cyc__ = 0.05, defining a CV of 10 % for the non-overlapping cell cycle scenario but resulting in smaller CVs for the other. The reference matrix of cross-correlations was set to:

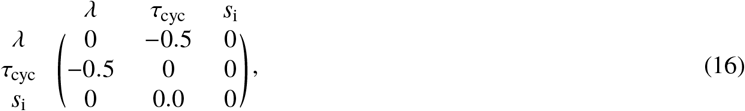

and the reference autocorrelations were taken to be:

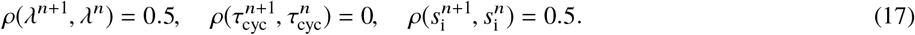

##### Algorithm 2

Function `GenerateLineage`.

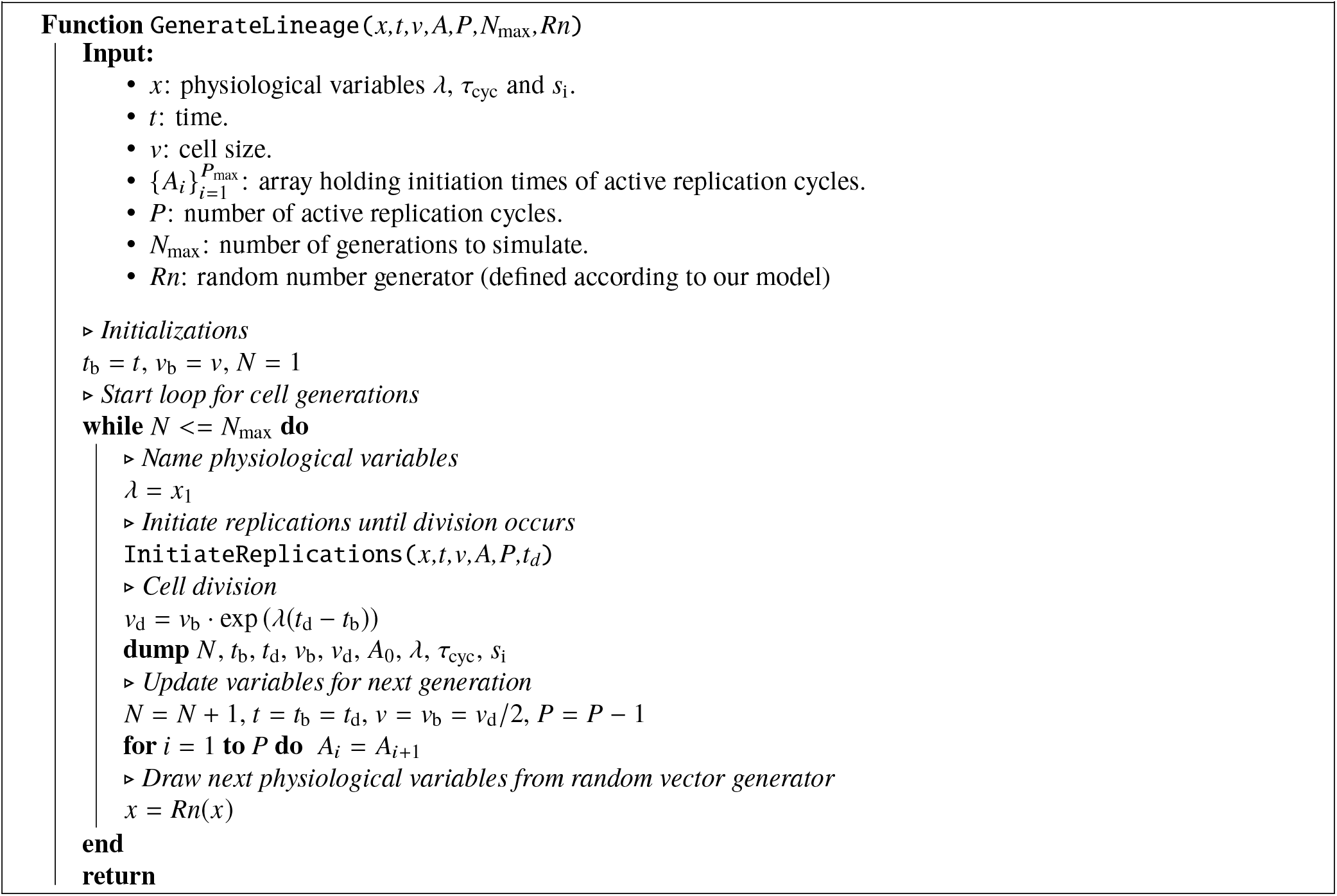

In Suppl. Fig. T9a, the co-regulation hypothesis from Eq. (14) holds for non-overlapping cell cycles. For two and three overlapping cell cycles, deviations from the co-regulation hypothesis are seen. This is due to the sources of noise still present in the system, which tend to uncouple distant generations. For example, reducing the CV of the growth rate to CV(*λ*) = 1 % dramatically reduces these deviations (not shown). Similarly, when increasing the noise in *τ*_cyc_ to *σ*_*τ*_cyc__ = 0.1, deviations from the co-regulation hypothesis are more pronounced (Suppl. Fig. T9B). However, despite the fact that other parameters can affect the division adder correlation (especially for overlapping cell cycles), the effect of the unit cell autocorrelation 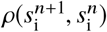 on the value of *ρ*(∆_d_, *S*_b_), was more systematic than cross-correlations or CVs. Therefore we concluded that even when fluctuations are introduced into the Helmstetter-Cooper model, altering the homeostasis of *s*_i_ should affect the cell size homeostasis.

#### F. Adder properties

The size autocorrelation can be used to characterize the cell size behavior. We focus now on the division size properties, yet the following development can also be applied to the initiation size.

##### Algorithm 3

Function `InitiateReplication`.

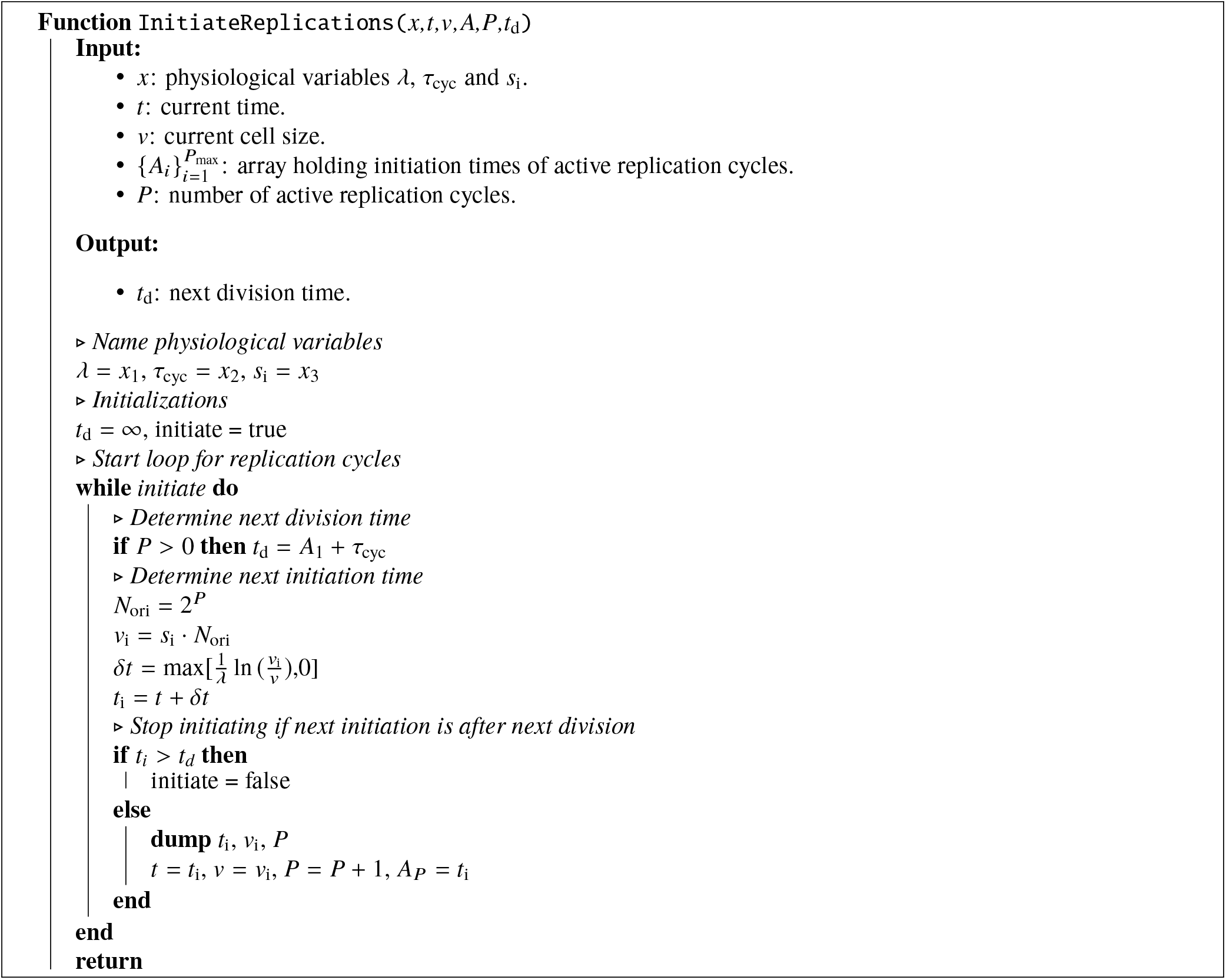

We denote 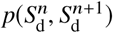 the joint probability distribution of cell size at division for a pair of mother/daughter cells. In the first approximation, we assume it is a Gaussian bivariate distribution with means 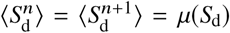 and covariance matrix:

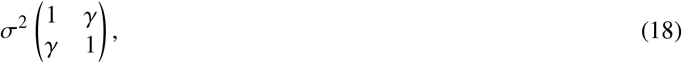

with σ = µ(*S*_d_) · CV(*S*_d_). Consequently, the conditional value of the daughter division size is obtained as (see appendix A):

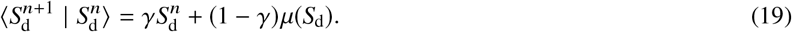

The added size between division for generation *n* is expressed as

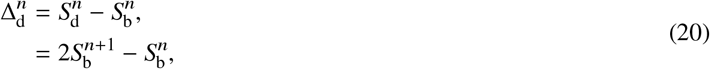

assuming symmetric division. Using the previous equation, one can show that:

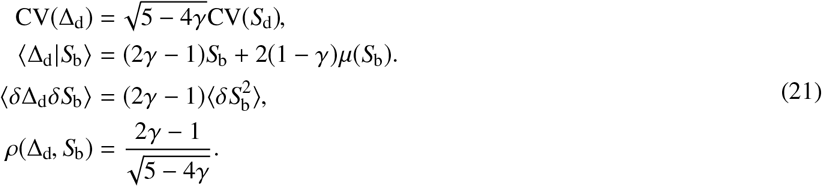

**Supplemental Figure T8:**
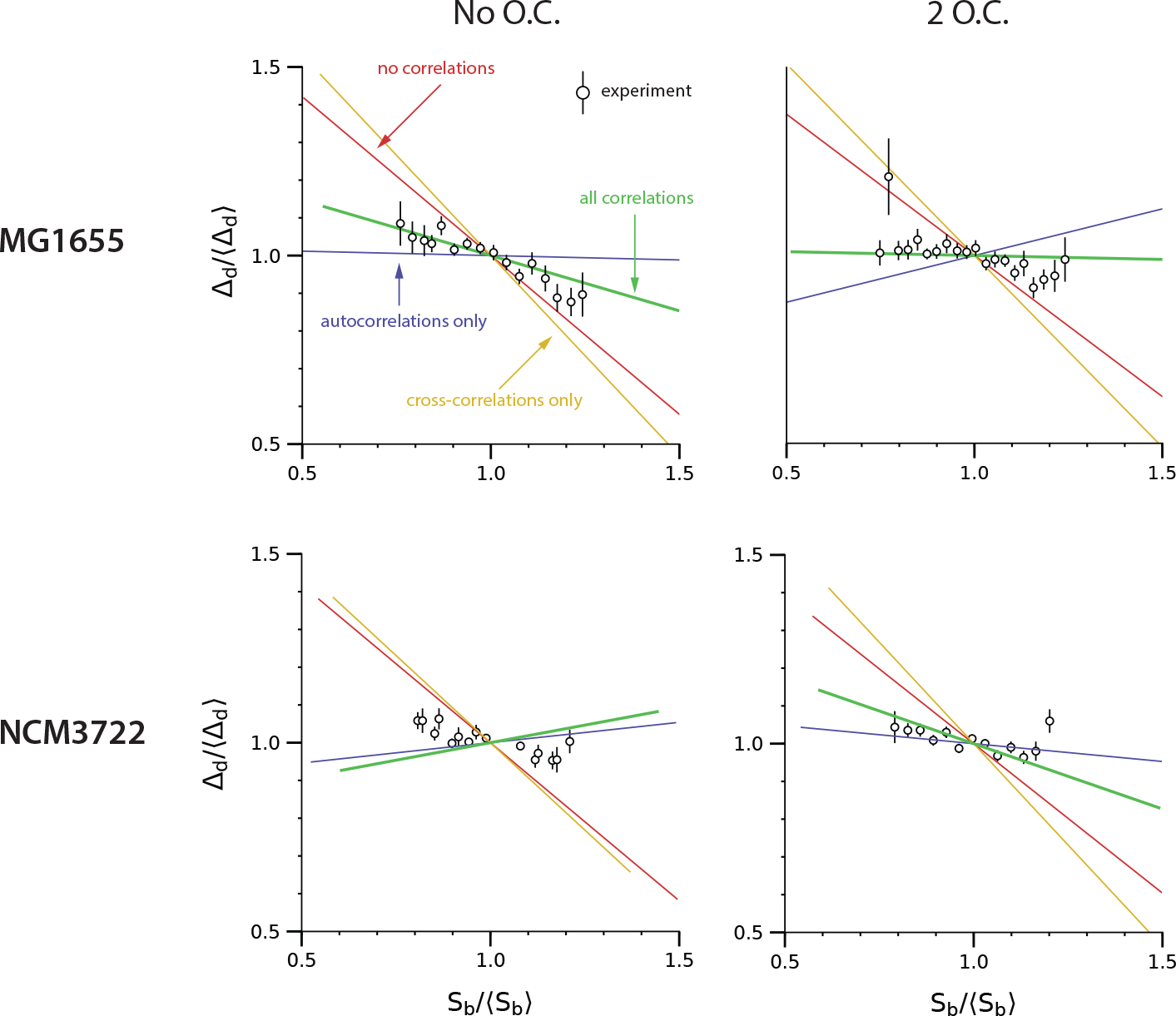
Comparison of the experimental adder behavior with simulation of the stochastic Helmstetter-Cooper model. The simulations are performed without correlations (red), with cross-correlations only (yellow), with autocorrelations only (blue) and with all correlations (green) between physiological variables. The experimental data for each condition were used to set the mean values, coefficient-of-variations and correlations of the joint distribution of the physiological variables.

The last expression is shown in Suppl. Fig. T10. Therefore, when γ = 1/2, the adder principle holds, meaning that the added size is independent of birth size: 〈*δ*∆_d_*δS*_b_〉 = 0 (or *ρ*(∆_d_, *S*_b_) = 0).

### II. THRESHOLD MODELS IN BALANCED GROWTH

#### A. Control of replication initiation

Replication initiation is influenced by several factors, the most important being probably the DnaA protein [17, 18, 19]. The DnaA protein is active when bound to ATP (DnaA-ATP) and inactive when bound to ADP (DnaA-ADP). While both active and inactive forms can bind the *oriC*, evidence indicates that only the active form can trigger initiation [20]. Approximately 10-20 DnaA-ATPs are required at the *oriC* to form a functional complex that can lead to replication initiation [21]. Here we neglect the role of DnaA-ADP, namely as a competitor to *oriC* binding. We therefore consider that replication initiation is under the exclusive control of DnaA-ATP. DnaA binds primarily ATP after being synthesized in the cytoplasm [18, 19], therefore the DnaA-ATP production coincides with the DnaA production. For these reasons we will abusively denote DnaA-ATP as DnaA, which we consider as the replication initiator. We also adopt the simple autorepressor model for the *dnaA* operon [22], *i.e.* the DnaA protein is maintained at a nearly fixed concentration by repressing its own expression.

**Supplemental Figure T9:**
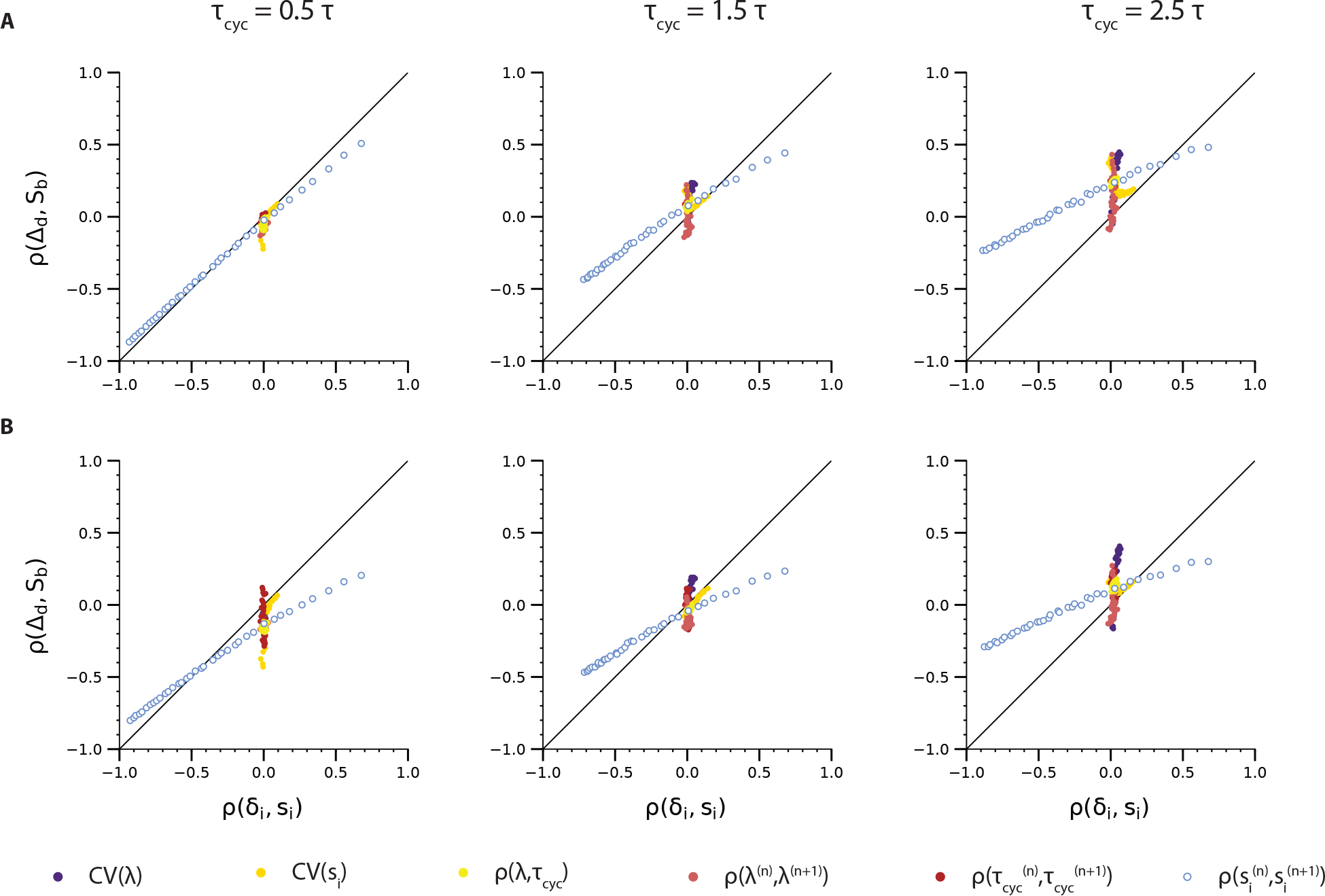
Co-regulation between division adder and initiation adder in the stochastic Helmstetter-Cooper model [Eq. (14)]. **(A)** *σ*_*τ*_cyc__ = 0.05. **(B)** *σ*_*τ*_cyc__ = 0.1, with *σ*_*τ*_cyc__ = 〈*τ*_cyc_〉 · CV(*τ*_cyc_).

#### B. Control of division

Z-ring formation is the predominant division process prior to constriction [23]. Once the Z-ring is functional, division and cell wall machinery proteins bind to this scaffold in order to complete cytokinesis. The Z-ring is made of protofilaments of the essential protein FtsZ. In our experimental assays, we have adopted a nearly functional FtsZ-mVenus fusion protein [24] in order to monitor the assembly of the Z-ring in single-cells. Our observations suggest that FtsZ accumulates to a threshold at the Z-ring. Indeed, the maximum intensity (in a cell life time) at the Z-ring was found to be independent of the cell size at division. This means that on average a fixed, critical amount of FtsZ in the Z-ring is required to trigger the assembly of the division machinery and cell constriction. This threshold mechanism parallels the control of replication initiation by DnaA.

In addition, our experimental assays also suggested that the concentration of FtsZ remains relatively constant during the division cycle, and across many generations. As far as we know, the FtsZ protein does not repress its own expression, like DnaA does. However, we explain this fixed concentration at steady state by postulating that FtsZ production is in balanced growth (see section III).

#### C. Threshold model

Let us now consider a generic protein responsible for the initiation of cell division (note that the same reasoning applies for the control of replication initiation). We assume that this protein accumulates and triggers cell division when its copy number *N* reaches a fixed threshold:

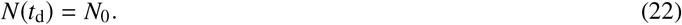

Following cell division, each daughter cell receives *N*_0_/2 copies of the protein. Under balanced growth (see section III), the protein copy number increases in proportion to the cell volume:

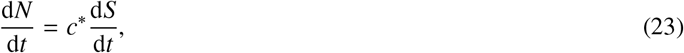

where *c*^∗^ is the steady-state protein concentration. Therefore, one obtains that the added volume from birth to division is given by:

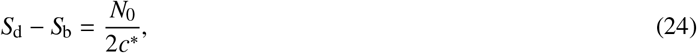

which is the adder principle.

**Supplemental Figure T10:**
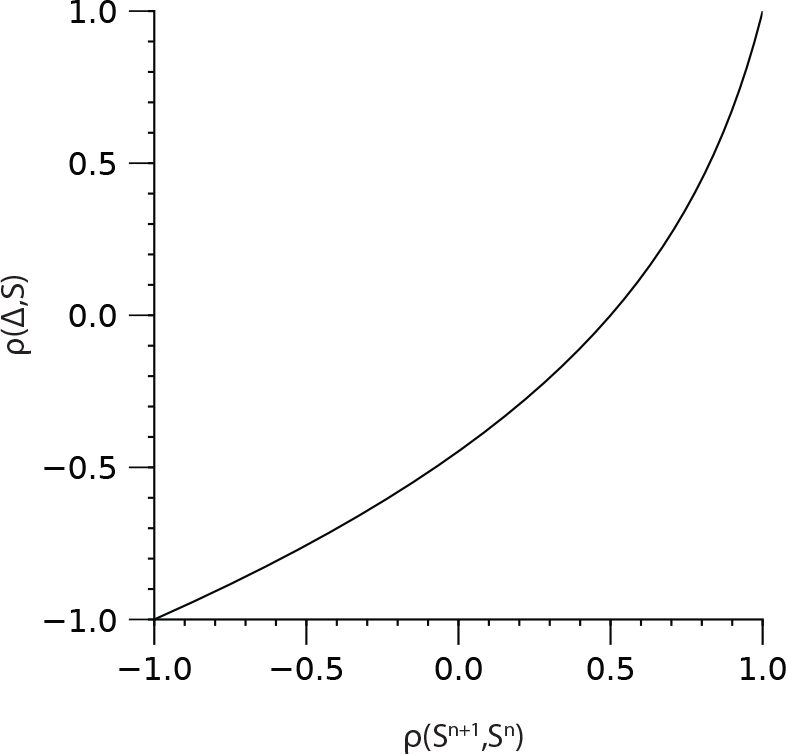
Functional dependence of *ρ*(Δ_d_,*S*_b_) on 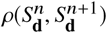 (see Eq. (21)).

The fixed threshold *N*_0_ does not necessarily imply that the cell physically senses the copy number *N* of proteins in the cytoplasm. Instead, *N* can be a proxy for the number of initiators bound to a cell compartment, namely the Z-ring. For example, let us assume that the binding/unbinding dynamics of the proteins in the cytoplasm (*N*) with a cell compartment (with copy number *n*) can be modeled with the linear system:

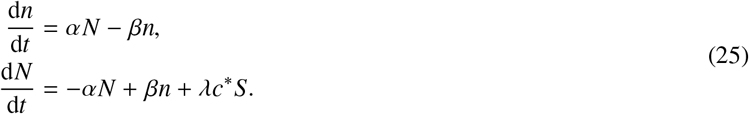

In the previous system, α and β are the rates of binding and unbinding to the cellular compartment, respectively. *λc*^∗^ *S* is the production rate of proteins in the cytoplasm in balanced growth (see section III). Denoting *X* = *n* + *N*, and assuming exponential growth of the cell volume at the rate λ, we immediately have:

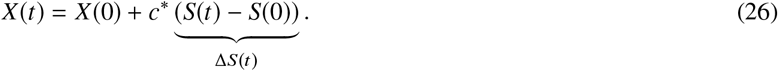

Using the previous equation, we obtain:

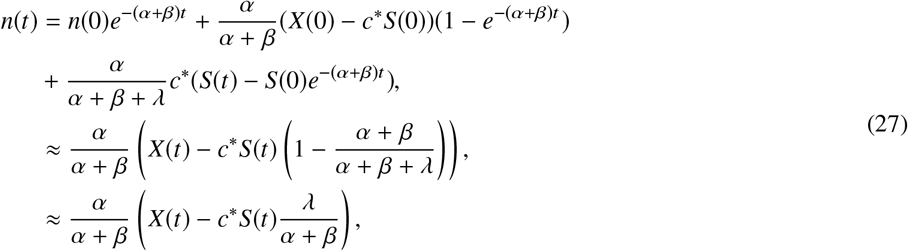

where in the first approximation we assumed that (*α* + *β*)^−1^ ≪ *t*, and in the second approximation that *λ* ≪ *α* + *β*. Therefore, as long as the elongation rate is much smaller than the binding/unbinding rates, the copy number of proteins bound to the cell compartment can be seen as a fixed fraction of the total copy number: *n* = *α*/(*α* + *β*)*X*, and similarly *N* = *β*/(*α* + *β*)*X*. In other words, the protein dynamics is fast compared to growth of the cell, therefore the cytoplasm reservoir and the cell compartment are always at equilibrium. When *λ* becomes comparable to *α* + *β*, deviations from this equilibrium appear. Namely, the number of proteins bound to the cell compartment is below its equilibrium value: *n* < *α*/(*α* + *β*)*X*, meaning that some delay is observed for the cell compartment to reach its threshold. For simplicity, we have considered here that the cytoplasm reservoir and cell compartment are at equilibrium. Eventually, a threshold *n*_0_ to be reached in the cell compartment translates into a threshold *N*_0_ to be reached in the cell cytoplasm, and in a global threshold *X*_0_ to be reached for the total protein copy number.

#### D. Relation to cell size homeostasis

We now consider the general case where *c*^∗^ is subject to fluctuations, and focus on the division size homeostasis. The same reasoning applies to the initiation size homeostasis. Using the definition *c*(*t*_d_) = *N*_0_/*S*(*t*_d_), the Pearson correlation coefficient for division size between consecutive generations is:

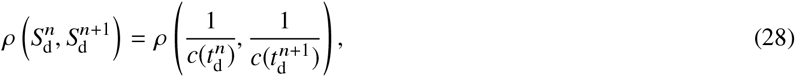

where 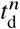 and 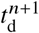 are the times at division for generations *n* and *n* + 1. Provided that the fluctuations in concentration are not too large, the previous expression can be approximated to (see appendix A):

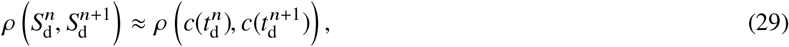

Therefore, cell size homeostasis appears to be linked to the initiator concentration homeostasis. In particular, if fluctuations in the protein concentration occur on time scales much shorter than the generation time, the division size correlation between consecutive generations should vanish, resulting in a “sizer” behavior.

We now relate the mother/daughter division concentrations correlation from Eq. (29) to the time autocorrelation of the protein concentration. Let us consider *L* lineages of cells in a time interval [0, *W*]. Let us denote by *c*_*a,i*_ the concentration at division for the cell corresponding to generation *i* of lineage *a*. The mother/daughter concentration correlation is computed as:

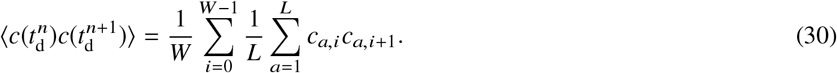

The previous average should converge to a fixed value for large *W* and large *L*. We now assume ergodicity. Specifically, the average in the previous equation with *W* → ∞ and *L* = 1 is equal the average with *W* = 1 and *L* → ∞:

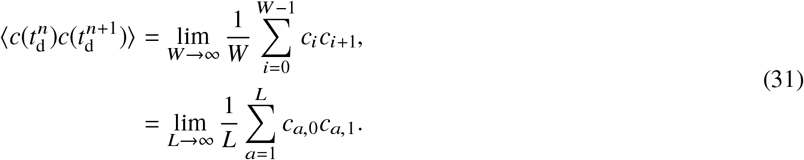

Let us introduce the conditional probability distribution *p(t′|t*) that a daughter cell divides at times *t′* given that its mother cell divided at time *t*. We rewrite the last expression as:

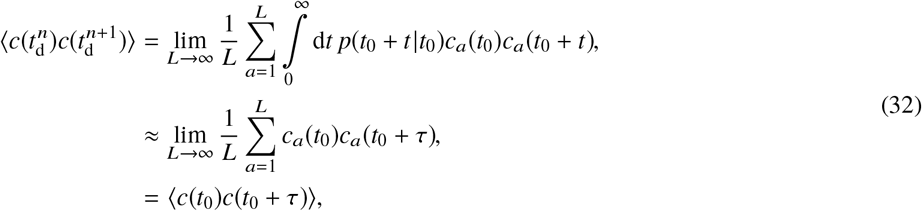

where in the approximation, we assumed *p*(*t*′|*t*) = *δ*(*t′* − *t* − *τ*). This approximation is valid as long as the fluctuations of the concentration are small in an interval [*τ* − *σ*_*τ*_, *τ* + *σ*_*τ*_] centered around the average generation time *τ*, with *σ*_*τ*_ being the width of its distribution (the typical fluctuations of the generation time). In the last expression, the concentration *c*(*t*) should be understood as a continuous stochastic process. Therefore, to a particular lineage of cells *a* corresponds one stochastic continuous process *c*_*a*_(*t*). The brackets means that an average is taken over all the realizations of {*c*_*a*_(*t*)}_*a*=1...*L*_. The previous equation is also valid for the centered concentration *δc*(*t*) = *c*(*t*) − 〈*c*〉. As a result, we obtain:

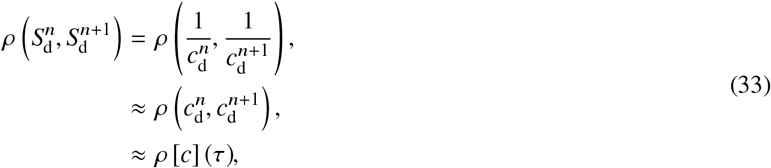

where:

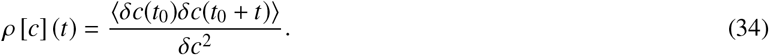

#### E. Adder property

In the next section, we will show that under balanced growth, *ρ*[*c*](*τ* = 1/2. From Eq. (33), and using the relations from Eq. (21), we conclude that:

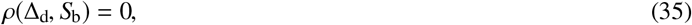

which is an other formulation of the adder principle.

### III. REPROGRAMMING CELL SIZE HOMEOSTASIS BY BREAKING BALANCED GROWTH

#### A. Balanced growth

Consider a type of protein whose mass fraction in the cell is *φ*^∗^ at steady state. If we denote by *m* the mass of these proteins, and *M* the total dry mass of the cell, we have in balanced growth [25, 26]:

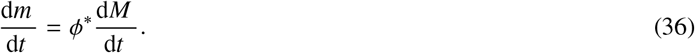

To reformulate the previous equation in terms of the protein copy number *N* in the cell, we introduce the mass of one protein *m*_*P*_, the cell volumic mass *ρ*_*c*_ and the cell size or volume *S*. A simple rewriting leads to:

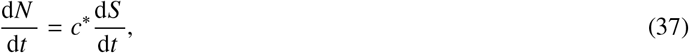

where *c*^∗^ = *φ*^∗^ *ρ*_*c*_/*m*_*P*_ is the protein concentration at steady state. The protein concentration is *c* = *N/S*, thus satisfies the first order differential equation:

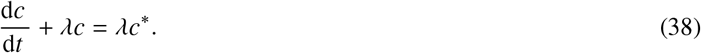

In Eq. (38), steady state is achieved when the protein synthesis rate per unit of volume, *λc*^∗^, balances the decrease in protein concentration due to dilution, *λc*.

#### B. Time-dependent production rate

Let us suppose now that the production rate in Eq. (38) is not constant. Instead, the protein synthesis allocation is a time-dependent function *p*(*t*). The protein concentration obeys the differential equation:

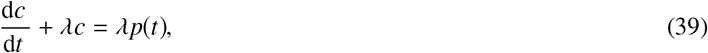

The solution of this ODE is:

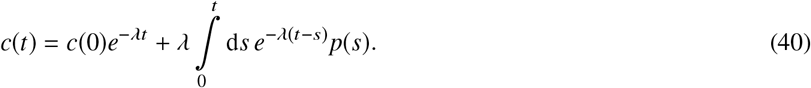

When the production rate is a periodic function of time, the steady state solution for the protein concentration will also be a periodic function with same period. For example, we give *c*(*t*) for the cases of cosine and periodic square production rates in Suppl. Tables T1 and T2. In practice, a time-dependent production rate can be achieved by imposing a time-dependent induction of a promoter. In particular, a periodic square production is obtained by switching between a medium without the inducer and a medium with the inducer every half-period.

**Supplemental Table T1:**
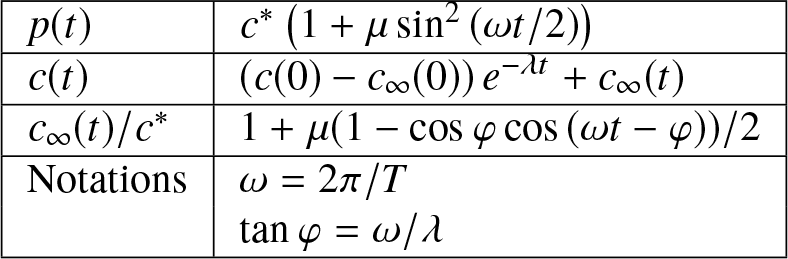
Protein concentration for a cosine production rate.

**Supplemental Table T2:**
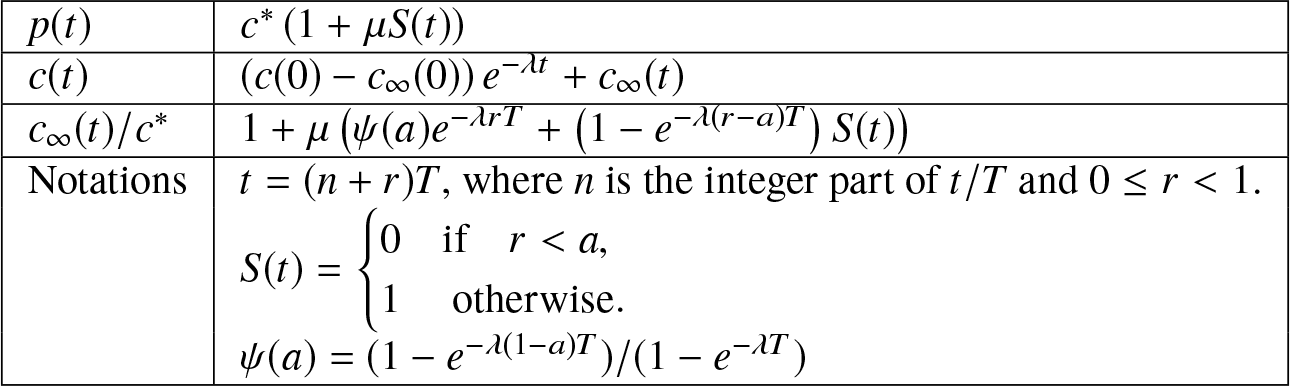
Protein concentration for a periodic production rate with pulses. The periodic square function is obtained when *a* = 1/2.

#### C. Stochastic production rate

Here we consider that the protein synthesis allocation undergoes stochastic fluctuations. The protein concentration obeys the differential equation:

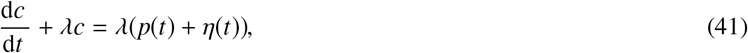

where *η*(*t*) is a Gaussian white noise such that 〈*η* (*t*)〉 = 0 and 〈*η*(*t*)*η*(*t* ′)〉 = 2Γ*δ* (*t-t* ′). The brackets denote an average over different realizations of the noise, for example over many different cells subject to the same production rate (*e.g.* through the same induction). We may decompose the deterministic and stochastic contributions by writing *c*(*t*) = 〈*c*(*t*)〉 + *y*(*t*). The average concentration 〈*c*(*t*)〉 follows the deterministic ODE in Eq. (39) while the fluctuations around the average are expressed as:

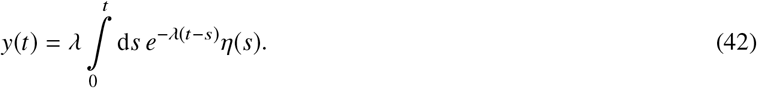

Being a sum of Gaussian random variables *y*(*t*) is also a Gaussian random variable, with mean 〈*y*(*t*)〉 = 0, and variance:

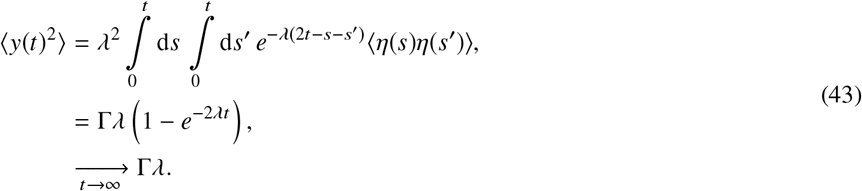

Similarly, the two-point correlation is:

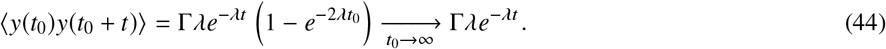

#### D. Concentration autocorrelation

In section II we have presented a model in which cell size homeostasis is driven by the autocorrelation function of division proteins concentration. Here we first give this time autocorrelation function in balanced growth, when the production rate of these protein is fixed. We then show how the autocorrelation function is modified when the production rate oscillates.

##### D1) Fixed production rate

In balanced growth, the production rate of proteins is fixed, namely *p*(*t*) = *φ*^∗^. Thus 〈*c*〉 = *c*^∗^. The Pearson time autocorrelation coefficient for protein concentration is defined as:

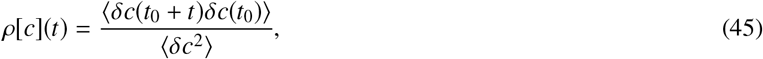

where *δc*(*t*) = *c*(*t*)−〈*c*〉 and the brackets denote an average over different realizations of the stochastic process in Eq. (41) (*i.e.* different lineages). Using Eq. (44), we obtain

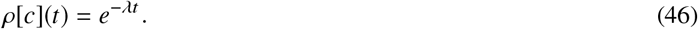

In particular, for *t* = *τ* = ln 2/〈*λ*〉, we have *ρ*[*c*](*τ*) = 1/2, which together with Eq. (33) ensures the adder behavior for cell size homeostasis in balanced growth.

##### D2) Time-dependent production rate

For a time-dependent production rate, the expression in Eq. (45) must be revised because time translational invariance is broken, and it is necessary to take into account variations in time for the production rate. In particular, the average concentration 〈*c*(*t*)〉 is a function of time. For a periodic production rate with period *T*, ergodicity can still be assumed, but Eq. (31) is modified to:

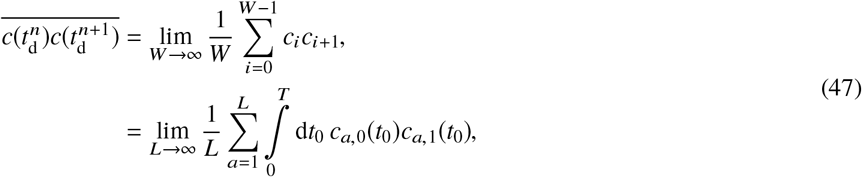

where *c*_*a*,0_(*t*_0_) and *c*_*a*,1_(*t*_0_) denote the concentrations of division proteins at division for a pair of mother/daughter cells such that the mother divides at *t*_0_. We used a bar symbol for this average to make a distinction from the previous average with brackets. Thus, Eq. (32) becomes:

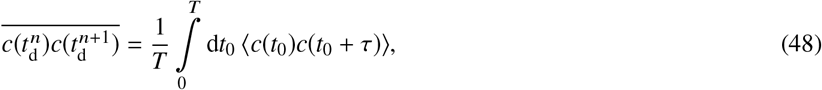

where the brackets as before denote an average over different lineages. The average concentration now reads:

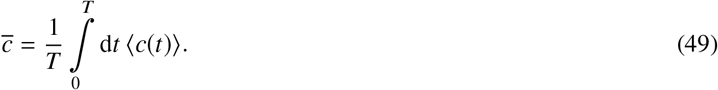

We rewrite the connected correlation for mother/daughter concentrations at division:

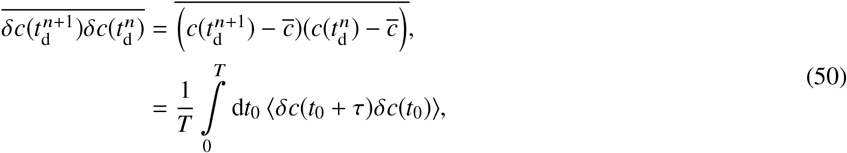

where 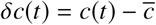 and *τ* is the mean generation time.

The last expression in Eq. (50) is the two-point correlation evaluated at *t* = *τ*. It can be decomposed as a sum of a deterministic contribution due to the time variations of the production rate, and a stochastic contribution due to the stochasticity in Eq. (41):

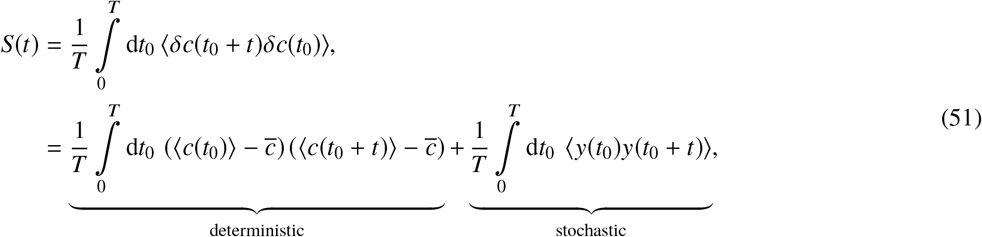

where as before *y*(*t*) = *c*(*t*) − 〈*c*(*t*)〉. Finally, the Pearson time autocorrelation coefficient is expressed as:

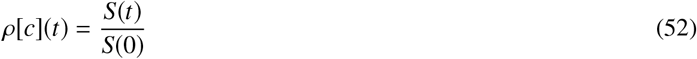

For example, for a cosine production rate (Suppl. Table T1), we find:

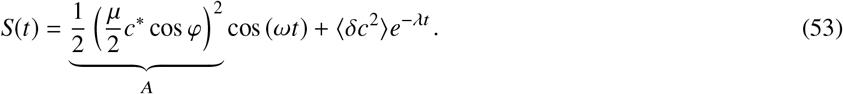

There are two contributions in the two-point correlation from the previous expression. The second is due to the inherent stochasticity in the protein production, while the first is imposed by the specific shape of the production rate function. We see that a careful choice of the amplitude of the oscillations *µ*, and of the period of the oscillations *T* (which determines the value of cos *φ*), can lead to *S*(*τ*) < 0. For example, taking *T* = 2*τ* leads to:

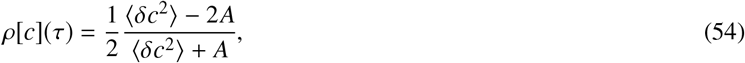

which can be made negative by increasing *µ*. Similarly, taking *T* = 4*τ* leads to:

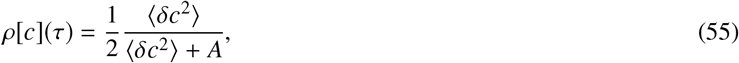

which is smaller than the value of 1/2 from balanced growth and converges to zero when increasing *µ*. As can be seen, the adjustable parameter to tune the autocorrelation coefficient is the ratio 〈*δc*^2^〉/*A* which essentially quantifies the stochastic fluctuations of protein concentration versus the amplitude of the induced oscillations of mean protein concentration. In particular when 〈*δc*^2^〉 ≫ *A*, one retrieves the static result: *ρ*(τ) → 1/2. On the contrary in the limit of vanishing noise 〈*δc*^2^〉 ≪ *A*, one obtains as expected *ρ*[*c*](*τ*) = −1 when *T* = 2*τ* and *ρ*[*c*](*τ*) = 0 when *T* = 4*τ*.

An undesired property of the time-autocorrelation function obtained from Eq. (53) is that for a period *T* ≫ *τ*, it does not converge to the exponential function from balanced growth. This comes from the fact that deviations in the concentration are taken with respect to the total average defined in Eq. (49). However, when *T* ≫ *τ*, the concentration is approximately constant in time intervals of length *τ*. Namely, when analyzing fluctuations, deviations should be considered around the average concentration in this interval, say [*t*, *t* + *τ*], rather than the average in Eq. (53). To circumvent this problem, we define the concentration average in a window of size *T*_*w*_:

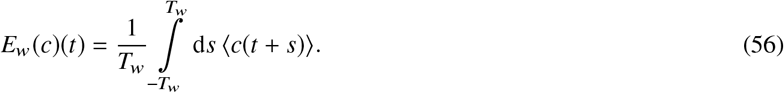

Substituting 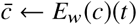 in Eq. (51), we obtain for the cosine induction:

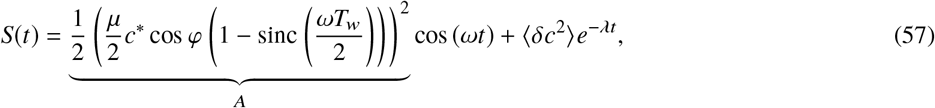

where sinc(*x*)= sin (*x*)/*x*. Effectively, this corresponds to rescaling the amplitude by a factor (1 - sinc(*ωT*_*w*_/2)). Considering that the generation time is the relevant time-scale for studying protein concentration fluctuations, we may take for simplicity *T*_*w*_ = 2*τ*. Therefore *ωT*_*w*_/2 = 2*πτ*/*T*, and for *T* ≫ *τ*, we retrieve the exponential function in Eq. (46) corresponding to a fixed induction, namely balanced growth. With such a definition, we show in Suppl. Fig. T11 how the concentration time-autocorrelation function varies when the induction amplitude µ and the period of oscillations *T* are changed for both cosine and square inductions.

#### E. Simulations of the combined thresholds model

We used simulations to generate lineages of cells according to the combined thresholds model. Here we start considering the growth rate is time-dependent, as discussed in appendix B. Following Eq. (94), we assume that each cell grows exponentially its size according to:

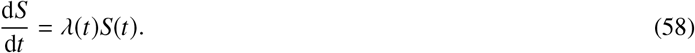

**Supplemental Figure T11:**
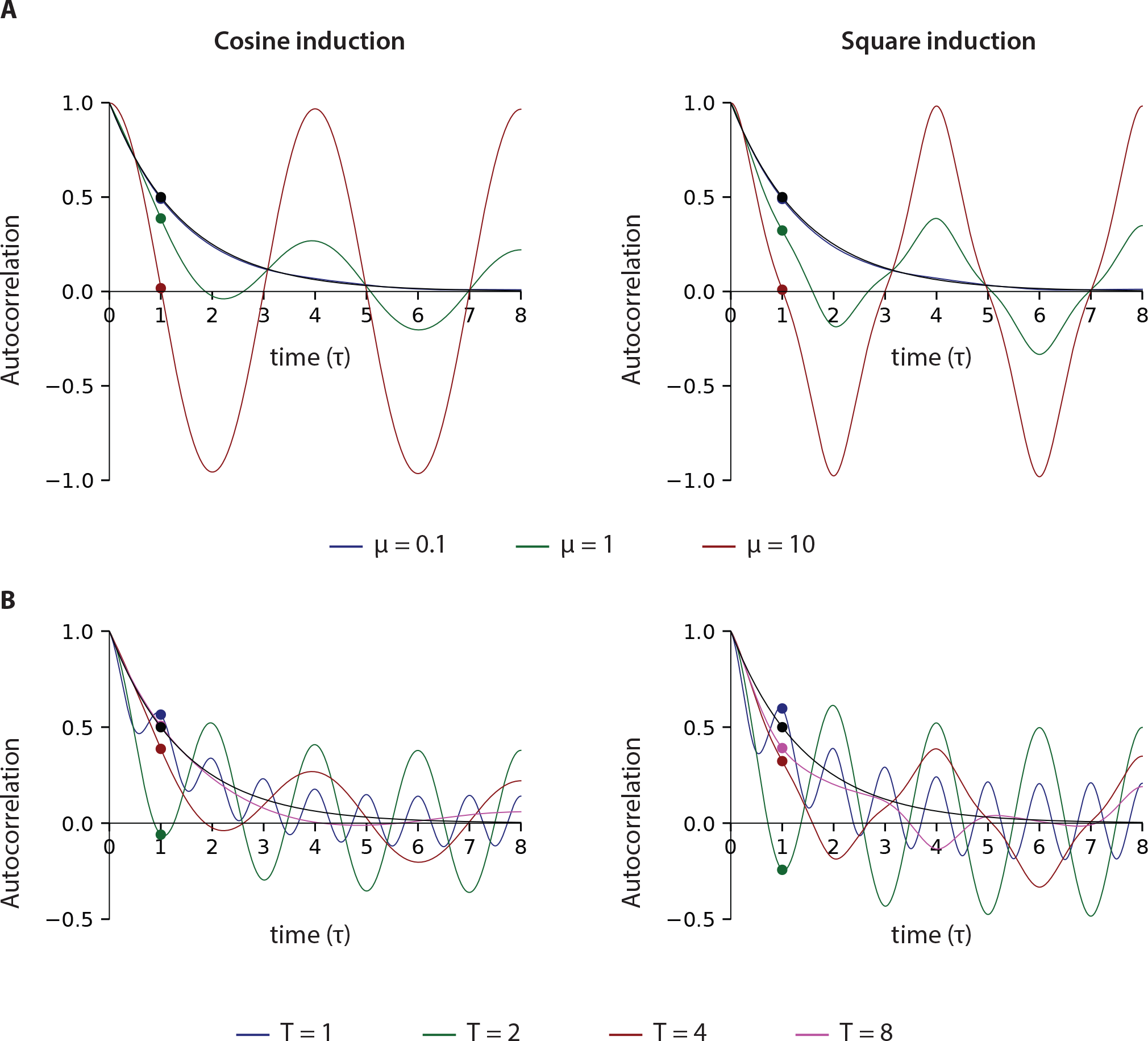
Concentration autocorrelation function for a cosine induction or for a square induction. **(A)** Dependence of the autocorrelation function on the amplitude of induction *µ* (we used a period *T* = 4). **(B)** Dependence of the autocorrelation function on the period of induction *T* (we used an amplitude *µ* = 1). The autocorrelation functions were computed by simulating the stochastic dynamics as defined in Eq. (41) for 10 000 trajectories and using Eq. (51). We defined the generation time *τ* = ln 2/*λ* as unit of time and we defined the unit of concentration so that the lower steady state value is *c*^∗^ = 1. The level of noise was set by taking 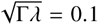, so that the CV of the concentration for a constant induction is 10 %. To compute the autocorrelation we used the running window average from Eq. (56) with *T_w_* = 2τ instead of the total average from Eq. (49). The autocorrelation function for a constant production rate, *i.e.* 2^−*t*/*τ*^, is denoted by a black line.

To describe the time-evolution of the instantaneous growth rate *λ*(*t*), we linearized Eq. (97) around the steady state growth rate λ^∗^, and introduced stochastic fluctuations:

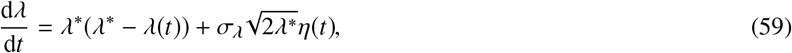

where *η*(*t*) is a Gaussian white noise with correlator 〈*η*(*t*) · (*t* ′)〉 = *δ* (*t′−t*). The normalization ensures that the linearized process (Ornstein-Uhlenbeck type) is such that the growth rate fluctuations at steady-state are 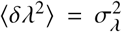. Similarly to describe the time-evolution of the concentration of initiation proteins, *c*_I_, and of the concentration of division proteins, *c*_D_, we linearized Eq. (100) around the steady states 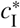 and 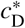, and introduced stochastic fluctuations:

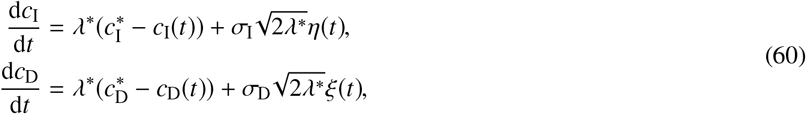

where 〈*η*(*t*) · *η*(*t*′)〉 = *δ*(*t*′ − *t*) and 〈*ξ*(*t*) · *ξ*(*t*′)〉 = *δ*(*t* ′ − *t*). Again, the normalizations ensure that the fluctuations of protein concentrations at steady state are such that 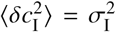 and 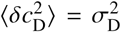. Note that in our implementation, the steady state concentrations 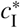 and 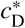 are time-dependent too since they vary with the induction level of the protein of interest. However, due to the periodic square induction, they remain constant between switches.

Initiation occurred when the total copy number of of initiation proteins per origin reached a fixed threshold: 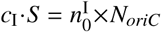. Similarly, division occurred when the total copy number reached a fixed threshold: 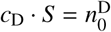. The practical implementation is described in Algorithms 4 to 6.

We have applied this method to simulate oscillation experiments performed in the laboratory. In Suppl. Fig. T12, we used 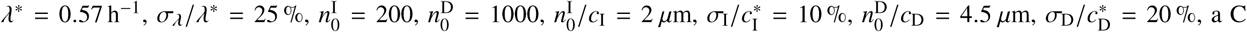 period of 40 min. Note that for simplicity, we expressed concentrations per unit of length since the cell width is constant. We generated 100 lineages of 100 cells. As can be seen in Suppl. Fig. T12A, the experimental distributions are well reproduced in our simulations. Similarly the adder plot is consistent with experimental data (Suppl. Fig. T12B). The autocorrelation function for the concentration of division proteins is well reproduced for short times, however some discrepancies arise from intermediate to long times (Suppl. Fig. T12C). We suspect this is due to some uncontrolled noise in our experimental readout for division protein concentration, because we use a fluorescent signal as a proxy. Similarly the autocorrelation function for division proteins is in good agreement with experimental data for short lags and more discrepancy arise for long lags (Suppl. Fig. T12D). Finally note the agreement between our model and the experimental measurements for protein concentration dynamics is very good (Suppl. Fig. T12E). The second plot emphasizes that our model for protein concentration dynamics based on balanced growth (Eqs. (97) and (100)) and instantaneous change in the fixed protein allocation *c*^∗^ is accurate. The third plot in Suppl. Fig. T12E uses the threshold model assumption 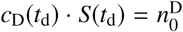. Despite some discrepancy, probably due to the simplicity of this model, we note a good agreement with the observed oscillations of cell size.

**Supplemental Figure T12:**
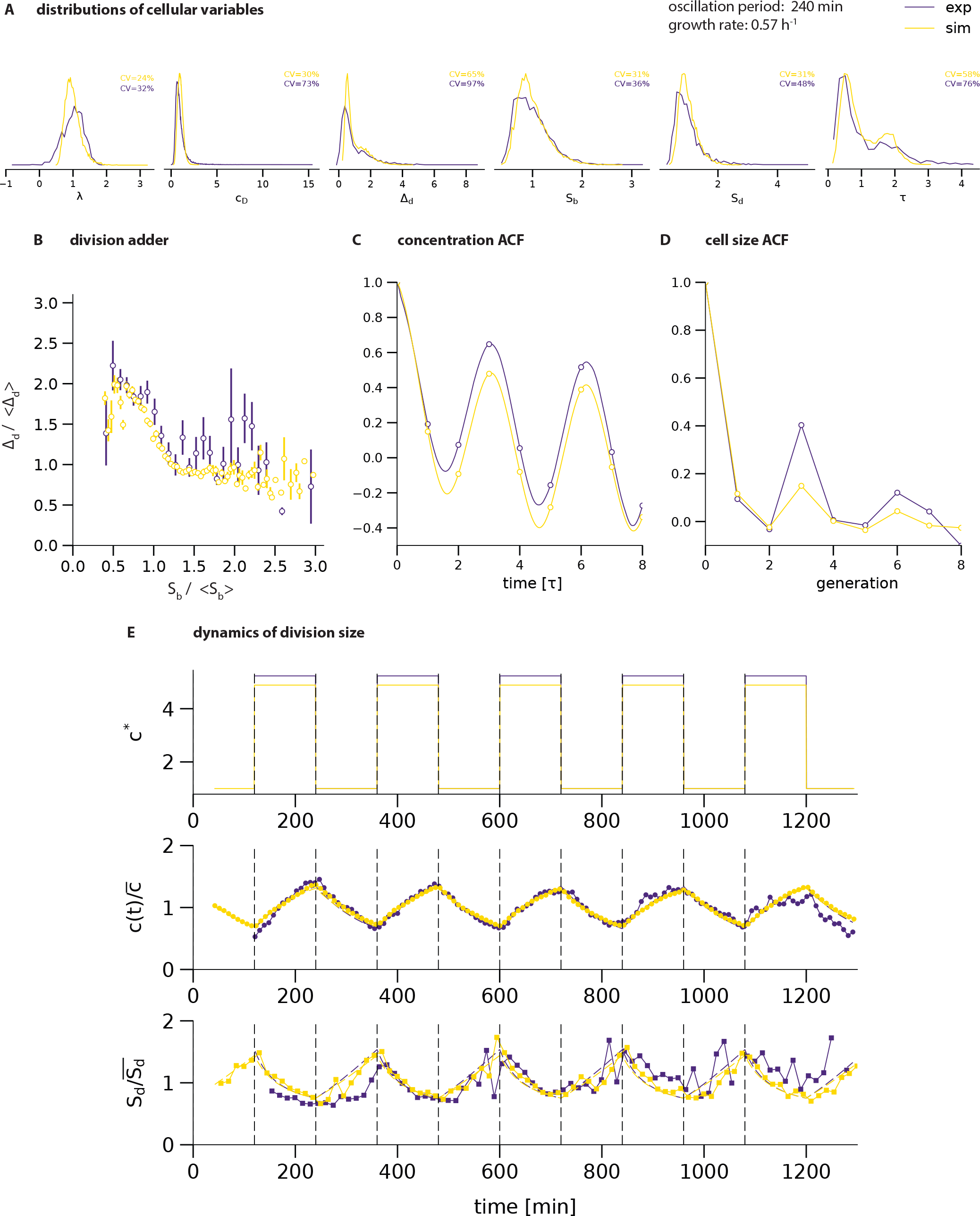
Overlay of experimental results and simulation of the combined threshold model.

##### Algorithm 4

Combined threshold models simulation.

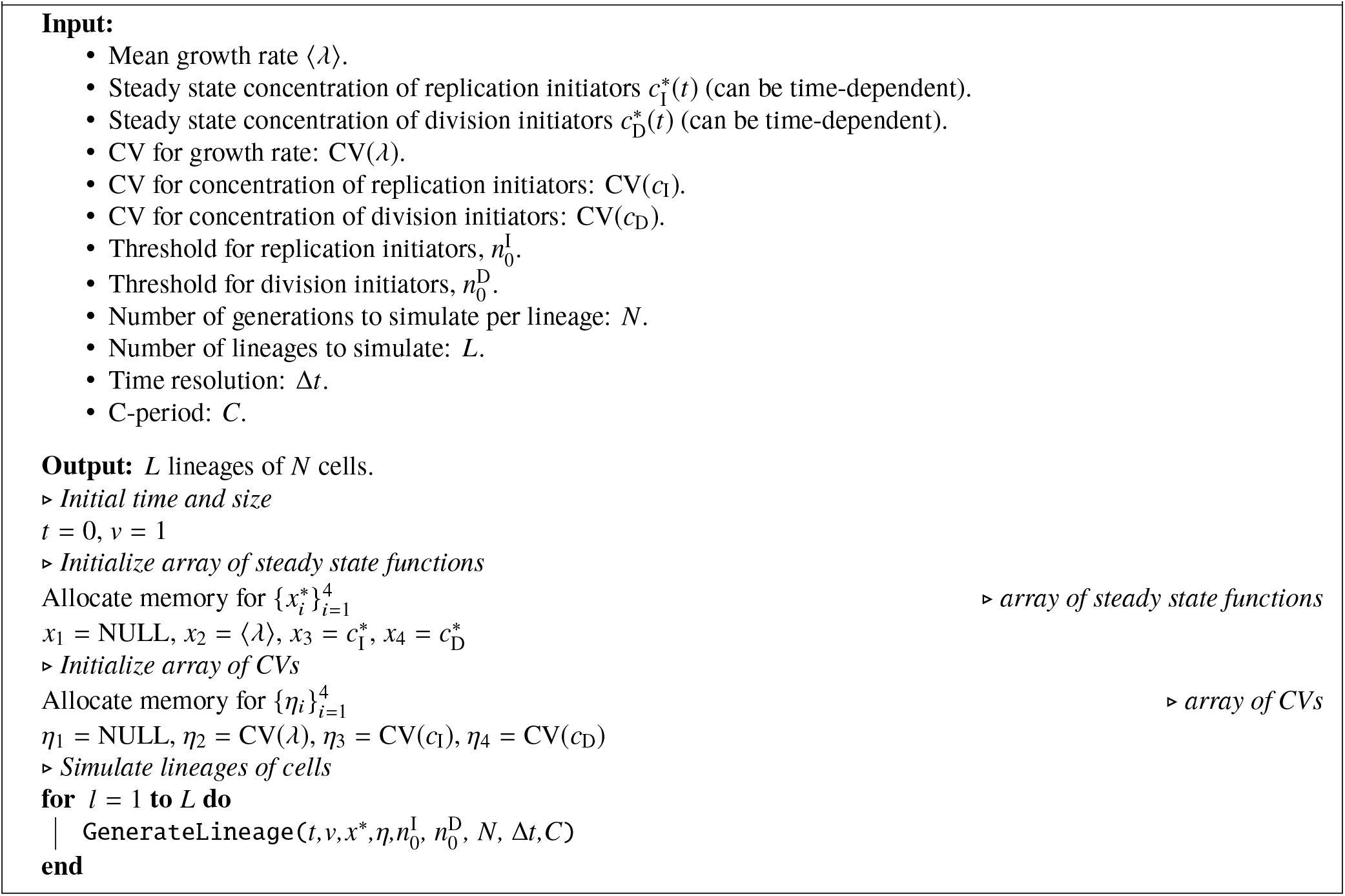

##### Algorithm 5

GenerateLineage function for threshold models simulation.

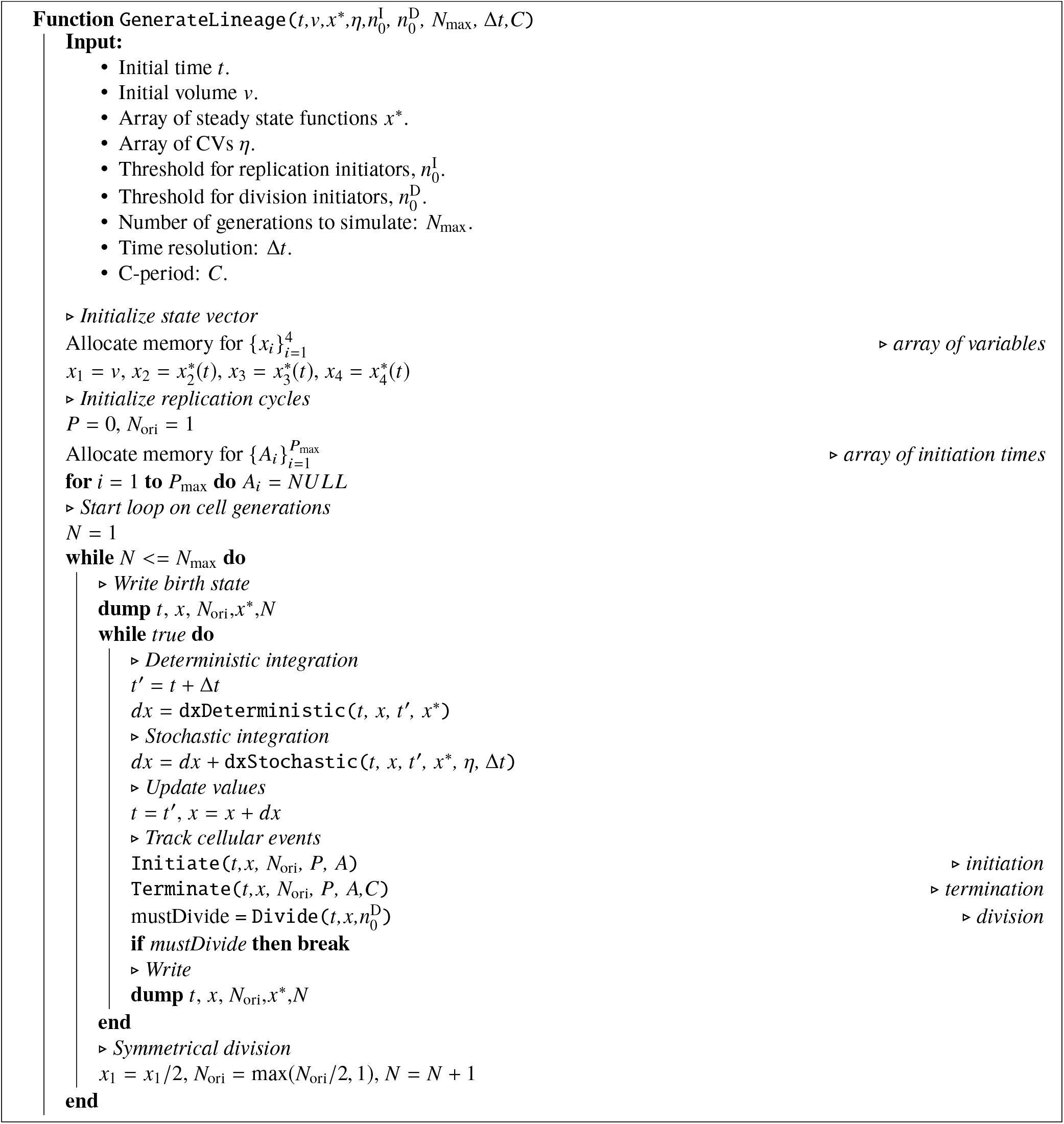

##### Algorithm 6

Functions for threshold models simulation.

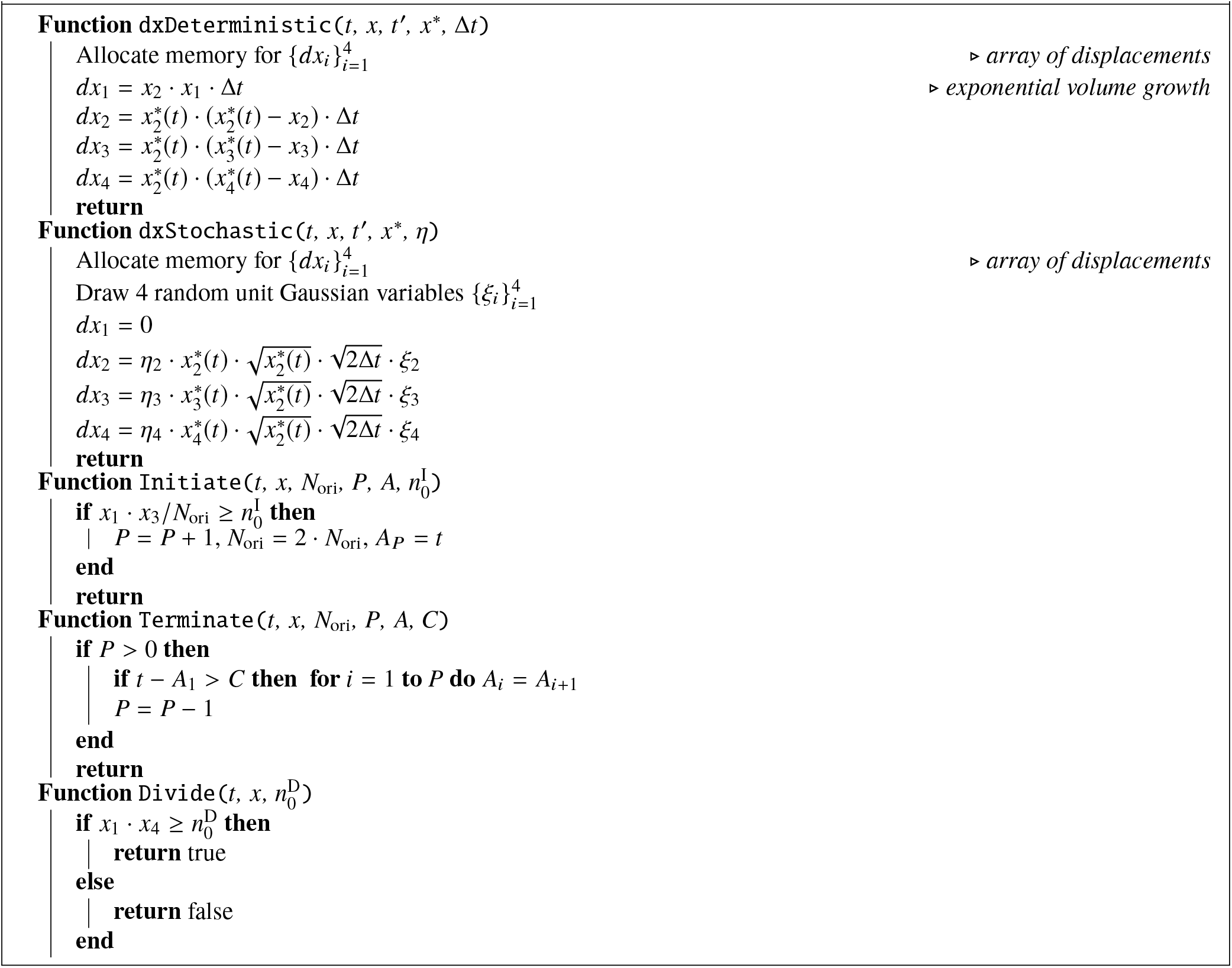

**Supplemental Table T3:** Correspondence with notations in Campos *et al.*[12].

| Parameter | Campos *et al.* | Notation |
| --- | --- | --- |
| Growth rate | $\alpha$ | $\lambda$ |
| Size at birth | $L_b$ | $S_b$ |
| Size at division | $L_d$ | $S_d$ |
| Added size from birth to division | $\Delta L$ | $\Delta_d$ |
| Added size between X-events | $\Delta L^*$ | $\Delta_d$ |
| Elapsed time between birth and division | $T$ | $\tau$ |
| Elapsed time between X-event and division | $\delta t$ | $\tau_X$ |

### IV. DISCUSSION ON THE PHASE-SHIFTED MODEL PREVIOUSLY REPORTED

#### A. Overview

In their paper [12], Campos and colleagues presented experimental evidences of a “constant elongation model”, stating that each individual cell grows in average of a constant mass between birth and division. This result is also known today as the adder principle [13]. Comparison of the distributions of the added size and of the birth size between experimental data and simulations served to validate this model.

They also used their results to discredit the conjecture that replication initiation and division are coupled. Specifically, they considered the alternative hypothesis that instead cells would add a constant mass between specific events (“X-events”) of the cell cycle, such as chromosome replication initiation. This defined a “phase-shifted model”. By comparison with their experimental results, they rejected such as model and concluded that the “constant elongation model” must hold and that division is therefore not coupled to a replication initiation event. In this comparative analysis, two main points were put forward. (i) The distribution of the added size between cell birth and cell division, ∆*L*, and the distribution of the cell size at birth, *L*_b_, were aberrantly broad in simulations of the phase-shifted model. These wide fluctuations were attributed by the authors to the fact that the number of X-events per generation could fluctuate a lot. (ii) The phase-shifted model resulted in correlations between mother/daughter cells for ∆*L*, in contradiction with the absence of correlations seen in experimental data. The authors argued that this is because the added size between X-events can overlap several generations in the phase-shifted model, resulting in correlation in ∆*L*.

In this section, we will show instead that the wide fluctuations obtained result from the choice of parameters for the phase-shifted model. Actually, it will appear that the cell size convergence in the phase-shifted model critically depends on the value of the phase shift. Specifically, it can deviate significantly from the adder convergence, and even become an unstable model. We conclude this discussion by suggesting an alternative model for the cell cycle. This model relies on an adder principle holding between replication initiation events, and assume that division and replication initiation are coupled. Yet for this model, cell size convergence is consistent with adder and the simulated data would be consistent with the experimental data from Campos and colleagues [12]. Altogether, this suggests that discarding the co-regulation hypothesis between replication initiation and division based on the simulation results of the phase-shifted model is not reasonable.

#### B. Cell size convergence with the phase-shifted model

To be consistent with our manuscript we adopt notations different from Campos and colleagues. The correspondence between their and our notations are summarized in Suppl. Table T3.

##### B1) Model

We now describe the phase-shifted model proposed by Campos and colleagues [12]. First, they assumed that cells elongate their size exponentially according to Eq. (1). Second, they introduced a cellular event, denoted by the lower script *X*, which determines division timing. Specifically, provided that a cellular event occured at time *t*_X_, cell division is bound to happen at time *t*_d_ = *t*_X_ + τ_X_. Such an event does not necessarily coicinde with cell birth. Instead, it will typically represent chromosome replication initiation. Also, the cellular event triggering cell division may occur in the mother cell or other ancestors. In this model, cell division timing is therefore related to the timing of these specific cellular events, or “X-events”. Third, they proposed that an X-event occurs when a fixed size ∆_d_ has been added since the last X-event. More accurately, we introduce the quantity ∆_X_*S* which is reset to ∆_X_*S* = 0 when an X-event occurs, and otherwise increases according to:

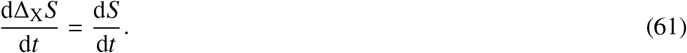

In particular, if the last X-event happened in the current generation, then:

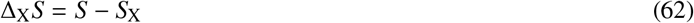

Yet if the last X-event happened in the previous generation (say with index *n* − 1), then:

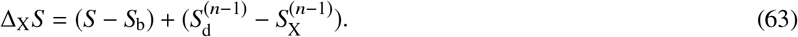

Whenever the added size since the last X-event reaches a fixed quantity, ∆_X_*S* = ∆_X_, an X-event occurs and ∆_X_*S* is reset to zero.

##### B2) Cell size convergence for small perturbations

We now investigate the convergence of cell size from a perturbed initial value in the phase shifted model. We distinguish two cases, depending on the value of *τ*_X_ compared to the generation time *τ* = ln 2/*λ*. Note that the results below are derived assuming that there is exactly one X-event per generation. In order for this assumption to hold, we restrain ourselves to small perturbations around the steady state cell size value. For larger perturbations, multiple X-events may occur in one generation, which will be dealt with numererically in the next section.

##### B3) Case 1

0 ≤ *τ*_X_ < *τ*

In this scenario, the X-event leading to cell division occurs in the same generation. As such the division size is expressed as:

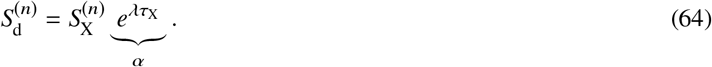

Therefore the convergence in cell size at division, is determined by the convergence of 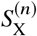. We now express the cell size at the X-event:

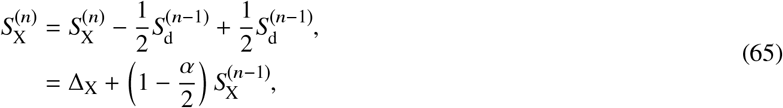

where we used that 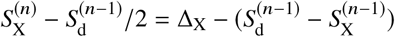. This is a first order recurrent series. We obtain:

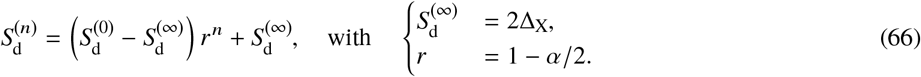

Note that Eq. (65) holds only when there is one X-event per generation. Namely, if 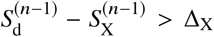 then 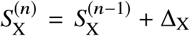 instead. In Eq. (66), we see that cell size converges exponentially to the steady state value 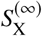. When *τ*_X_ = 0, we obtain *r* = 1/2 which is the adder convergence. Indeed, in the latter case, the phase-shifted model reduces to the adder principle [13]. However, when *τ*_X_ > 0, then *r* < 1/2 and the convergence is faster than adder.

##### B4) Case 2

τ ≤ τ_X_ < 2τ

In this scenario, the X-event leading to cell division occurs in the previous cell generation because *τ*_X_ > *τ*. As such, the division size is expressed as:

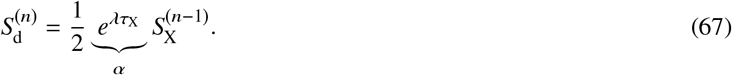

Similarly as before, we express the cell size at the X-event as:

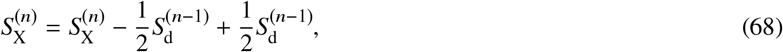

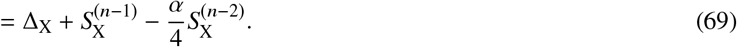

Therefore, the series 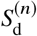 satisfies the second order recurrence relation:

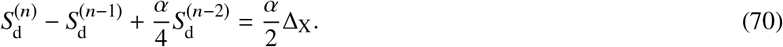

Eq. (70) is solved using standard results on series. The homogeneous solution is obtained by considering the characteristics equation:

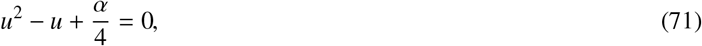

with imaginary solutions (because *α* > 1):

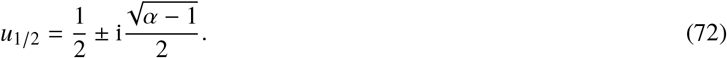

The general solution must be a linear combination of the series 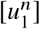 and 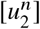. Using the particular solution 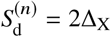 to Eq. (70) and the fact that the solution must be real, we finally find the solution:

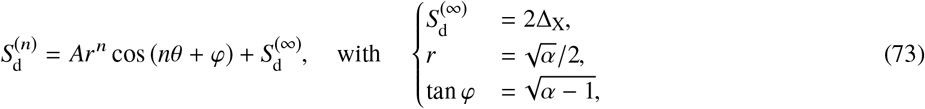

where *A* and *φ* are two constants determined by the initial condition 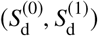. Eq. (73) defines a regime in which the convergence is slower than for adder since *r* > 1/2. In addition, the presence of oscillations in the response to perturbations to cell size suggests that in the presence of stochastic fluctuations the distribution of cell size would be quite large.

##### B5) Cell size convergence for general perturbations

As emphasized earlier, the analytical expressions Eqs. (66) and (73) are only valid for small perturbations. For larger perturbations, more than one X-event may occur per cell cycle. The actual generation time of individual cells during convergence may then vary significantly, resulting in the cell size convergence to be a combination of the scenari discussed previously. We investigated numerically cell size relaxation from a perturbed initial condition (Suppl. Fig. T13). We defined *τ* = ln 2/*λ* = 1 as unit of time and ∆_d_ = 1 as unit of size. We took the initial condition 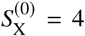. We observed cell size convergence in agreement with the analytical cases discussed above. In particular, for *τ*_X_ = 0 the cell size convergence is like adder. For *τ*_X_ = 0.5*τ* we find that the cell size converges faster than adder. Gradually as *τ*_X_ increases, the cell size convergence becomes slower than adder, and even oscillations appear.

##### B6) Comments

In summary, both analytical expressions and numerical simulations indicate that the cell size convergence in the phase-shifted model can deviate significantly from adder. This stems from the very definition of the phase-shifted model. In particular, for the values tested by Campos and colleagues [12], *τ*_X_ = 1.3*τ* and *τ*_X_ = 2.2*τ*, cell size convergence not only is slower than adder but also exhibits an oscillatory response to perturbations. As such, it is expected that the distribution of cell size in a stochastic implementation of this model will be very broad, which is one of the reasons invoked to reject the phase-shifted model and consequently refute the idea that division is controlled by chromosome replication initiation. However, this feels somehow excessive since there are other models implementing a control of division by initiation that would not lead to such an aberrant convergence property for cell size.

#### C. Alternative adder model for cell cycle based on replication initiation control

In this section, we present an alternative model for the cell cycle controlled by initiation events, yet satisfying the adder convergence for cell size and the absence of correlations for the added size. Namely, we consider that chromosome replication initiates after a fixed volume per origin of replication has been added since the last replication initiation. Since at division, the number of origins of replication is divided by two, Eqs. (65) and (69) become:

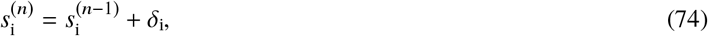

where *s*_i_ = *S*_i_/*N*_ori_ is the volume per origin of replication at initiation and δ_i_ = ∆_d_/*N*_ori_. This ensures that:

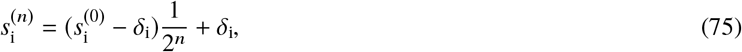

which is the adder convergence.

**Supplemental Figure T13:**
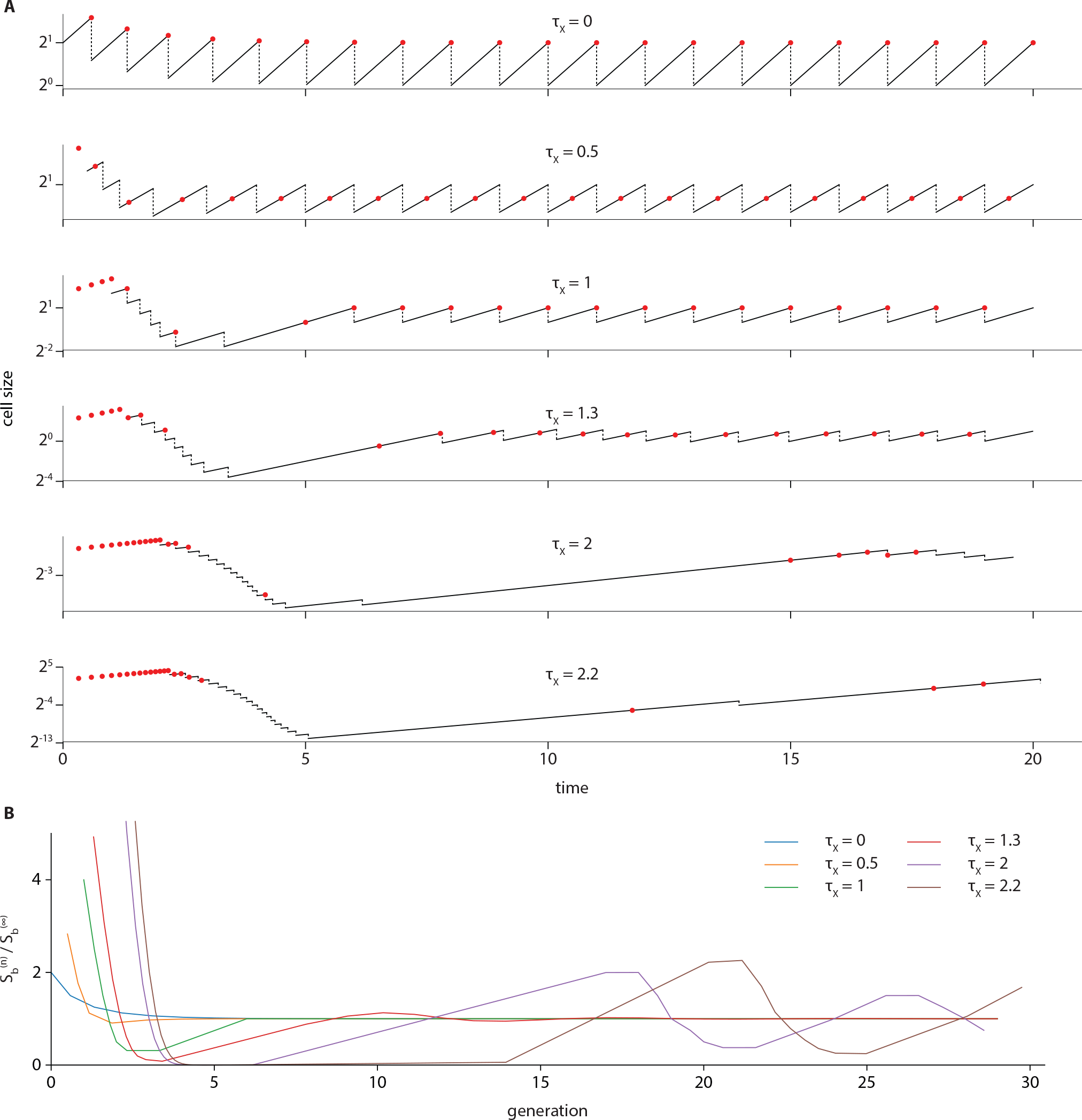
Deterministic cell size relaxation in the phase-shifted model [12]. **(A)** Simulated lineages with different values of the phase shift *τ*_X_. **(B)** Overlay of cell size convergence for the simulated lineages.

We now consider that each initiation event leads to a cell division event after a time equal to *τ*_cyc_ has elapsed, hence assuming that division is regulated by chromosome replication initiation. In general, *τ*_cyc_ may be larger than the generation time *τ* = *λ*/ln 2. Therefore, the size at cell division is given by:

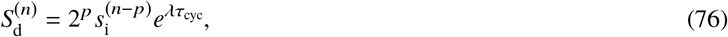

where *p* is the integer part of *τ*_*cyc*_/*τ*. Note that the number of origin of replication is *N*_ori_ = 2^*p*^. For simplicity, here we assume that *p* is fixed, meaning that the replication initiation event leading to the cell division of the current generation always occurs in the same relative ancestor (*e.g.* mother, grand-mother, *etc*.). The added size between division events is then expressed as:

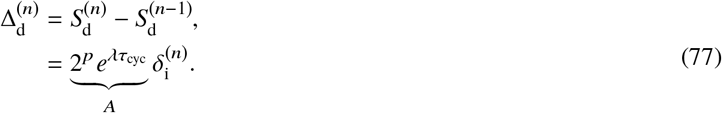

We thus obtain that the mother/daughter correlation for the division adder is related to the mother/daughter correlation for the initiation adder:

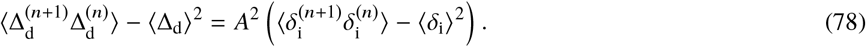

Provided that the added size per origin of replication in Eq. (74) is uncorrelated to the next, we retrieve that the added size between divisions is uncorrelated from mother to daughter cells.

## APPENDICES

## Appendix A: Properties of Gaussian bivariate distributions

### A1) Conditional probability

Let us consider two stochastic variables *X* and *Y* distributed according to a Gaussian bivariate distribution. We shall consider for simplicity that both *X* and *Y* are centered:

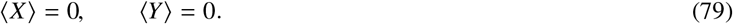

The distribution of a random Gaussian vector ***R*** = (*X*, *Y*) is characterized by the covariance matrix:

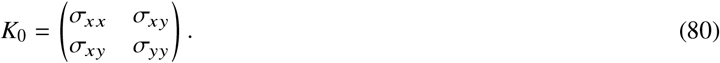

The variance of *X* (resp. *Y*) is *σ*_*xx*_ (resp. *σ*_*yy*_) and the covariance between variables *X* and *Y* is given by cov (*X*, *Y*) = *σ*_*xy*_. The Pearson correlation coefficient between variables *X* and *Y* is expressed as:

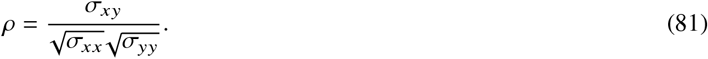

The probability distribution of the random vector ***R*** = (*X*, *Y*) is given by:

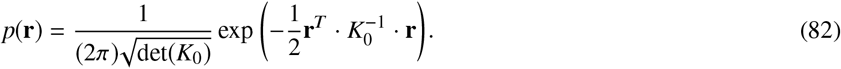

Denoting *p*(*x*, *y*) = *p*(**r**), and using the definition for conditional probabilities: *p*(*x*|*y*) = *p*(*x*, *y*)/*p*(*y*), we obtain:

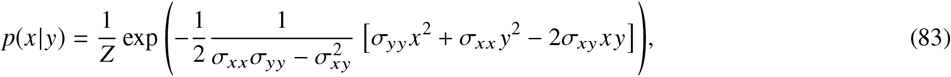

where *Z* is a normalization constant depending on *y*. This normalization is obtained by ensuring that 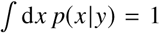. We finally obtain:

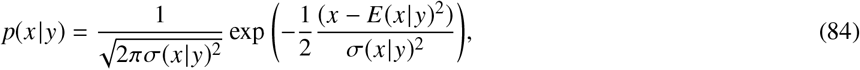

which is a Gaussian distribution with mean:

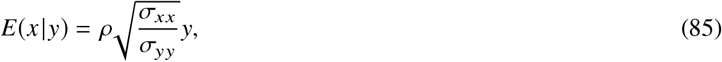

and variance:

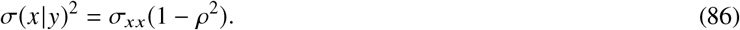

When *ρ* = 0, we find that this Gaussian distribution does not depends on *y*. When *ρ* = ±1, the variance *σ*(*x*|*y*)^2^ → 0, *i.e. x* becomes a deterministic variable, with value *y* (or −*y*) if *σ*_*xx*_ = *σ*_*yy*_.

### A2) Correlation of the inverse

We first consider the random vector *f* (***R***) = (*f* (*X*), *f* (*Y*)), where *f* is a quadratic function:

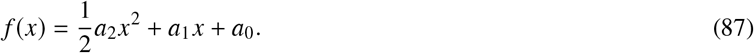

We now ask what is the Pearson correlation between variables *f* (*X*) and *f* (*Y*) given the correlation between variables *X* and *Y*. Introducing the vector Λ = (*λ, η*), one can for instance consider the characteristic function:

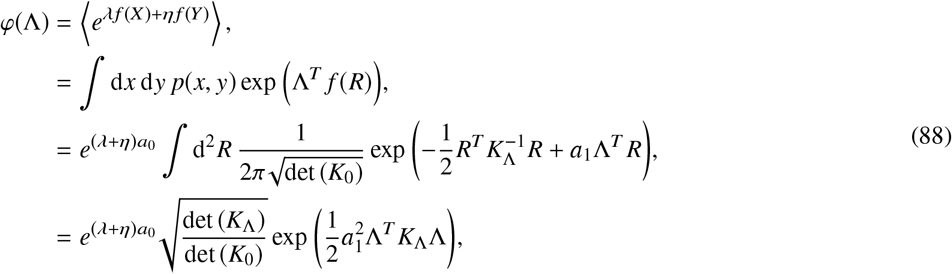

where we introduced the matrix:

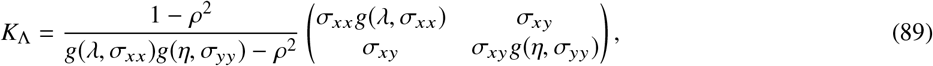

and the function *g*(*λ, σ*^2^) = 1 − *λσ*^2^*a*_2_(1 − *ρ*^2^). We then obtain the desired correlations by taking the derivative of the logarithm of the characteristic function:

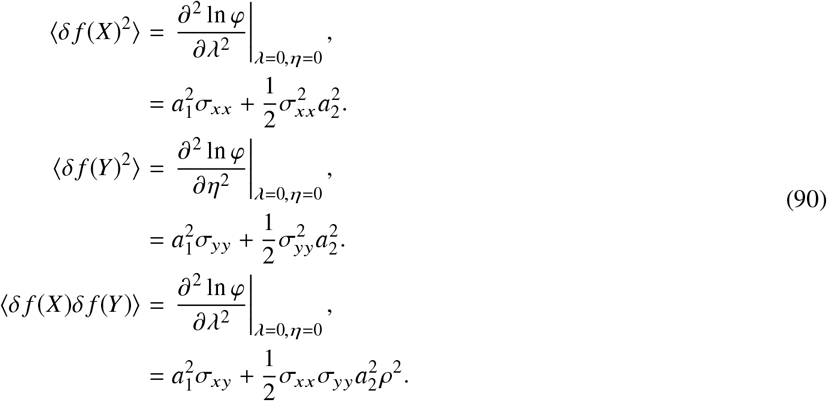

We therefore obtain for the Pearson correlation coefficient:

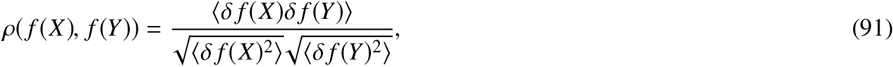

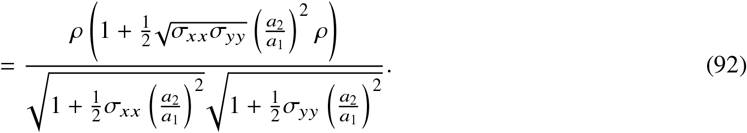

Eq. (92) is an exact result. We see that when 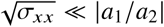 and 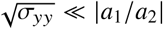, the correlation between the transformed variables is equal to the correlation between the two variables: *ρ*(*f*(*X*),*f*(*Y*)) ≈ *ρ*(*X,Y*).

Let us consider now the case where the function of interest is the inverse function:

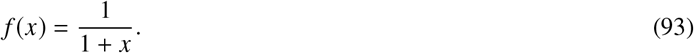

The result in Eq. (92) does not strictly hold because Eq. (93) is not a quadratic form. However, if the fluctuations in *X* are not too large, one might expect that the fluctuations of *f*(*X*) around 1 are not too large either. In this case, one might approximate *f*(*x*) to a Taylor expansion. We thus obtain a quadratic form as in Eq. (87), with *a*_2_ = 2, *a*_1_ = −1 and *a*_0_ = 1. Considering that the two variables have the same variance *σ*_*xx*_ = *σ*_*yy*_ = *σ*^2^, we show in Suppl. Fig. T14 that when *σ* ≪ |*a*_1_/*a*_2_| = 0.5, the Pearson correlation of the transformed variables is approximately equal to the Pearson correlation between the two variables: *ρ*(*f* (*X*), *f* (*Y*)) ≈ ρ(*X*, *Y*) as long as σ is not too large.

## Appendix B: Time-dependent growth rate in balanced growth

In this section, we derive the equation describing the time evolution of the instantaneous growth rate in single cells (*i.e.* the elongation rate for rod-shaped bacteria). We then generalize the equation describing concentration dynamics, namely Eq. (39).

Let us denote *M* the dry mass of an individual cell. We assume that the total mass increase is directly proportional to the number of ribosomes in the cytoplasm. Thus we have:

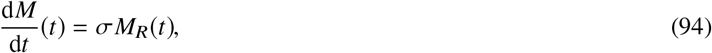

where σ corresponds to the amount of new biomass produced per ribosome and *M*_*R*_ is the mass of ribosomes in the cell. Note that we assume that σ is constant, that is to say that the translation load of ribosomes is invariant through time. Furthermore, in balanced growth, a fixed fraction of the mass increase is allocated to ribosome synthesis:

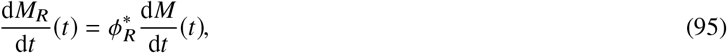

where 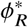 is the fixed fraction of the mass flux allocated to ribosome synthesis. Introducing the instantaneous mass fraction of ribosomes *φ*_*R*_(*t*) = *M*_*R*_(*t*)/*M*(*t*), we obtain from Eq. (94) the equation for exponential growth:

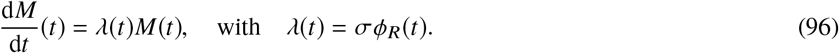

**Supplemental Figure T14:**
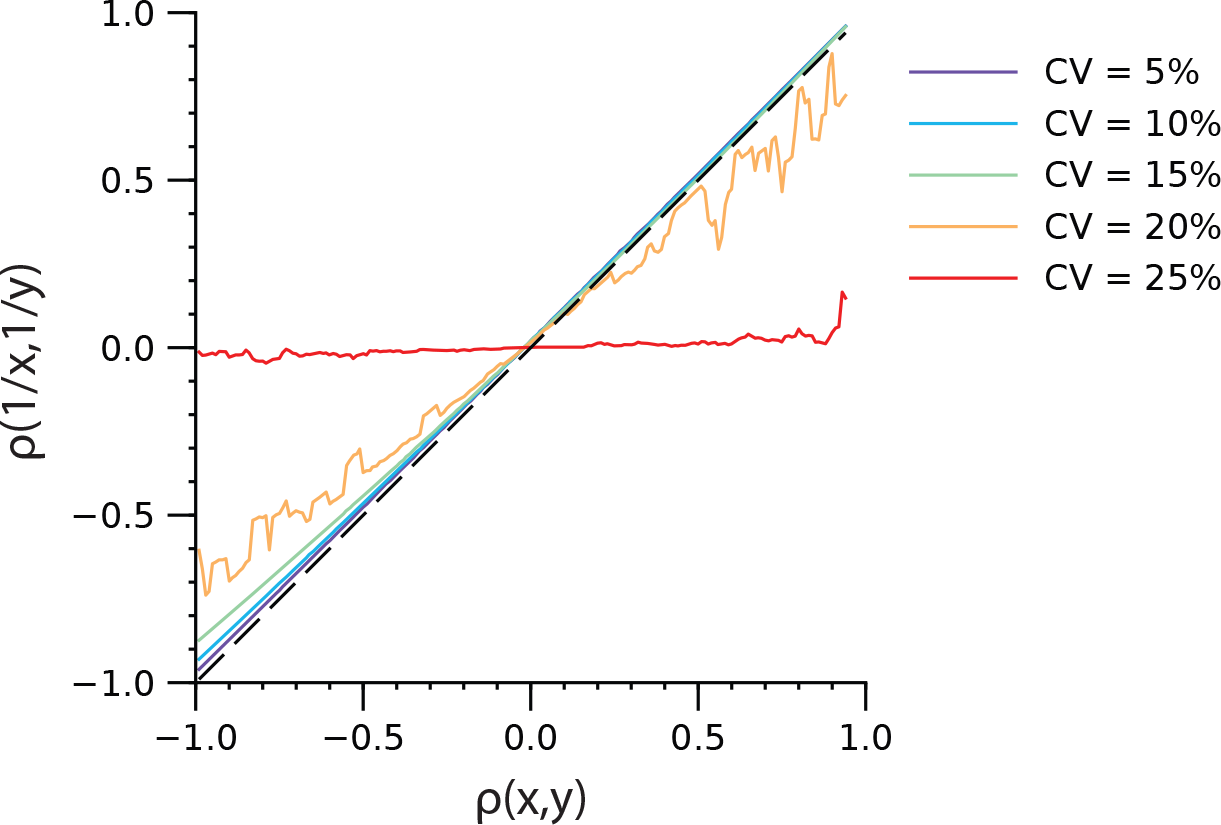
Pearson correlation between *f*(*X*) and *f*(*Y*) when *f*(*x*) is the function in Eq.(93). (*X,Y*) is a random vector distributed according to a Gaussian bivariate distribution, with means *〈X〉* = *〈Y〉* = 1, variances *σ*_*xx*_ = *σ*_*yy*_ = *σ*^2^ and covariance *σ*_*xy*_ = *ρ*(*x,y*)*σ*. In this case, the coefficient-of-variation is CV = *σ*.

Using Eqs. (94) and (95), we obtain the equation describing the time-evolution of the instantaneous growth rate:

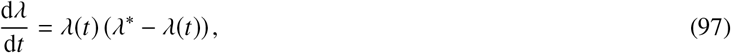

where 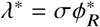 is the steady-state growth rate.

Let us now consider a generic protein with instantaneous mass *M*_*P*_ in the cell. Again we assume that this protein is produced under balanced growth:

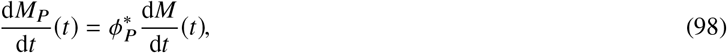

where 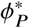 is the fixed fraction of the mass flux allocated to the biosynthesis of protein *P*. Using Eq. (96), we find that the instantaneous mass fraction *φ*_*P*_(*t*) = *M*_*P*_(*t*)/*M*(*t*) satisfies:

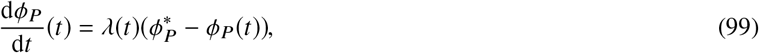

or in terms of concentrations:

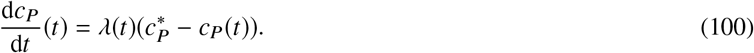

## REFERENCES

1. Kiviet, D. J., Nghe, P., Walker, N., Boulineau, S., Sunderlikova, V., and Tans, S. J. (2014). Stochasticity of metabolism and growth at the single-cell level. Nature 514, 376–9.

2. Campos, M., Surovtsev, I. V., Kato, S., Paintdakhi, A., Beltran, B., Ebmeier, S. E., and Jacobs-Wagner, C. (2014). A constant size extension drives bacterial cell size homeostasis. Cell 159, 1433–1446.

3. Taheri-Araghi, S., Bradde, S., Sauls, J. T., Hill, N. S., Levin, P. A., Paulsson, J., Vergassola, M., and Jun, S. (2015). Cell-size control and homeostasis in bacteria. Curr. Biol. 25, 385–391.

4. Di Talia, S., Skotheim, J. M., Bean, J. M., Siggia, E. D., and Cross, F. R. (2007). The effects of molecular noise and size control on variability in the budding yeast cell cycle. Nature 448, 947–951.

5. Santi, I., Dhar, N., Bousbaine, D., Wakamoto, Y., and Mckinney, J. D. (2013). Single-cell dynamics of the chromosome replication and cell division cycles in mycobacteria. Nat. Commun. 4, 2470.

6. Iyer-Biswas, S., Wright, C. S., Henry, J. T., Lo, K., Burov, S., Lin, Y., Crooks, G. E., Crosson, S., Dinner, A. R., and Scherer, N. F. (2014). Scaling laws governing stochastic growth and division of single bacterial cells. Proc. Natl. Acad. Sci. U.S.A 111, 15912–15917.

7. Deforet, M., Van Ditmarsch, D., and Xavier, J. B. (2015). Cell-size homeostasis and the incremental rule in a bacterial pathogen. Biophys. J. 109, 521–528.

8. Logsdon, M. M., Ho, P.-Y., Papavinasasundaram, K., Richardson, K., Cokol, M., Sassetti, C. M., Amir, A., and Aldridge, B. B. (2017). A parallel adder coordinates mycobacterial cell-cycle progression and cell-size homeostasis in the context of asymmetric growth and organization. Curr. Biol. 27, 3367–3374.e7.

9. Soifer, I., Robert, L., and Amir, A. (2016). Single-cell analysis of growth in budding yeast and bacteria reveals a common size regulation strategy. Curr. Biol. 26, 356–361.

10. Chandler-Brown, D., Schmoller, K. M., Winetraub, Y., and Skotheim, J. M. (2017). The adder phenomenon emerges from independent control of pre- and post-start phases of the budding yeast cell cycle. Curr. Biol. 27, 2774–2783.

11. Varsano, G., Wang, Y., and Wu, M. (2017). Probing mammalian cell size homeostasis by channel-assisted cell reshaping. Cell Rep. 20, 397–410.

12. Cadart, C., Monnier, S., Grilli, J., Sáez, P. J., Srivastava, N., Attia, R., Terriac, E., Baum, B., Cosentino-Lagomarsino, M., and Piel, M. (2018). Size control in mammalian cells involves modulation of both growth rate and cell cycle duration. Nat. Commun. 9, 3275.

13. Jun, S. and Taheri-Araghi, S. (2015). Cell-size maintenance: universal strategy revealed. Trends Microbiol. 23, 4–6.

14. Sauls, J. T., Li, D., and Jun, S. (2016). Adder and a coarse-grained approach to cell size homeostasis in bacteria. Curr Opin Cell Biol. 38, 38–44.

15. Jun, S., Si, F., Pugatch, R., and Scott, M. (2018). Fundamental principles in bacterial physiology - history, recent progress, and the future with focus on cell size control: A review. Rep. Prog. Phys. 81, 056601.

16. Willis, L. and Huang, K. C. (2017). Sizing up the bacterial cell cycle. Nat. Rev. Microbiol. 15, 606–620.

17. Amir, A. (2017). Is cell size a spandrel? eLife 6, e22186.

18. Donachie, W. D. (1968). Relationship between Cell Size and Time of Initiation of DNA Replication. Nature 219, 1077–1079.

19. Wallden, M., Fange, D., Lundius, E. G., Baltekin, ö., and Elf, J. (2016). The synchronization of replication and division cycles in individual *E. coli* cells. Cell 166, 729–739.

20. Si, F., Li, D., Cox, S. E., Sauls, J. T., Azizi, O., Sou, C., Schwartz, A. B., Erickstad, M. J., Jun, Y., Li, X., and Jun, S. (2017). Invariance of initiation mass and predictability of cell size in *Escherichia coli*. Curr. Biol., 27, 1278–1287.

21. Basan, M., Zhu, M., Dai, X., Warren, M., Sévin, D., Wang, Y. P., and Hwa, T. (2015). Inflating bacterial cells by increased protein synthesis. Mol. Syst. Biol. 11, 836.

22. Bertaux, F., Kügelgen, J. V., Marguerat, S., and Shahrezaei, V. (2016). A unified coarse-grained theory of bacterial physiology explains the relationship between cell size, growth rate and proteome composition under various growth limitations. bioRxiv doi: https://doi.org/10.1101/078998.

23. Osella, M., Nugent, E., and Lagomarsino, M. C. (2014). Concerted control of *Escherichia coli* cell division. Proc. Natl. Acad. Sci. U.S.A 111, 3431–3435.

24. Micali, G., Grilli, J., Marchi, J., Osella, M., and Lagomarsino, M. C. (2018a). Dissecting the control mechanisms for DNA replication and cell division in *E. coli*. Cell Rep. 25, 761–771.e4.

25. Harris, L. K. and Theriot, J. A. (2016). Relative rates of surface and volume synthesis set bacterial cell size. Cell 165, 1479–1492.

26. Wang, P., Robert, L., Pelletier, J., Dang, W. L., Taddei, F., Wright, A., and Jun, S. (2010). Robust growth of *Escherichia coli*. Curr. Biol. 20, 1099–1103.

27. Adiciptaningrum, A., Osella, M., Moolman, M. C., Lagomarsino, M. C., and Tans, S. J. (2015). Stochasticity and homeostasis in the *E. coli* replication and division cycle. Sci. Rep. 5, 18261.

28. Mangiameli, S. M., Veit, B. T., Merrikh, H., and Wiggins, P. A. (2017). The replisomes remain spatially proximal throughout the cell cycle in bacteria. PLoS Genet. 13, e1006582.

29. Holden, S. J., Uphoff, S., and Kapanidis, A. N. (2011). Daostorm: an algorithm for high-density super-resolution microscopy. Nat. Methods 8, 279–280.

30. Cox, S., Rosten, E., Monypenny, J., Jovanovic-Talisman, T., Burnette, D. T., Lippincott-Schwartz, J., Jones, G. E., and Heintzmann, R. (2011). Bayesian localization microscopy reveals nanoscale podosome dynamics. Nat. Methods, 9, 195–200.

31. Sompayrac, L. and Maaløe, O. (1973). Autorepressor model for control of DNA replication. Nature 241, 133–135.

32. Li, X.-T., Jun, Y., Erickstad, M. J., Brown, S. D., Parks, A., Court, D. L., and Jun, S. (2016). tCRISPRi: tunable and reversible, one-step control of gene expression. Sci. Rep. 6, 39076.

33. Schaechter, M., Maaløe, O., and Kjeldgaard, N. O. (1958). Dependency on medium and temperature of cell size and chemical composition during balanced growth of *Salmonella typhimurium*. J. Gen. Microbiol. 19, 592–606.

34. Koch, A. L. (1977). Does the initiation of chromosome replication regulate cell division? In Advances in Microbial Physiology, A.H. Rose and D.W. Tempest ed. (Elsevier), pp 49–98.

35. Helmstetter, C. E. and Cooper, S. (1968). DNA synthesis during the division cycle of rapidly growing *Escherichia coli* B/r. J. Mol. Biol. 31, 507–518.

36. Cooper, S. and Helmstetter, C. E. (1968). Chromosome replication and the division cycle of *Escherichia coli* B/r. J. Mol. Biol., 31, 519–540.

37. Amir, A. (2014). Cell size regulation in bacteria. Phys. Rev. Lett. 112, 208102.

38. Susman, L., Kohram, M., Vashistha, H., Nechleba, J. T., Salman, H., and Brenner, N. (2018). Individuality and slow dynamics in bacterial growth homeostasis. Proc. Natl. Acad. Sci. U.S.A 115, E5679–E5687.

39. Hansen, F. G., Atlung, T., Braun, R. E., Wright, A., Hughes, P., and Kohiyama, M. (1991). Initiator (DnaA) protein concentration as a function of growth rate in *Escherichia coli* and *Salmonella typhimurium*. J. Bacteriol. 173, 5194–9.

40. Skarstad, K. and Katayama, T. (2013). Regulating DNA replication in bacteria. Cold Spring Harb. Perspect. Biol. 5, a012922.

41. Hansen, F. G. and Atlung, T. (2018). The DnaA tale. Front. Microbiol. 9, 319.

42. Løbner-Olesen, A., Skarstad, K., Hansen, F. G., Meyenburg, K. V., and Boye, E. (1989). The DnaA protein determines the initiation mass of *Escherichia coli* K-12. Cell 57, 881–889.

43. Riber, L., Olsson, J. A., Jensen, R. B., Skovgaard, O., Dasgupta, S., Marinus, M. G., and Løbner-Olesen, A. (2006). Hda-mediated inactivation of the DnaA protein and DnaA gene autoregulation act in concert to ensure homeostatic maintenance of the *Escherichia coli* chromosome. Genes Dev. 20, 2121–2134.

44. Donachie, W. D., and Begg, M. K. J. (1971). Independence of Cell Division and DNA Replication in *Bacillus subtilis*. Nature 231, 274–276.

45. Micali, G., Grilli, J., Osella, M., and Lagomarsino, M. C. (2018b). Concurrent processes set *E. coli* cell division. Sci. Adv. 4, eaau3324.

46. Harry, E., Monahan, L., and Thompson, L. (2006). Bacterial cell division: the mechanism and its precision. Int. Rev. Cytol. 253, 27–94.

47. Haeusser, D. P. and Margolin, W. (2016). Splitsville: structural and functional insights into the dynamic bacterial Z ring. Nat. Rev. Microbiol. 14, nrmicro.2016.26.

48. Palacios, P., Vicente, M., and Sánchez, M. (1996). Dependency of *Escherichia coli* cell-division size, and independency of nucleoid segregation on the mode and level of FtsZ expression. Mol. Microbiol., 20, 1093–1098.

49. Zheng, H., Ho, P.-Y., Jiang, M., Tang, B., Liu, W., Li, D., Yu, X., Kleckner, N. E., Amir, A., and Liu, C. (2016). Interrogating the *Escherichia coli* cell cycle by cell dimension perturbations. Proc. Natl. Acad. Sci. U.S.A. 113, 15000–15005.

50. Chien, A.-C., Hill, N. S., and Levin, P. A. (2012). Cell size control in bacteria. Curr. Biol. 22, R340–9.

51. Garrido, T., Sánchez, M., Palacios, P., Aldea, M., and Vicente, M. (1993). Transcription of FtsZ oscillates during the cell cycle of *Escherichia coli*. EMBO J. 12, 3957–65.

52. Jameson, K. H. and Wilkinson, A. J. (2017). Control of initiation of DNA replication in *Bacillus subtilis* and *Escherichia coli*. Genes 8, 22.

53. Goranov, A. I., Breier, A. M., Merrikh, H., and Grossman, A. D. (2009). YabA of *Bacillus subtilis* controls DnaA-mediated replication initiation but not the transcriptional response to replication stress. Mol. Microbiol. 74, 454–466.

54. Moore, D. A., Whatley, Z. N., Joshi, C. P., Osawa, M., and Erickson, H. P. (2017). Probing for binding regions of the FtsZ protein surface through site-directed insertions: Discovery of fully functional FtsZ-fluorescent proteins. J. Bacteriol. 199, e00553–16.

55. Sigal, A., Milo, R., Cohen, A., Geva-Zatorsky, N., Klein, Y., Liron, Y., Rosenfeld, N., Danon, T., Perzov, N., and Alon, U. (2006). Variability and memory of protein levels in human cells. Nature 444, 643–646.

56. Jajoo, R., Jung, Y., Huh, D., Viana, M. P., Rafelski, S. M., Springer, M., and Paulsson, J. (2016). Accurate concentration control of mitochondria and nucleoids. Science 351, 169–172.

57. Teather, R. M., Collins, J. F., and Donachie, W. D. (1974). Quantal behavior of a diffusible factor which initiates septum formation at potential division sites in *Escherichia coli*. J. Bacteriol. 118, 407–13.

58. Bi, E. and Lutkenhaus, J. (1990). FtsZ regulates frequency of cell division in *Escherichia coli*. J. Bacteriol. 172, 2765–8.

59. Ghusinga, K. R., Vargas-Garcia, C. A., and Singh, A. (2016). A mechanistic stochastic framework for regulating bacterial cell division. Sci. Rep. 6, 30229.

60. Aarsman, M. E. G., Piette, A., Fraipont, C., Vinkenvleugel, T. M. F., Nguyen-Distèche, M., and Blaauwen, T. D. (2005). Maturation of the *Escherichia coli* divisome occurs in two steps. Mol. Microbiol., 55, 1631–1645.

61. van der Ploeg, R., Verheul, J., Vischer, N. O. E., Alexeeva, S., Hoogendoorn, E., Postma, M., Banzhaf, M., Vollmer, W., and Blaauwen, T. (2013). Colocalization and interaction between elongasome and divisome during a preparative cell division phase in *Escherichia coli*. Mol. Microbiol. 87, 1074–1087.

62. Söderström, B., Skoog, K., Blom, H., Weiss, D. S., Heijne, G., and Daley, D. O. (2014). Disassembly of the divisome in *Escherichia coli*: evidence that FtsZ dissociates before compartmentalization. Mol. Microbiol. 92, 1–9.

63. Coltharp, C., Buss, J., Plumer, T. M., and Xiao, J. (2016). Defining the rate-limiting processes of bacterial cytokinesis. Proc. Natl. Acad. Sci. U.S.A 113, E1044–E1053.

64. Sekar, K., Rusconi, R., Sauls, J. T., Fuhrer, T., Noor, E., Nguyen, J., Fernandez, V. I., Buffing, M. F., Berney, M., Jun, S., Stocker, R., and Sauer, U. (2018). Synthesis and degradation of FtsZ quantitatively predict the first cell division in starved bacteria. Mol. Syst. Biol. 14, e8623.

65. Männik, J., Walker, B. E., and Männik, J. (2018). Cell cycle-dependent regulation of FtsZ in *Escherichia coli* in slow growth conditions. Mol. Microbiol. 110, 1030–1044

66. Fantes, P. A., Grant, W. D., Pritchard, R. H., Sudbery, P. E., and Wheals, A. E. (1975). The regulation of cell size and the control of mitosis. J. Theor. Biol. 50, 213–244.

67. Ho, P.-Y. and Amir, A. (2015). Simultaneous regulation of cell size and chromosome replication in bacteria. Front. Microbiol. 6, 662.

68. Tanouchi, Y., Pai, A., Park, H., Huang, S., Stamatov, R., Buchler, N. E., and You, L. (2015). A noisy linear map underlies oscillations in cell size and gene expression in bacteria. Nature 523, 357–360.

69. E. Lyons, M. Freeling, S. Kustu, and W. Inwood. (2011). Using genomic sequencing for classical genetics in *E. coli* k12. PLoS ONE 6, e16717.

70. S. D. Brown and S. Jun. (2015). Complete genome sequence of *Escherichia coli* ncm3722. Genome Announc. 3, e00879–15.

71. R. Reyes-Lamothe, D. J. Sherratt, and M. C. Leake. (2010). Stoichiometry and architecture of active DNA replication machinery in *Escherichia coli*. Science, 328, 498–501.

72. S. Nishida, K. Fujimitsu, K. Sekimizu, T. Ohmura, T. Ueda, and T. Katayama. (2002). A nucleotide switch in the *Escherichia coli* DnaA protein initiates chromosomal replication evidence from a mutant DnaA protein defective in regulatory ATP hydrolysis in vitro and in vivo. J. Biol. Chem. 277, 14986–14995.

## SUPPLEMENTAL REFERENCES

[1] F. Si, D. Li, S. E. Cox, J. T. Sauls, O. Azizi, C. Sou, A. B. Schwartz, M. J. Erickstad, Y. Jun, X. Li, and S. Jun, Invariance of initiation mass and predictability of cell size in *Escherichia coli*. Current Biology, 27, 1278–1287. (2017).

[2] E. Lyons, M. Freeling, S. Kustu, and W. Inwood, Using genomic sequencing for classical genetics in *E. coli* k12. PLoS ONE, 6, e16717. (2011).

[3] S. D. Brown and S. Jun, Complete genome sequence of *Escherichia coli* ncm3722. Genome Announcements, 3, e00879–15. (2015).

[4] X.-T. Li, Y. Jun, M. J. Erickstad, S. D. Brown, A. Parks, D. L. Court, and S. Jun, tCRISPRi: tunable and reversible, one-step control of gene expression. Scientific Reports, 6, 39076. (2016).

[5] T. Garrido, M. Sánchez, P. Palacios, M. Aldea, and M. Vicente, Transcription of FtsZ oscillates during the cell cycle of *Escherichia coli*. The EMBO journal, 12, 3957–65. (1993).

[6] R. Reyes-Lamothe, D. J. Sherratt, and M. C. Leake, Stoichiometry and architecture of active DNA replication machinery in *Escherichia coli*. Science, 328, 498–501. (2010).

[7] A. Amir, F. Babaeipour, D. B. Mcintosh, D. R. Nelson, and S. Jun, Bending forces plastically deform growing bacterial cell walls. Proceedings of the National Academy of Sciences, 111, 5778–5783. (2014).

[8] S. Nishida, K. Fujimitsu, K. Sekimizu, T. Ohmura, T. Ueda, and T. Katayama, A nucleotide switch in the *Escherichia coli* DnaA protein initiates chromosomal replication evidence from a mutant DnaA protein defective in regulatory ATP hydrolysis in vitro and in vivo. Journal of Biological Chemistry, 277, 14986–14995. (2002).

[9] L. Riber, J. A. Olsson, R. B. Jensen, O. Skovgaard, S. Dasgupta, M. G. Marinus, and A. Løbner-Olesen, Hda-mediated inactivation of the DnaA protein and DnaA gene autoregulation act in concert to ensure homeostatic maintenance of the *Escherichia coli* chromosome. Genes & Development, 20, 2121–2134. (2006).

[10] A. I. Goranov, A. M. Breier, H. Merrikh, and A. D. Grossman, YabA of *Bacillus subtilis* controls DnaA-mediated replication initiation but not the transcriptional response to replication stress. Molecular Microbiology, 74, 454–466. (2009).

[11] S. M. Mangiameli, B. T. Veit, H. Merrikh, and P. A. Wiggins, The replisomes remain spatially proximal throughout the cell cycle in bacteria. PLOS Genetics, 13, e1006582. (2017).

[12] M. Campos, I. Surovtsev, S. Kato, A. Paintdakhi, B. Beltran, S. Ebmeier, and C. Jacobs-Wagner, A constant size extension drives bacterial cell size homeostasis. Cell, 159, 1433–1446. (2014).

[13] S. Taheri-Araghi, S. Bradde, J. Sauls, N. Hill, P. Levin, J. Paulsson, M. Vergassola, and S. Jun, Cell-size control and homeostasis in bacteria. Current Biology, 25, 385–391. (2015).

[14] M. Wallden, D. Fange, E. Lundius, Ö. Baltekin, and J. Elf, The synchronization of replication and division cycles in individual *E. coli* cells. Cell, 166, 729–739. (2016).

[15] S. Cooper and C. E. Helmstetter, Chromosome replication and the division cycle of *Escherichia coli* B/r. Journal of Molecular Biology, 31, 519–540. (1968).

[16] W. D. Donachie, Relationship between cell size and time of initiation of DNA replication. Nature, 219, 1077–1079. (1968).

[17] M. L. Mott and J. M. Berger, DNA replication initiation: mechanisms and regulation in bacteria. Nature Reviews Microbiology, 5, 343–354. (2007).

[18] T. Katayama, S. Ozaki, K. Keyamura, and K. Fujimitsu, Regulation of the replication cycle: conserved and diverse regulatory systems for DnaA and oric. Nature Reviews Microbiology, 8, 163–170. (2010).

[19] K. Skarstad and T. Katayama, Regulating DNA replication in bacteria. Cold Spring Harbor Perspectives in Biology, 5, a012922. (2013).

[20] K. Sekimizu, B. Y. Yung, and A. Kornberg, The dnaa protein of *Escherichia coli*. abundance, improved purification, and membrane binding. The Journal of biological chemistry, 263, 7136–40. (1988).

[21] W. Messer, The bacterial replication initiator DnaA. DnaA and oric, the bacterial mode to initiate DNA replication. FEMS Microbiology Reviews, 26, 355–374. (2002).

[22] L. Sompayrac and O. Maaløe, Autorepressor model for control of DNA replication. Nature, 241, 133–135. (1973).

[23] A.-C. Chien, N. S. Hill, and P. A. Levin, Cell size control in bacteria. Current Biology, 22, R340–R349. (2012).

[24] D. A. Moore, Z. N. Whatley, C. P. Joshi, M. Osawa, and H. P. Erickson, Probing for binding regions of the FtsZ protein surface through site-directed insertions: Discovery of fully functional FtsZ-fluorescent proteins. Journal of Bacteriology, 199, e00553–16. (2017).

[25] M. Scott, C. W. Gunderson, E. M. Mateescu, Z. Zhang, and T. Hwa, Interdependence of cell growth and gene expression: Origins and consequences. Science, 330, 1099–1102. (2010).

[26] M. Scott and T. Hwa, Bacterial growth laws and their applications. Current Opinion in Biotechnology, 22, 559–565. (2011).

